# Opponent control of behavior by dorsomedial striatal pathways depends on task demands and internal state

**DOI:** 10.1101/2021.07.23.453573

**Authors:** Scott S. Bolkan, Iris R. Stone, Lucas Pinto, Zoe C. Ashwood, Jorge M. Iravedra Garcia, Alison L. Herman, Priyanka Singh, Akhil Bandi, Julia Cox, Christopher A. Zimmerman, Jounhong Ryan Cho, Ben Engelhard, Jonathan W. Pillow, Ilana B. Witten

## Abstract

A classic view of the striatum holds that activity in direct and indirect pathways oppositely modulates motor output. Whether this involves direct control of movement, or reflects a cognitive process underlying movement, remains unresolved. Here we find that strong, opponent control of behavior by the two pathways of the dorsomedial striatum (DMS) depends on the cognitive requirements of a task. Furthermore, a latent state model (a hidden markov model with generalized linear model observations) reveals that—even within a single task—the contribution of the two pathways to behavior is state-dependent. Specifically, the two pathways have large contributions in one of two states associated with a strategy of evidence accumulation, compared to a state associated with a strategy of repeating previous choices. Thus, both the demands imposed by a task, as well as the internal state of mice when performing a task, determine whether DMS pathways provide strong and opponent control of behavior.

## INTRODUCTION

The striatum is composed of two principal outputs, the direct and indirect pathways, which are thought to exert opposing effects on behavior^1^. In support of this view, many influential studies have shown that pathway-specific activation of the striatum produces opposing behavioral biases^2–14^. For example, direct or indirect pathway activation oppositely influences locomotion^2–4,14^, licking^5,11,15^, left/right rotations^2,3,11,16^, repetition/cessation of activation-paired behaviors^6–8^, and left/right movements to report value-based decisions^9,13^.

Despite this pioneering work, it remains unresolved whether the endogenous activity of the two pathways provides opposing control over the generation of movements, or instead contributes to the cognitive process of deciding which movement to perform. This is in part because pathway-specific manipulations have disproportionately relied on artificial and synchronous activation, rather than inhibition of endogenous activity^2–11,13^. The imbalance towards reports of activation suggests a wealth of negative results from inhibition, raising questions about the function of the endogenous activity, and whether it contributes to cognition. In fact, most previous pathway-specific activation studies have not used cognitively demanding tasks, making it difficult to dissociate a role in the decision towards a movement versus the generation of the movement itself^2–4,6,11,14,17^. In contrast, studies of the striatum that were not pathway-specific have instead focused on cognitively demanding behaviors^18–25^. Taken together, this raises the possibility that striatal pathways exert opposing control of movement in the context of decision-making, rather than directly controlling motor output irrespective of cognition.

Thus, to determine if the contribution of endogenous activity in striatal pathways depends on cognition, we examined the effects of pathway-specific inhibition across a set of virtual reality tasks that had the same motor output and similar sensory features, but different cognitive requirements. This allowed us to ask if a task’s demands determined the effect of pathway-specific inhibition on behavior. Second, we used a latent state model to identify time-varying states within the same task. This allowed us to determine if the contribution of each pathway to behavior changed across time, even within the same task.

We found that inhibition of neither pathway produced a detectable influence on behavior as mice navigated a virtual corridor in the absence of a decision-making requirement. In contrast, pathway-specific inhibition produced strong and opposing biases on decisions based on the accumulation of evidence in a virtual T-maze^26^, and had weaker effects on choice during less demanding task variants. Our latent state model further revealed that even within the evidence accumulation task, mice occupy different states across time that differ in the weighting of sensory evidence and trial history, as well as the extent that pathway-specific inhibition impacts choice. Thus, by comparing the effects of pathway-specific inhibition across behavioral tasks, and across time within a task, we conclude that both demands of the task and internal state of the mice determine whether striatal pathways exert strong and opposing control over behavior.

## RESULTS

### Inhibition of pathway-specific DMS activity is effective

We first sought to validate the effectiveness of halorhodopsin^19^ (NpHR)-mediated inhibition of indirect and direct striatal pathway activity in awake, head-fixed mice (**Fig. 1a** and **Extended Data Fig. 1a-b**). Toward this end, we bilaterally delivered virus carrying Cre-dependent NpHR to the dorsomedial striatum (DMS) in transgenic mouse lines (A2a-Cre/D2R-Cre/D1R-Cre), which we verified to have high degrees of specificity and pentrance for each pathway (**Supplementary Fig. 1**). We confirmed that 532-nm (5mW) light delivery to the DMS through a tapered optical fiber produced rapid, sustained, and reversible inhibition of spiking in mice expressing NpHR in the indirect (**Fig. 1b** and **Extended Data Fig. 1c-e**, n = 18/60, 30% of neurons significantly inhibited) or direct pathway (**Fig. 1c** and **Extended Data Fig. 1f-h**, n = 21/50, 42% of neurons significantly inhibited). Moreover, we observed: (1) minimal excitation during illumination^27,28^ (**Extended Data Fig. 1d,g**, *left*), (2) minimal effects on spiking upon laser offset (**Extended Data Fig. 1d,g**, *right*), indicating limited post-inhibitory rebound, and (3) stability in the efficacy of inhibition across time (**Supplementary Fig. 2**). All together, our findings indicate that NpHR-mediated inhibition of DMS pathways is effective.

**Figure 1.**
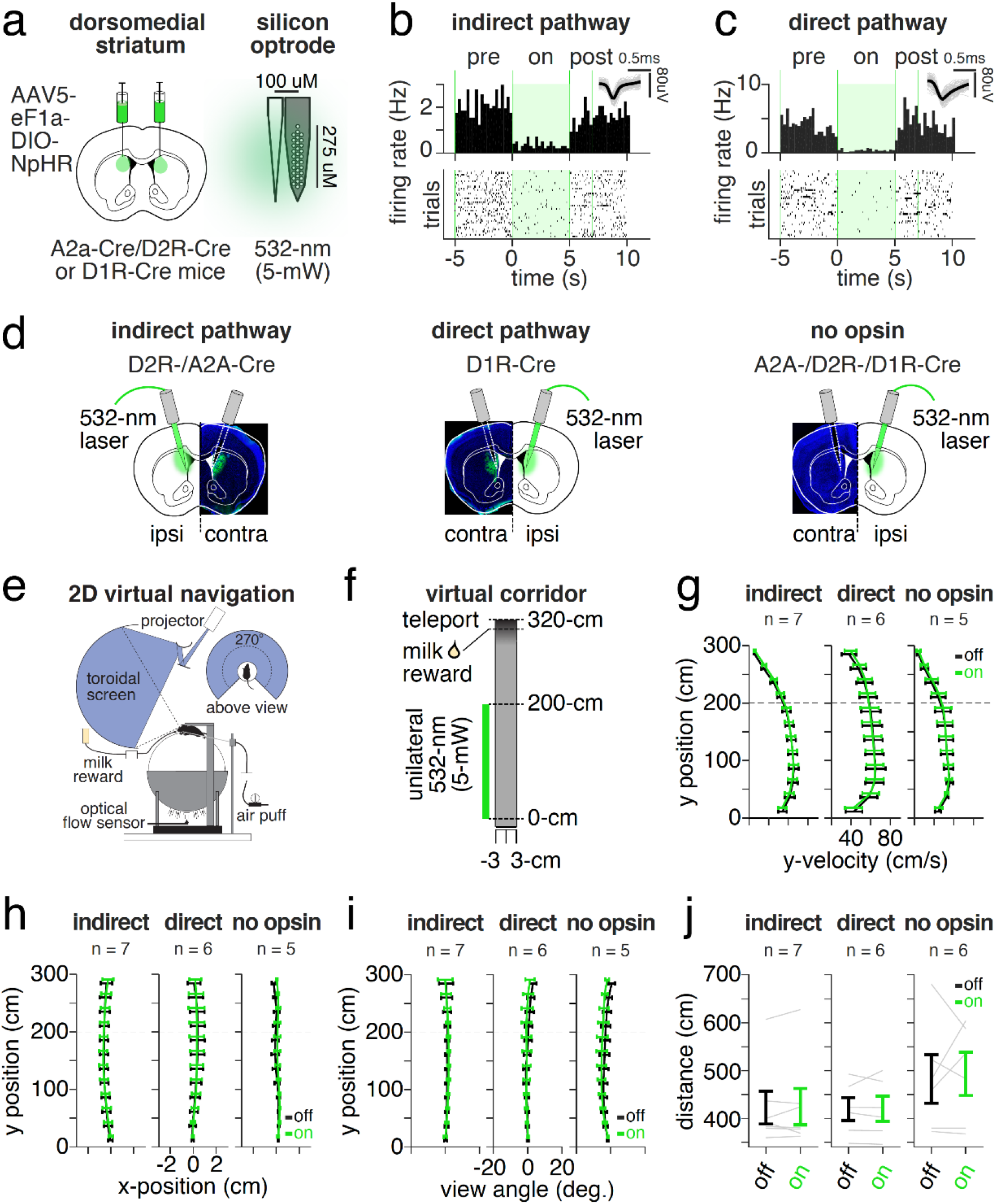
Pathway-specific DMS inhibition has no detectable impact on movement in mice navigating a virtual corridor. (**a**) Schematic of viral delivery of Cre-dependent halorhodopsin (NpHR) to the dorsomedial striatum (DMS) of A2a-Cre, D2R-Cre, or D1R-Cre mice (*left*). Schematic of optrode (*right*): a 32-channel silicon probe coupled with tapered optical fiber, which delivered 532-nm (5-mW) light to the DMS of awake, ambulating mice. (**b**) Example peristimulus time histograms (PSTH) (*top*) and rasters of trial-by-trial spike times (*bottom*) from a DMS single-unit recorded in an ambulating A2a-Cre mouse expressing Cre-dependent NpHR (indirect pathway). *Inset*: average spike waveform (black) and 100 randomly sampled spike waveforms (grey). A trial consisted of 5-s without laser (*pre*, −5 to 0-s), 5-s laser sweep (*on*, 0 to 5-s), and 10-s ITI (40 total trials). (**c**) As in **b** but for DMS single-unit in a D1R-Cre mouse expressing Cre-dependent NpHR (direct pathway). (**d**) Schematic of bilateral fiberoptic implantation of DMS and unilateral illumination in behaving mice, with example histology from a mouse expressing NpHR in the indirect (*left*, D2R-/A2a-Cre) or direct (*middle*, D1R-Cre) pathways, or control mouse without opsin (*right*, no opsin, A2a-/D2R- or D1R-Cre). 532-nm light (5-mW) was delivered unilaterally to the left or right hemisphere on alternate testing sessions and lateralized behavior was defined as ipsilateral or contralateral relative to the laser hemisphere. (**e**) Schematic of head-fixation of mice in a virtual reality (VR) apparatus allowing 2-D navigation. Displacements of an air-suspended spherical ball in the anterior-posterior (and medial-lateral) axes of the mouse controlled y- (and x-) position movements in a visual VR environment. (**f**) Schematic of virtual corridor 6-cm in width and 330-cm in length, consisting of a start region (−10-0cm), an inhibition region (0-200cm) in which mice received unilateral 532-nm light on a random subset of trials (30%), a reward location (310cm) where mice received reward, and a teleportation location (320cm) where mice were transported to the start region following a variable ITI with mean of 2-s. (**g**) Average y-velocity (cm/s) across mice as a function of y-position (0-300cm in 25-cm bins) while navigating the virtual corridor on laser off (black) or laser on (green) trials in groups receiving DMS indirect (*left*, n = 7 mice, n = 1,712 laser off and n = 1,288 laser on trials) or direct pathway inhibition (*middle*, n = 6 mice, n = 1,088 laser off and n = 757 laser on trials), or illumination of the DMS in the absence of NpHR expression (*right*, no opsin, n = 5 mice, n = 1,178 laser off and n = 827 laser on trials). (**h**) Same as **g** but for average x-position (cm) contralateral to the unilaterally-coupled laser hemisphere. (**i**) Same as **g** but for view angle (degrees, contralateral to laser hemisphere). (**j**) Average across-mouse distance travelled (cm) to traverse the virtual corridor during laser off (black) or laser on (green) trials for mice receiving DMS indirect (n = 7 mice, n = 2,109 laser off and n = 1,574 laser on trials) or direct pathway inhibition (n = 6 mice, n = 1,330 laser off and n = 930 laser on trials), or DMS illumination in the absence of NpHR (n = 6 mice, n = 1,688 laser off and n = 1,199 laser on trials). Solid bars depict mean +/-S.E.M. across mice; grey lines indicate individual mouse mean.

### DMS pathway inhibition does not impact virtual corridor navigation

To determine if endogenous activity in DMS pathways provides bidirectional control of motor output in the absence of a decision, we carried out unilateral inhibition of indirect and direct pathways in head-fixed mice running on an air-supported ball to traverse a 2-dimensional linear corridor in virtual reality (VR) (**Fig. 1d-f**, 6-cm x 330-cm corridor). Illumination of the DMS was restricted to 0-200cm (laser on 30% of trials; hemisphere of illumination alternated across days). The parameters of the virtual corridor and inhibition period were selected to closely match the stem of the VR-based T-maze decision-making tasks that are the focus of subsequent experiments.

We found no detectable impact of pathway-specific DMS inhibition, nor DMS illumination alone, on multiple indicators of motor output during virtual corridor navigation. This included measures of velocity, x-position or view angle relative to the laser hemisphere, and distance traveled (**Fig. 1g-j**; see **Extended Data Fig. 2** for additional measures). Similarly, we obtained null effects of pathway-specific inhibition on velocity (and spatial preference) in freely behaving mice in a conditioned place preference assay (**Supplementary Fig. 3**).

These negative findings argue against a major involvement of endogenous activity in DMS pathways in the execution of movement in the absence of a decision. This is consistent with the dearth of reports demonstrating strong and opposing modulation of behavior by striatal pathways using pathway-specific optogenetic inhibition.

### Three virtual reality T-mazes with varying cognitive demands

We next considered the possibility that rather than contributing directly to a motor output, endogenous activity in DMS pathways may instead have opposing influence over decisions in a manner that is dependent on cognitive demand. To test this idea, we trained mice to perform a set of VR-based, decision-making tasks^26^ that shared identical motor readouts (left or right choice), had highly similar sensory environments, yet differed in their cognitive requirements (**Fig. 2a-b**).

**Figure 2.**
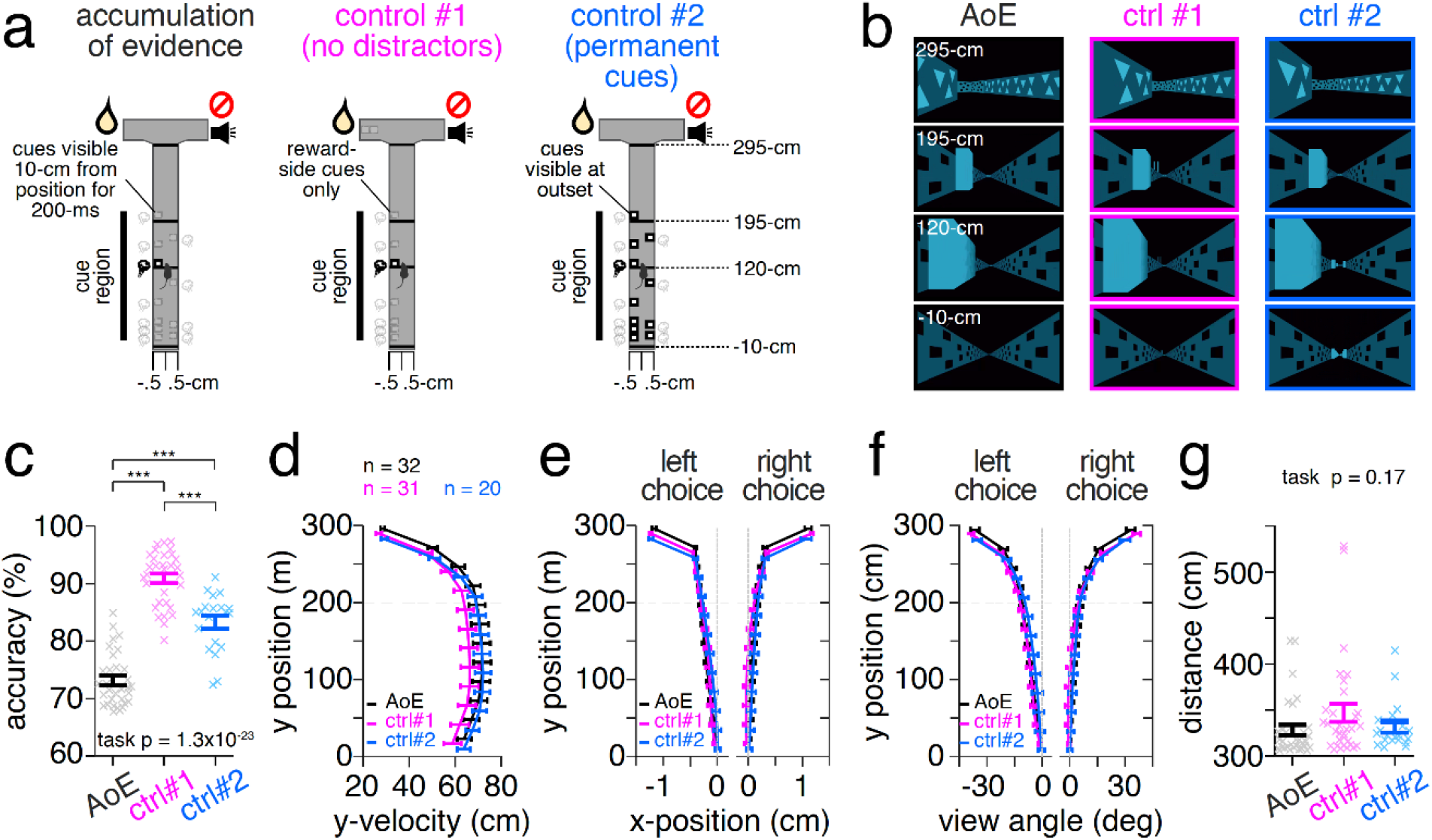
A set of virtual reality T-mazes have similar sensory features and identical motor requirements but different cognitive demands. (**a**) Schematic of three virtual reality (VR)-based T-maze tasks. (**b**) Example mouse perspective at the same maze position (−10cm, 120cm, 195cm, and 295cm) from the example trial depicted in **a** of the evidence accumulation (*left*, black), no distractors (*middle*, ctrl #1), or permanent cues (*right*, ctrl #2) tasks. (**c**) Average choice accuracy (% correct) across mice performing the accumulation of evidence (black, n = 32 mice, n = 52,381 trials), no distractors (magenta, ctrl #1: n = 31 mice, n = 56,783 trials), or permanent cues (cyan, ctrl #2: n = 20 mice, n = 27,870 trials) tasks. p-value denotes one-way ANOVA of task on accuracy (p = 1.3 × 10^−23^, F_2,80_ = 109.4). Asterisks indicate statistical significance of post-hoc, unpaired, two-tailed ranksum comparisons of accuracy between groups (*top to bottom*: ***p = 3.9 × 10^−7^, z = −5.1; ***p = 2.1 × 10^−11^, z = −6.7; ***p = 2.1 × 10^−5^, z = 4.3). (**d**) Average y-velocity (cm/s) across mice as a function of y-position (0-300 cm in 25cm bins) during performance of each task (colors and n as in **c**). (**e**) Same as **d** but for average x-position (cm) on left and right choice trials. (**f**) Same as **d** but for average view angle (degrees) on left and right choice trials. (**g**) Average distance (cm) traveled per trial across mice (evidence accumulation, n = 32 mice, n = 53,833 trials; no distractors (control #1): n = 32 mice, n = 60,074 trials; permanent cues (control #2): n = 20 mice, n = 29,192 trials). p-value reflects one-way ANOVA of task on distance (p = 0.16, F_2,81_ = 1.8). Throughout solid bars denote across mouse mean values +/-S.E.M. and transparent ‘x’ indicate individual mouse mean.

The first task was an “evidence accumulation” task, in which visuo-tactile cues were transiently presented on each side of the central stem of a virtual T-maze according to a Poisson distribution (“cue region”, 0-200cm), and mice were rewarded for turning to the maze side with the greater number of cues (**Fig. 2a,b**; black, *left*). Thus, mice were required to continually accumulate sensory cues over several seconds into a memory (or motor plan) that guided their left/right decision.

In two additional control tasks, we made modifications intended to weaken the cognitive demands of each task. In the first control task (“no distrators”), cues were presented on the rewarded maze side during the same maze region (0-200cm) according to the same Poisson distribution, but distractor cues on the non-rewarded arm side were omitted (**Fig. 2a-b**; magenta, *middle*). The absence of distractors on the non-rewarded side meant that each cue signaled reward with 100% probability, and thus gradual evidence accumulation was not required. Further ensuring that evidence accumulation was not required, an additional cue at the end of the maze was present only during the cue period (0-200cm) to signal the rewarded side.

In the second control task (“permanent cues”), the sensory statistics of the cues were identical to that in the evidence-accumulation task, but rather than *transient* visual cue presentation, visual cues were *permanently* visible from trial onset (**Fig. 2a-b**; cyan, *right*). This maintained the same conceptual task structure of the evidence accumulation task while decreasing the memory demands, as the sensory cues (or the motor plan) did not need to be remembered until the cues were passed.

We assessed how task demands impacted choice accuracy in each task. Consistent with the greatest cognitive and mnemonic demand in the evidence accumulation task, we found that overall choice accuracy was significantly lower compared to both control tasks (**Fig. 2c**, one-way ANOVA of task on accuracy, p = 1.3×10^−23^, F_2,80_ = 109.4; unpaired, two-tailed Wilcoxon ranksum test between evidence accumulation vs. no distractors, p = 2.1×10^−11^, z = −6.7, and evidence accumulation vs permanent cues, p = 3.9×10^−7^, z = −5.1).

While the motor requirements of a decision were the same across tasks (crossing an x-position threshold at the end of the central stem, see **Methods**), we examined the possibility that cross-task differences in cognitive requirements altered movement within the stem of the maze (0-300cm). We observed no consistent cross-task differences in velocity, x-position or view angle on left or right choice trials, nor distance traveled (**Fig. 2d-g**; see **Extended Data Fig. 3a-f** for additional measures). We further compared the relationship between behavior in the stem of the maze and choice across tasks by using a decoder to predict choice based on the trial-by-trial x-position or view angle (**Extended Data Fig. 3g-j**) at successive maze positions (0-300cm in 25-cm bins). While we were able to predict choice from either measure above chance levels in all three tasks (consistent with previous studies^26^), choice prediction accuracy was statistically indistinguishable across tasks (**Extended Data Fig. 3g-j**). Together, this indicated that cross-task differences in task demands did not prompt mice to systematically adopt distinct motor strategies.

### Behavioral effects of DMS pathway inhibition depend on task demand

We performed unilateral inhibition of DMS indirect and direct pathways restricted to the cue region (0-200-cm) of each task (**Fig. 3a-b**; laser on 10-20% of trials; hemisphere of illumination alternated across days). We found that inhibition of the indirect pathway produced a large bias towards contralateral choices during the accumulation of evidence task (**Fig. 3c** and **3d**, *left*), which was significantly greater than that observed in control animals that did not express opsin (**Fig. 3e**, average contralateral bias: DMS indirect, 42.3 +/- 4.4%, vs. no opsin, 5.9 +/- 3.6%). Similarly, inhibition of the direct pathway also produced a large choice bias during the accumulation of evidence task (**Fig. 3d**, *middle*; average contralateral bias: DMS direct, −36.8 +/- 8.6%), which was also significantly greater than that observed in control animals (**Fig. 3e**). However, in this case the direction of the choice bias was in the opposite (ipsilateral) direction to that observed with indirect pathway inhibition (also see **Extended Data Fig. 4a-i** for psychometric curves).

**Figure 3.**
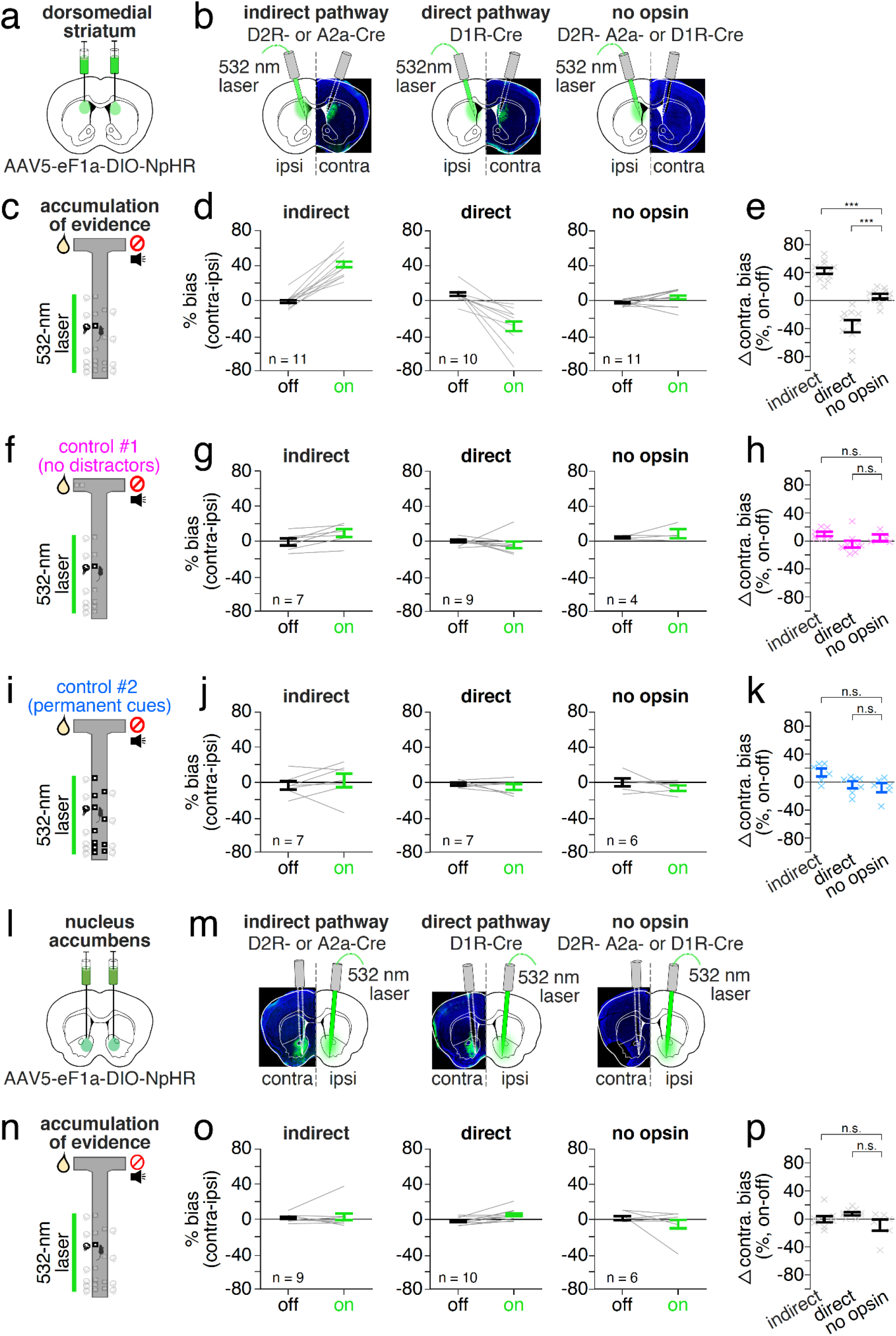
Inhibition of DMS but not NAc pathways has strong and opposing influence on choice during an evidence accumulation task, while having weaker effects during task variants with diminished cognitive demands. (**a**) Schematic of bilateral viral delivery of Cre-dependent NpHR to the dorsomedial striatum (DMS). (**b**) Schematic of bilateral fiberoptic implantation of the DMS and unilateral inhibition in behaving mice, with example histology from a mouse expressing NpHR in the indirect (*left*, D2R-/A2a-Cre) or direct (*middle*, D1R-Cre) pathways, or DMS illumination in the absence of NpHR (*right*, no opsin, A2a-/D2R- or D1R-Cre). 532-nm light (5-mW) was delivered unilaterally to the left or right hemisphere on alternate testing sessions and choice bias contralateral or ipsilateral to the hemisphere of inhibition was quantified. (**c**) Schematic of the evidence accumulation task with delivery of 532-nm light restricted to the cue region (0-200cm) on a random subset of trials (10-20%). (**d**) Average across mouse choice bias during the evidence accumulation task. Choice bias was defined as the difference between percent correct performance on trials when the correct choice was contralateral or ipsilateral to the inhibited hemisphere (% correct, contralateral-ipsilateral, positive values indicate a contralateral bias). Bias was calculated separately on laser off (black) and laser on (green) trials for mice receiving unilateral indirect pathway inhibition (*left*, n = 11 mice, n = 16,935 laser off and n = 3,390 laser on trials), unilateral direct pathway inhibition (*middle*: n = 10 mice; n = 14,030 laser off and n = 3,103 laser on trials), or unilateral illumination of the DMS in the absence of NpHR (*right*, n = 11 mice, n = 21,422 laser off and n = 5,113 laser on trials). (**e**) Difference in contralateral choice bias (% correct, contralateral-ipsilateral) between laser off and on trials (% bias, on-off) in mice performing the evidence accumulation task and receiving indirect pathway inhibition, direct pathway inhibition, or DMS illumination in the absence of NpHR. Asterisks indicate significance of unpaired, two-tailed Wilcoxon ranksum comparison of indirect to no opsin: ***p = 1.1×10^4^, z = 3.9; direct to no opsin: ***p = 2.2×10^4^, z = −3.7). (**f-h**) Same as **c-e** but for the no distractors (control #1) task. Indirect: n = 7 mice, n = 13,706 laser off and n = 3,288 laser on trials; direct: n = 9 mice, n = 14,647 laser off and n = 3,682 laser on trials; no opsin: n = 4 mice, n = 3,654 laser off and n = 901 laser on trials. Asterisks indicate significance of unpaired, two-tailed Wilcoxon ranksum comparison of indirect to no opsin: not significant (n.s.), p = 0.22, z = 1.2. Direct to no opsin: not significant (n.s.), p = 0.08, z = −1.8. (**i-k**) As in **c-e** but for the permanent cues (control #2) task. Indirect: n = 7 mice, n = 4,033 laser off and n = 929 laser on trials; direct: n = 7 mice, n = 6,061 laser off and n = 1,494 laser on trials; no opsin: n = 6 mice, n = 3,975 laser off and n = 923 laser on trials. Asterisks indicate significance of unpaired, two-tailed Wilcoxon ranksum comparison of indirect to no opsin: not significant (n.s.), p = 0.13, z = 1.5. Direct to no opsin: not significant (n.s.), p = 0.62, z = 0.5. (**l**) As in **a** but for bilateral viral delivery of Cre-dependent NpHR to the nucleus accumbens (NAc). (**m**) Same as **b** but for bilateral fiberoptic implantation of the NAc and unilateral inhibition in behaving mice, with example histology from a mouse expressing NpHR in the indirect (*left*, D2R-/A2a-Cre) or direct (*middle*, D1R-Cre) pathways, or NAc illumination in the absence of NpHR (*right*, no opsin, A2a-/D2R- or D1R-Cre). (**n-p**) As in **c** but for pathway-specific NAc inhibition during the accumulation of evidence task. Indirect: n = 9 mice, n = 11,978 laser off and n = 2,604 laser on trials; direct: n = 10 mice, n = 15,430 laser off and n = 3,348 laser on trials; no opsin: n = 7 mice, n = 9,819 laser off and n = 1,488 laser on trials. Asterisks indicate significance of unpaired, two-tailed Wilcoxon ranksum comparison of indirect to no opsin: not significant (n.s.), p = 0.86, z = 0.18; direct to no opsin: not significant (n.s.), p = 0.04, z = 2.0. Throughout solid bars denote across mouse mean values +/-S.E.M. and transparent ‘x’ indicate individual mouse mean. To account for multiple group comparisons we considered p-values significant after Bonferroni correction (2 comparisons).

Providing a stark contrast to the large effects of pathway-specific DMS inhibition on choice during the evidence accumulation task, inhibition of either pathway had significantly less impact on choice during the “no distractors” and “permanent cues” control tasks (**Fig. 3f-k**; also see **Extended Data Fig. 5a-c**; unpaired, two-tailed Wilcoxon ranksum test of indirect pathway inhibition evidence accumulation vs no distractors, p = 8.0×10^−4^, z = 3.4, or evidence accumulation vs permanent cues, p = 0.001, z = 3.3; and direct pathway inhibition evidence accumulation vs no distractors, p = 0.002, z = −3.1, or evidence accumulation vs permanent cues, p = 0.005, z = −2.8). In fact, the effects of pathway-specific DMS inhibition on choice bias in either control task did not significantly differ from those observed in control animals (**Fig. 3h**, for “no distractors”; **Fig, 3k**, for “permanent cues”; see also **Extended Data Fig. 4a-i** for psychometric curves).

Thus, inhibition of DMS pathways elicited strong and opposing effects on choice in the task with the greatest cognitive demand, which required the accumulation of sensory evidence across multiple seconds to arrive at a decision, and had a far limited impact on choice in task variants with reduced cognitive demand.

While DMS pathway inhibition had minimal impact on movement in a virtual corridor (**Fig. 1** and **Extended Data Fig. 2**), we considered the possibility that pathway-specific DMS inhibition altered motor performance in a manner that depended on the cognitive demands of the task. We found no cross-task differences in the effects of pathway-specific inhibition on measures of velocity, distance traveled, or per-trial standard deviation in view angle (**Extended Data Fig. 6a-i**). However, we found subtle but opposing effects of pathway-specific inhibition on average x-position and view angle (**Extended Data Fig. 6j-k**) in the evidence accumulation task. The direction of these biases was similar in the control tasks, but consistently smaller than in the evidence accumulation task. Thus, in line with the close relationship between x-position/view angle and choice in the absence of inhibition in each task (**Extended Data Fig. 3g-j**), pathway-specific DMS inhibition produced the same general pattern of cross-task effects on choice bias (**Extended Data Fig. 5b-d**) and x-position/view angle (**Extended Data Fig 6j-k**). As the quantitative relationship between x-position or view angle and choice is indistinguishable across tasks in the absence of neural inhibition (**Extended Data Fig. 3g-j**), cross-task differences in motor strategy does not provide a trivial explanation for these effects. Rather, taken together with the absence of an effect of pathway-specific DMS inhibition on motor output in the virtual corridor (**Fig. 1h-i**), these data imply that the effects of inhibition on behavior depends on cognitive demands.

### Little effect of NAc pathway inhibition on choice

We next sought to determine whether opponent control of choice by striatal pathways during the evidence accumulation task was specific to DMS, or if it extended to the ventral striatum. Towards this end, we delivered unilateral laser illumination to the nucleus accumbens (NAc) of mice expressing NpHR in the indirect or direct pathways (or non-opsin control mice), which was restricted to the cue-region (0-200cm) of the evidence accumulation task (**Fig. 3l-p**; see also **Extended Data Fig. 4j-l**).

Providing a clear functional dissociation between DMS and NAc, effects of pathway-specific NAc inhibition on choice bias were significantly smaller than those observed with inhibition of DMS pathways (**Extended Data Fig. 5e-f**; unpaired, two-tailed Wilcoxon ranksum test of DMS vs NAc indirect pathway inhibition, p = 2.6×10^−4^, z = 3.6; of DMS vs NAc direct pathway inhibition, p = 1.8×10^−4^, z = −3.7), and were also not significantly different from NAc control animals (**Fig. 3o-p**). It is unlikely that this dissociation between DMS and NAc can be explained by greater co-expression of pathway-specific markers in ventral versus dorsal striatum^29^, as both subregions exhibited equally low co-localization of D1R and D2R receptors (**Supplementary Fig. 1j-l**).

### Bernoulli GLM does not fully capture psychometric curves

Our inactivation experiments suggest that DMS pathways make strong contributions to behavior during a cognitively demanding evidence accumulation task, but do not contribute strongly to similar tasks with weaker cognitive demands. However, even during the evidence accumulation task, it is possible that the animals’ level of cognitive engagement varies over time. This raises the possibility that the contributions of the two pathways to behavior could change over time, even within the same task.

To address this possibility, we sought to understand the factors that contribute to decisions in the evidence accumulation task. As a first step, we used a Bernoulli generalized linear model (GLM) to predict choice based on a set of external covariates (**Fig. 4a-b**). These covariates included the sensory evidence (difference between the number of right and left cues, or “Δ cues”), the recent choice and reward history, the delivery of optical inhibition (“laser”), as well as a bias. Note that we set the value of the laser covariate to +1 (or −1) on trials with right (or left) hemisphere inhibition, and zero otherwise. A positive (or negative) GLM weight on this covariate thus captured an ipsilateral (or contralateral) “laser”-induced bias in choices relative to the hemisphere of inhibition. For the choice history covariates, a positive weight indicates a tendency toward repeating past choices (see **Methods** for details).

**Figure 4.**
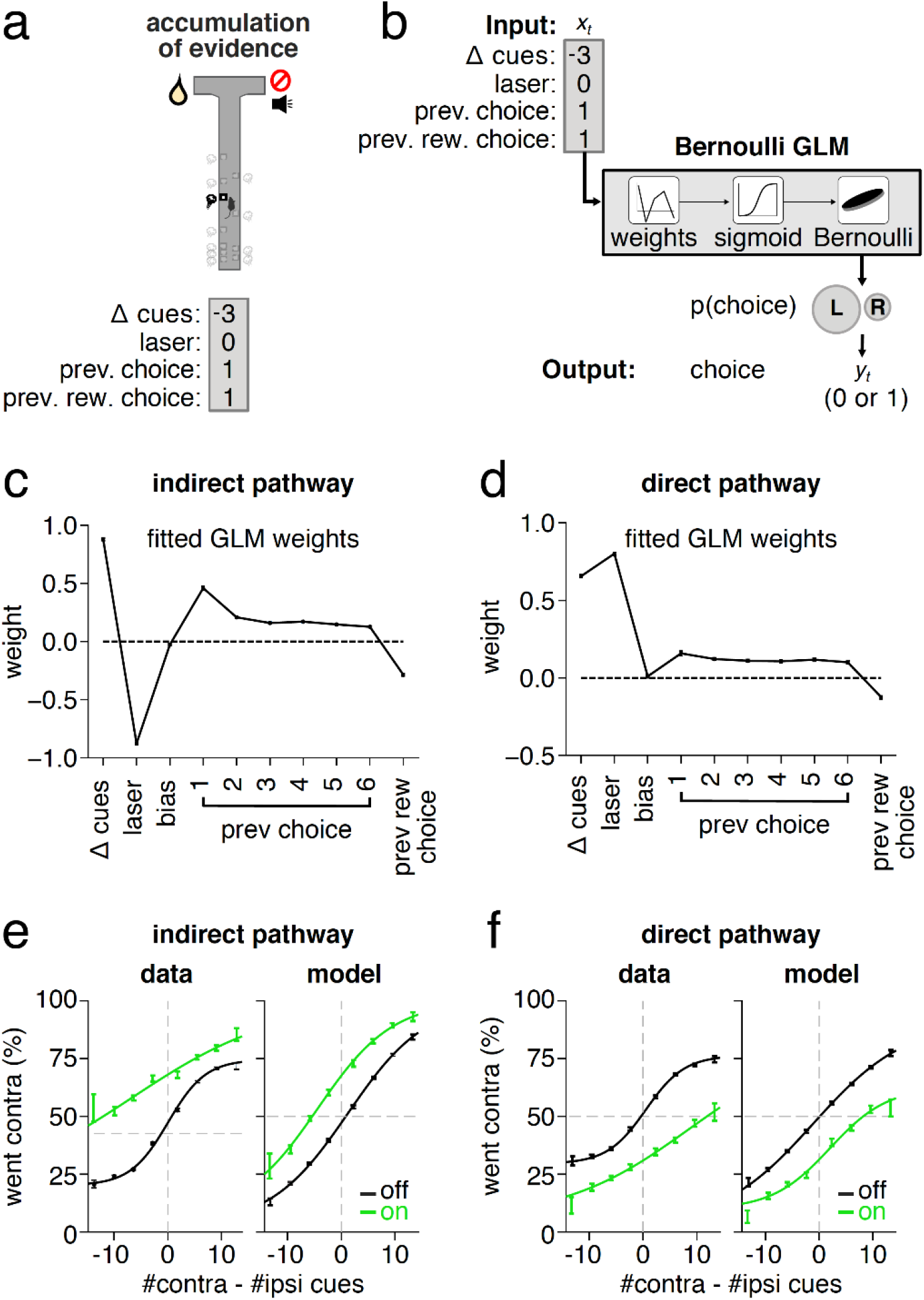
A GLM reveals that sensory evidence, DMS pathway inhibition, and trial history predict choice during the evidence accumulation task, but does not precisely recapitulate the shape of the psychometric curve. (**a**) Schematic of the evidence accumulation task and the coding of the external covariates for an example trial. (**b**) Schematic of the Bernoulli GLM for an example trial, showing the relationship between external covariates (inputs) and choice on each trial. On each trial, a set of GLM weights maps each input (Δ cues, laser, bias, previous choice, and existence of a previous rewarded choice) to the probability of each outcome through a sigmoid function, which gives the probability of a “righward” choice on the current trial. (**c**) Fitted GLM weights using aggregated data from all mice in the indirect pathway DMS inhibition group. The magnitude of each weight indicates the relative importance of that covariate in predicting choice, whereas the sign of the weight indicates the direction of the effect (e.g. a negative laser weight indicates that if inhibition is in the right hemisphere, the mice will be more likely to turn left, while a positive weight on previous choice indicates that if the previous choice was to the right, in the current trial this will bias the mice to turn right again). Error bars denote (+/-1) posterior standard deviation credible intervals. (**d**) Same as **c** but for mice receiving DMS direct pathway inhibition. (**e**) Fraction of contralateral choice trials as a function of the difference in contralateral versus ipsilateral cues for laser off (black) and on (green) trials, for mice receiving indirect pathway DMS inhibition for the data (left) and for simulations of the model (right). Error bars denote 95% confidence intervals around the fraction of choices in each bin of the data; solid curves denote logistic fits (n=13 mice, n = 46,313 laser off and n = 8,570 laser on trials). (**f**) Same as **e** but for the mice receiving direct pathway inhibition of the DMS (n=13 mice, n = 41,250 laser off and n = 7,927 laser on trials).

We fit the GLM to aggregated behavioral data from mice inhibited in each DMS pathway and found that sensory evidence, trial history, and optical inhibition all contributed to predicting choice (**Fig. 4c-d**). As expected, the effect of inhibition of each pathway was large and opposite in sign. However, the GLM did not accurately capture the animal’s psychometric curve, describing the probability of a rightward choice as a function of the sensory evidence (**Fig. 4e-f**). This led us to consider variants of the standard GLM that might better account for choice behavior.

### GLM-HMM better explains choice data with DMS inhibition

The standard GLM describes choice as depending on a fixed linear combination of sensory evidence, trial history, and optical inhibition. However, an alternative possibility is that mice use a weighting function that changes over time. To test this idea, we adopted a latent state model that allowed different GLM weights in different states, using the same external covariates as the standard 1-state GLM (**Fig. 4**). The model consists of a Hidden Markov Model (HMM) with Bernoulli Generalized Linear Model (GLM) observations, or GLM-HMM^30–33^ (**Fig. 5a-b**). Each hidden state is associated with a unique set of GLM weights governing choice behavior in that state. Probabilistic transitions between states occur after every trial, governed by a fixed matrix of transition probabilities (see **Methods** for details).

**Figure 5.**
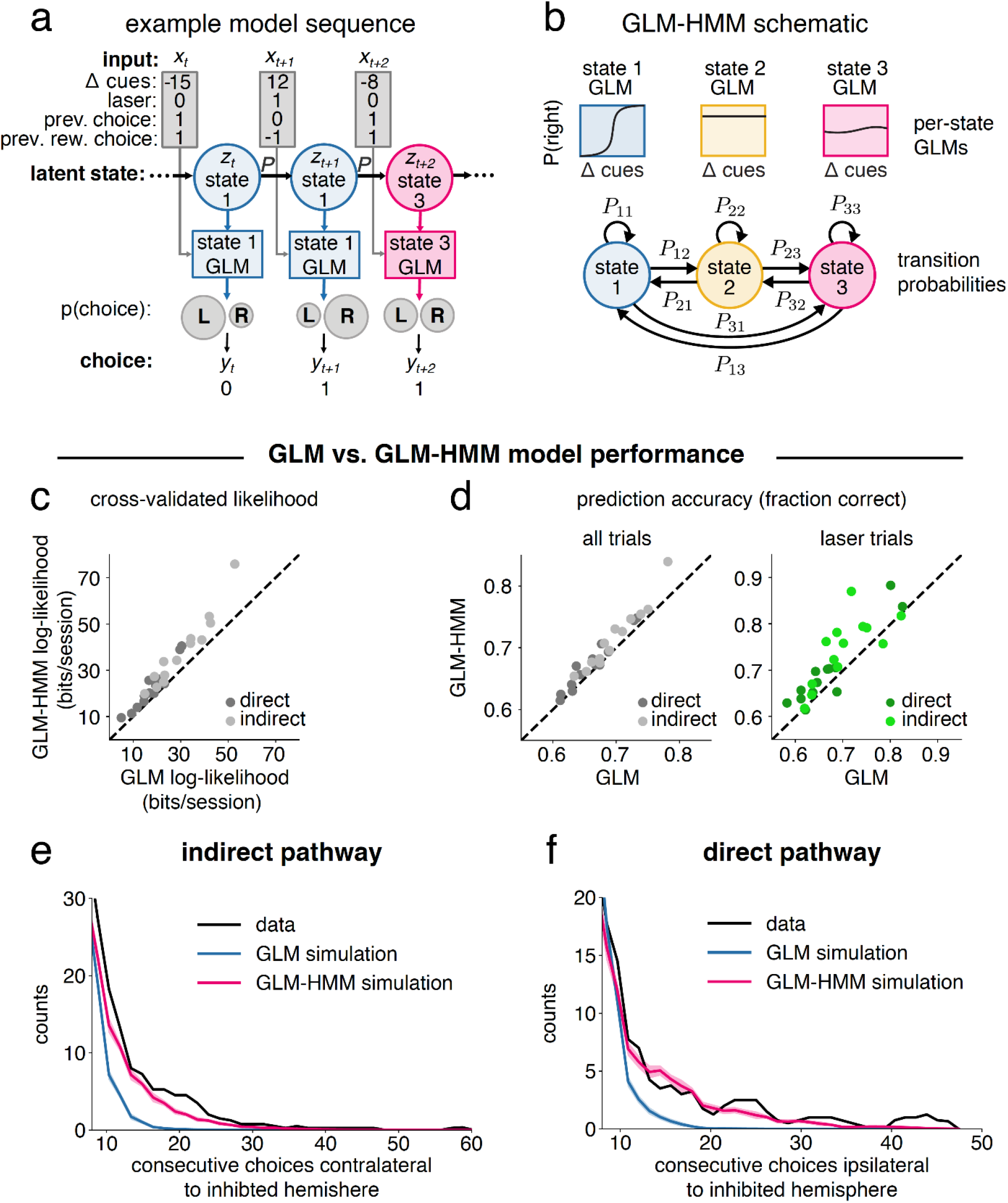
A GLM-HMM better explains choice during the evidence accumulation task than the GLM, particularly on trials with DMS pathway inhibition. (**a**) Example sequence of 3 trials of the evidence accumulation task, showing the relationship between external covariates (inputs), latent state, and choice on each trial. On each trial, the latent state defines which GLM weights map inputs (Δ cues, laser, previous choice, and previous rewarded choice) to the probability of choosing right or left. The transition probability *P* governs the probability of changing states between trials. See **Methods** for information on how the inputs were coded. (**b**) Schematic of GLM-HMM. The model has 3 latent states with fixed probabilities of transitioning between them. Each state is associated with a distinct decision-making strategy, defined by a mapping from external covariates, or inputs, such as Δ cues, to choice probability. (**c**) Cross-validated log-likelihood demonstrating the increased performance of the GLM-HMM over a standard Bernoulli GLM on held-out sessions. Dots represent model performance for individual mice (n=13 for each group). (**d**) Same as **c** but showing prediction accuracy as a fraction of the choices correctly predicted by each model across all trials (left) or on the subset of trials when the laser was on (right). (**e**) Histograms showing the number of consecutive laser trials for which the animal’s choice was in the same direction as the expected biasing effect of the laser (i.e. a choice contralateral to the laser hemisphere during DMS indirect pathway inhibition). Data (black), GLM simulation (blue), GLM-HMM simulation (pink). For the simulations, data of the same length as the real data was generated 100 times and the resulting histograms averaged. Curves denote smoothed counts using a sliding window average (window size = 3 bins). Shaded regions around the GLM and GLM-HMM curves indicate 95% confidence intervals. (**f**) Same as **e** but for mice receiving direct pathway inhibition of the DMS, therefore laser-biased choices are defined as those ipsilateral to the hemisphere of inhibition.

The GLM-HMM explained the choice data in the evidence accumulation task better than the GLM across multiple measures. We compared the likelihood of each animal’s data under the GLM-HMM to the standard Bernoulli GLM using cross-validation with held-out sessions (3-state GLM-HMM in **Fig. 5**; see **Extended Data Fig. 7a-e** for more information on model selection and demonstration that ~3-5 latent states was sufficient to reach a plateau in likelihood). The 3-state GLM-HMM achieved an average of 6.2 bits/session increase in log-likelihood, making an average session ~76 times more likely under the GLM-HMM (**Fig. 5c**). Furthermore, the GLM-HMM correctly predicted choice on held-out data more often than the GLM, especially on laser trials (**Fig. 5d**; average improvement across mice of 1.6% on all trials, 3.5% on trials with optical inhibition, and 4.1% on trials with optical inhibition when considering only mice with at least 100 inhibition trials).

Most interestingly, the GLM-HMM was better able to capture the temporal structure in the effect of inhibition on choice. Specifically, the choice data contained long runs in which choice was consistent with the bias direction predicted by pathway-specific inhibition (“laser”), a feature which GLM-HMM simulations recapitulated, but GLM simulations did not (**Fig. 5e-f**). Thus, taken together, the GLM-HMM provided a better model of the choice data than a standard GLM, particularly on trials with pathway-specific DMS inhibition.

### GLM-HMM identifies states with varying DMS-dependence

We examined the state-dependent weights of the GLM-HMM and found substantial differences across states in the weighting of sensory evidence, previous choice, and most intriguingly, inhibition of DMS pathways (**Fig. 6a-b**). In particular, two of the three states (states 1 and 2) displayed a large weighting of sensory evidence on choice, while the “laser” weight was large only in state 2. In contrast, in state 3 choice history had a larger weight than in the other states, and neither sensory evidence nor “laser” had much influence on choice.

**Figure 6.**
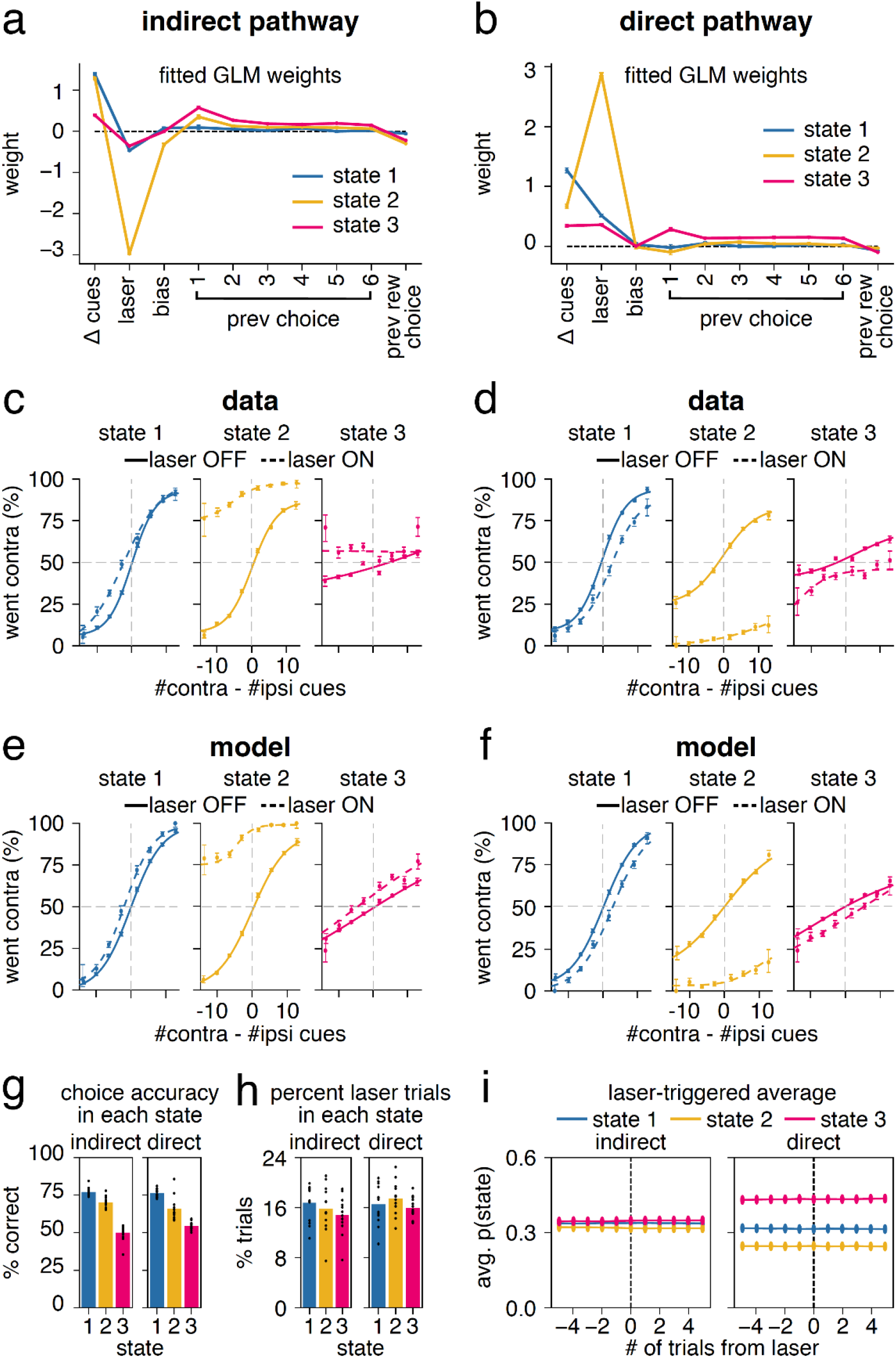
A GLM-HMM uncovers states during the evidence accumulation task with different weighting on sensory evidence, choice history, and DMS pathway inhibition. (**a**) Fitted GLM weights for 3-state model from mice in the indirect pathway DMS inhibition group. Error bars denote (+/-1) posterior standard deviation for each weight. The magnitude of the weight represents the relative importance of that covariate in predicting choice, whereas the sign of the weight indicates the side bias (e.g. a negative laser weight indicates that if inhibition is in the right hemisphere, the mice will be more likely to turn left, while a positive weight on previous choice indicates that if the previous choice was to the right, in the current trial this will bias the mice to turn right again). (**b**) Same as **a** but for the direct pathway group. (**c**) Fraction of contralateral choices as a function of the difference in contralateral versus ipsilateral cues in each trial for mice in the indirect pathway inhibition group. To compute psychometric functions, trials were assigned to each state by taking the maximum of the model’s posterior state probabilities on each trial. Error bars denote +/-1 SEM for light off (solid) and light on (dotted) trials. Solid curves denote logistic fits to the concatenated data across mice for light off (solid) and light on (dotted) trials. (**d**) Same as **c** but for the mice receiving direct pathway inhibition of the DMS. (**e**) Same as **c** but for data simulated from the model fit to mice receiving indirect pathway inhibition of the DMS (see **Methods**). (**f**). Same as **e** but for mice receiving direct pathway inhibition of the DMS. (**g**) Performance in each state for mice receiving DMS inhibition in the indirect pathway (left) and direct pathway (right), shown as the percentage of total trials assigned to that state in which the mice made the correct choice. Colored bars denote the average performance across all mice. Black dots show averages for individual mice (n=13 mice for both groups). (**h**) Percentage of laser-on trials that the model assigned to each state for mice receiving DMS inhibition in the indirect pathway (left) and direct pathway (right). Colored bars denote the average performance across all mice. Black dots show averages for individual mice (n=13 mice for both groups). (**I**) The posterior probability of each state for the five trials before and after a laser-on trial, averaged across all such periods (n=8570, indirect; n=7927, direct).

To characterize state-dependent psychometric performance, we used the fitted model to compute the posterior probability of each state given the choice data and assigned each trial to its most probable state (**Fig. 6c-d**). We then examined the psychometric curves for trials assigned to each state. In state 3, performance was low (**Fig. 6g**) and DMS inhibition had little effect on behavior (**Fig. 6c-d**). This is consistent with the high GLM weight on choice history in this state, and low weights on sensory evidence and laser (**Fig. 6a-b**). This implies relatively little contribution of DMS pathways during a task-disengaged state when mice pursued a strategy of repeating previous choices rather than accumulating sensory evidence. When considered together with comparisons of the effect of pathway-specific DMS inhibition in control T-maze tasks where performance is high (**Fig. 2c**) but effects of inhibition are limited (**Fig. 3f-k**; **Extended Data Fig. 5b-c**), this implies a dissociation between task performance and the contributions of DMS pathways to behavior.

Compared to state 3, sensory evidence heavily modulated behavior in both states 1 and 2, and performance was accordingly high (**Fig. 6c-d,g**). Interestingly, the effect of DMS pathway inhibition was much larger in state 2. These results were again consistent with the GLM weights: both state 1 and 2 had high weighting of sensory evidence, low weighting of choice history, but greatly differed in their weighting of the “laser” (**Fig. 6a-b**). The discovery of state 2 implies that DMS pathways contribute most heavily to choices in a state in which mice are pursuing a strategy of evidence accumulation, consistent with cross-task comparisons of the effects of inhibition (**Fig. 3**). The discovery of state 1, which differed most noticeably from state 2 in the extent that the laser affected choice, may suggest the existence of another neural mechanism for evidence accumulation with minimal DMS dependence.

We found that GLM-HMM simulations closely recapitulated these state-dependent psychometric curves (**Fig. 6e-f**). This not only validated our fitting procedure, but also provided additional evidence that a multi-state model provides a good account of the animals’ decision-making behavior during the evidence accumulation task.

While the effect of the laser differed across states, the probability of being in a particular state did not change on or after trials with optical inhibition (**Fig. 6i**), implying that DMS pathway inhibition itself did not generate transitions between states. In addition, the fraction of trials with optical inhibition was equivalent across states (~15% of all trials in each state; **Fig. 6h**). This implies that the model did not identify states simply based on the presence of laser trials.

We obtained similar states when fitting the model to a combined dataset including all groups of mice (those receiving DMS indirect and direct pathway inhibition, as well as control mice receiving DMS illumination in the absence of NpHR, **Extended Data Fig. 7f**). As when fitting each group separately, the combined fit revealed that both inhibition groups contained a single state with large weights on sensory evidence and the laser. In contrast, the control mice had small laser weights across all three states.

We also examined the results of fitting the 4-state GLM-HMM (**Extended Data Fig. 7d-e**), given it had a slightly higher cross-validated log-likelihood than the 3-state model (**Extended Data Fig. 7a**). In this case, the weights for states 1 and 2 were very similar to the 3-state model; the key difference was that the choice history state (state 3 of the 3-state model), was further subdivided into two states that differed in having a slight contralateral versus a slight ipsilateral bias. This suggests that while the model may uncover finer-grained structure in the data beyond 3-states (**Extended Data Fig. 7d-e**), these states yield diminishing interpretive insight on the weighting of sensory evidence, choice history, and DMS pathway inhibition across time.

### Diversity in timing and number of GLM-HMM state transitions

The fitted transition matrix revealed a high probability of remaining in the same state across trials (**Fig. 7a-b**). These transition probabilities produced a diversity in the timing and number of state transitions across sessions, which we visualized by calculating the maximum posterior probability of each state on each trial (**Fig. 7c-d**; see **Methods**, **GLM-HMM**). In some sessions, mice persisted in the same state, while in many sessions, mice visited two or even all three states (example sessions in **Fig. 7c-d**; summaries of state occupancies across sessions in **Fig. 7e-h**; and summary of all individual mice in **Supplementary Fig. 4**). Average single-state dwell times ranged from 39-86 trials (**Fig. 7g**). This was shorter than the average session length of 194 trials, consistent with visits to multiple states per session.

**Figure 7.**
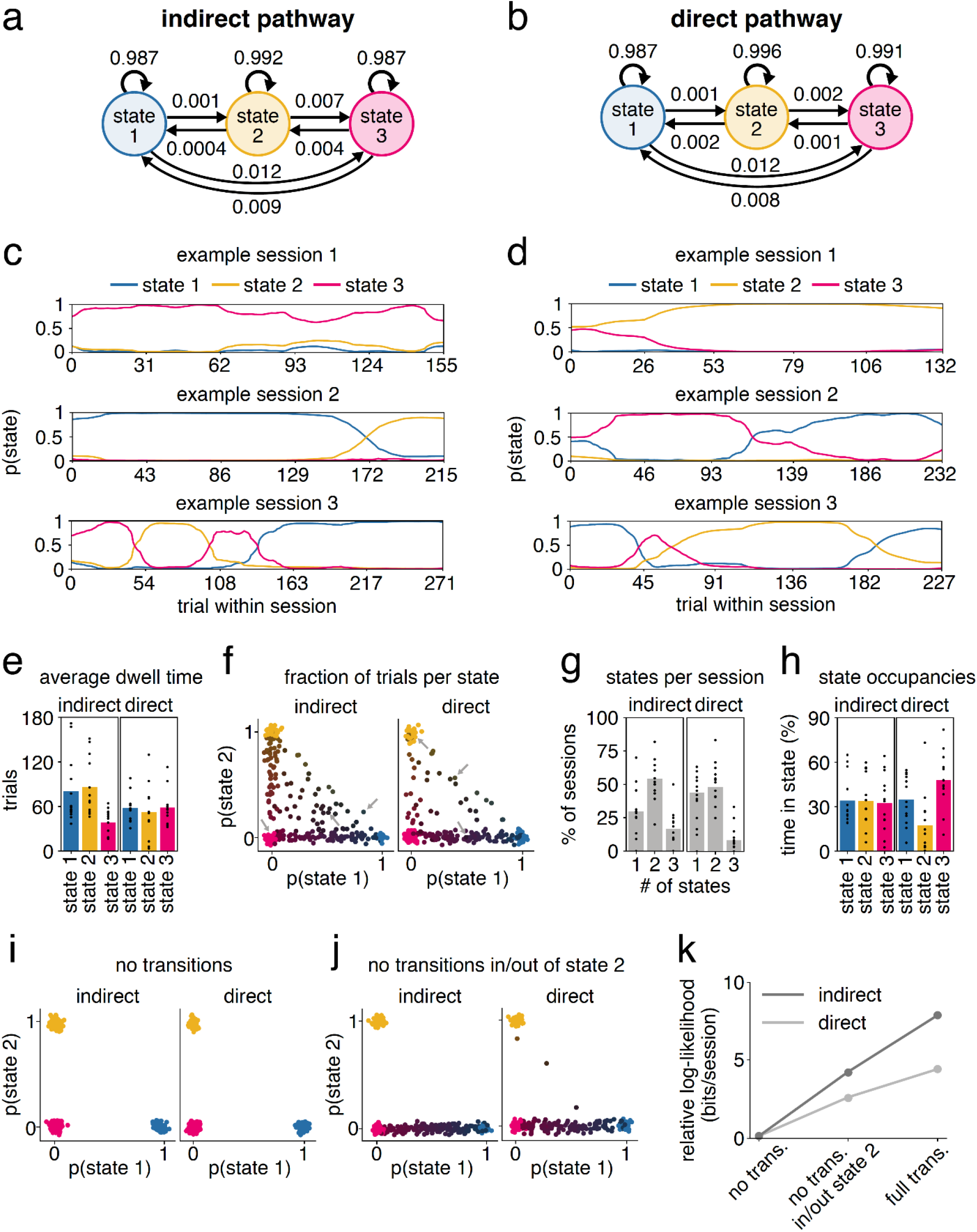
Diversity across sessions in the timing and number of GLM-HMM state transitions. (**a**) Transition probabilities for the indirect pathway group. (**b**) Same as **a** but for the direct pathway group. (**c**) The posterior probability of being in each state for each trial for 3 example sessions from a mouse in the indirect pathway group. (**d**) Same as **c** but for two mice from the direct pathway group. (**e**) Dwell times showing the average consecutive number of trials that the mice spent in each state for mice with indirect (left; range 39-86 trials, average session length 202 trials) and direct (right; range 52-59 trials, average session length 185 trials) pathway inhibition. Black dots show averages for individual mice (n=13 mice for both groups). (**f**). The fraction of trials that the mice spent in each state in each session. Each dot represents an individual session (n=271, indirect pathway; n=266, direct pathway). Color-coding reinforces the state composition of each session (e.g. blue indicates the mouse spent 100% of the session in state 1). A small amount of Gaussian noise was added to the position of each dot for visualization purposes. Grey arrows identify the example sessions shown in **c** and **d**. (**g**) The fraction of sessions in which the mice entered one, two, or all three states. Gray bars denote the average fraction of sessions for all mice. Black dots show averages for individual mice (n=13 mice for both groups). (**h**) Time spent in each state represented as a percentage of total trials for mice inhibited in the indirect pathway (left) and direct pathway (right). Colored bars denote the average state occupancies across all mice. Black dots show averages for individual mice (n=13 mice both groups). (**i**) Same as **f** except state assignments were obtained from a model in which the transition probabilities were restricted to disallow transitions between states (i.e. all off-diagonal transition probabilities equal zero; see **Methods**). (**j**) Same as **f** except state assignments were obtained from a model in which transitions were disallowed between state 2 and the other states. (**k**) Comparison of the cross-validated log-likelihood of the data when fitting GLM-HMMs with the reduced models from **i** and **j**, relative to the log-likelihood of the full model, in bits per session.

While individual sessions were heterogeneous in terms of their state occupancies, averaged across sessions, the posterior probability of being in each state tended to be stable across trials (**Fig. 7e**; **Extended Data Fig. 8a-b**). Model simulations recapitulated these state transition characteristics, including dwell times and state occupancies (**Extended Data Fig. 9**), further indicating our model captures latent structure in our data.

One notable exception in the stability of posterior probabilities of each state across time was an increase in state 3 probability towards the end of a session (**Extended Data Fig. 8a**), potentially reflecting a decrease in task engagement related to reward satiety. Consistent with a relationship to satiety, within-session transitions into state 3 were associated with higher amounts of previously accumulated reward and higher preceding rates of reward (**Extended Data Fig. 8c-d,e-f**). In addition, while the posterior probability of each state showed minimal modulation surrounding a rewarded trial, the probability of state 3 was much more likely surrounding trials with excess travel (**Extended Data Fig. 8g-h,i-j**), an indicator of non-goal directed movement and task disengagement. Indeed, the probability of state 3 gradually increased and decreased approximately 25 trials prior to and following excess travel trials, consistent with the average state 3 dwell time (**Fig. 7e**).

Given the presence of sessions in which mice occupied a single state, we considered model variants that disallowed within-session state transitions. Our goal was to determine if these variant models could provide a better explanation of the data, or alternatively, if within session state transitions are in fact an important structural feature for explaining the data. In one model variant, we disallowed transitions between states entirely (**Fig. 7k**). In the other, we tested the possibility that state 2, which is unique in the strength of its laser weight, captured a session-specific feature of inhibition by disallowing transitions in and out of that state (**Fig. 7l**). Using cross-validation, we found that neither alternative model explained the data as well as a model with unrestricted transitions (**Fig. 7m**), indicating that within-session transitions between states was an important feature of the model.

### Motor performance across GLM-HMM states

Given the close relationship between excess travel and the posterior probability of state 3, we considered the possibility that other measures of motor behavior varied across states. We found that on trials without DMS pathway inhibition (**Extended Data Fig. 10a-g,o-u**), mice exhibited no obvious differences across states in velocity, x-position, or view angle (**Extended Data Fig. 10a-d,o-r**). However, during state 3 relative to state 1 and 2, we observed an increased tendency in measures of non-goal directed movements (**Extended Data Fig 10e-g,s-u**). This is consistent with the higher probability of state 3 around trials with excess travel (**Extended Data Fig. 8h,j**), and the interpretation of state 3 as a task-disengaged state.

We also considered the possibility that DMS pathway inhibition had state-dependent effects on motor output (**Extended Data Fig. 10h-n,v-bb**). We observed limited effects of inhibition on velocity, per-trial standard deviation in view angle, and distance traveled across all three states. However, similar to our cross-task comparisons (**Extended Data Fig. 6j,k**), DMS pathway inhibition produced a small but opposing bias in average x-position (**Extended Data Fig. 10j,x**) and view angle (**Extended Data Fig. 10k,y**), which was greatest in the state with the largest laser weight (state 2, **Fig. 6**). This is consistent with our conclusions that the effects of DMS inhibition on behavior are state-dependent, and that x-position and view angle are closely linked indicators of choice in the context of VR-based T-maze tasks (**Extended Data Fig. 3g-j**).

## DISCUSSION

Our findings indicate that opposing contributions of DMS pathways to movement is minimal in the absence of a decision (**Fig. 1**), while the pathways provide large and opponent contributions to decision-making. Moreover, this contribution depends on the demands of a task (**Fig. 2**), as the effect of inhibition is much larger during decisions that require evidence accumulation relative to control tasks with weaker cognitive requirements yet similar sensory features and motor requirements (**Fig. 3**). The GLM-HMM further revealed that even within the evidence accumulation task, the contribution of DMS pathways to choice is not fixed. For example, DMS pathways have little contribution when mice pursue a strategy of repeating previous choices during the evidence accumulation task (**Fig. 6**). Thus, together our findings imply that opposing contributions of DMS pathways to behavior are task- and state-dependent.

### Cross-task differences in effects of DMS pathway inhibition

Our finding that DMS activity contributes to the evidence accumulation task, but not to task variants with weaker cognitive demands, is broadly consistent with previous work based on lesions, pharmacology, and recordings implicating DMS in short-term memory and the dynamic comparison of the value of competing options^19–22,25,34–37^. But then why have most previous optogenetic pathway-specific manipulations emphasized an opposing role for DMS pathways in the direct control of motor output^2,3,16,4,8,11^, rather than on decision-making? Prior work has overwhelmingly relied on the synchronous activation of striatal pathways, as opposed to the inhibition employed here. While DMS pathway activation is sufficient to bias movements such as spontaneous rotations, we observed relatively little impact of inhibition on behavior in the absence of a decision (**Fig. 1**). Taken together with previous work, our findings may thus imply limits in the use of artificial activations in assessing striatal pathway function. Our results may also imply that DMS pathways would not necessarily display opposing correlates of movements^14,18,38–41^, but rather opposing correlates of a decision process^12,13,20,22,34,37^.

While our observations are consistent with the classic view of opposing contributions of striatal pathways to behavior^1^, several prominent studies have instead challenged this view by reporting non-opposing behavioral effects of activating each pathway^42–48^. This may be because the pathways of a specific striatal subregion only exert opposing control on behavior in a specific context, for example during cognitively demanding decision-making as shown here for the DMS, or during an interval timing task that requires the proactive suppression of actions, as shown for the dorsolateral striatum^28^. Along these lines, our comparison to NAc pathways, where inhibition produced weak effects on behavior in similar directions (**Fig. 3p**), may imply that we have not discovered the context in which NAc pathways have opposing contributions to behavior.

We designed our T-maze tasks to have very similar sensory features and identical motor requirements, and yet very different cognitive demands, as assessed by task accuracy. That being said, the sensory features were not identical. Therefore, while unlikely, we cannot rule out that the subtle sensory differences contributed to the cross-task differences in the effect of pathway inhibition. A future direction would be to maintain an identical sensory environment across tasks and instead change the decision-making rule to be more cognitively demanding.

### Within-task changes in effects of DMS pathway inhibition

In addition to our cross-task comparison, we reveal the novel insight that mice occupy time-varying latent states within a single task and that the contribution of DMS pathways to choice depends on the internal state of mice. The application of a GLM-HMM was critical in uncovering this feature of behavior, allowing the unsupervised discovery of latent states that differ in how external covariates were weighted to influence a choice^32,49,50^. This provided two insights on the contributions of DMS pathways to behavior.

First, the impact of DMS inhibition was diminished when mice occupied a task-disengaged state in which choice history heavily predicted decisions, while conversely, the impact of DMS inhibition was accentuated when mice occupied a task-engaged state in which sensory evidence strongly influenced choice (**Fig. 6**). This strengthens our conclusion from our cross-task comparison, which is that DMS pathways have a greater contribution to behavior when actively accumulating evidence towards a decision output.

Second, mice occupied two similar task-engaged states that were modestly distinguished in overall accuracy (**Fig. 6g**) and prominently distinguished by the influence of DMS inhibition on choice (**Fig. 6a-d**). While transitions between these two states were relatively rare on the same day, there were days that included both states (**Fig. 7**). The discovery of these two states leads to the intriguing suggestion that mice are capable of accumulating evidence towards a decision in at least two neurally distinct manners: one that depends on each DMS pathway (state 2), and another that does not (state 1). This is reminiscent of demonstrations that neural circuits have significant capacity for compensation to perturbations^51,52^, and our modeling approach may provide a new avenue for the identification of such compensatory mechanisms that will facilitate their future study.

While our work focused on the 3-state GLM-HMM, our conclusions do not depend on assuming exactly three states. In fact, the cross-validated log-likelihood of our data is higher for four states than three. Yet the conclusions from the 4-state model were similar to those from the 3-state model (compare **Fig 6a-b** to **Extended Data Fig. 7d-e**), and additional gains in log-likelihood decrease significantly for larger numbers of states. Nevertheless, it will be important for future work to compare the GLM-HMM framework employed here, which assumes discrete states, to models that assume continuously varying states^33,53^.

All together, our studies provide new perspectives on the neural mechanisms by which DMS pathways exert opponent control over behavior, with particular emphasis on the importance of accounting for task demands, internal state and associated behavioral strategies when assessing neural mechanisms. Toward this end we expect our behavioral and computational frameworks to be of broad utility in uncovering the neural substrates of decision-making in a wide range of settings.

## Acknowledgements

We would like to thank the entire BRAINCoGs team as well as the Witten and Pillow labs for feedback on this work. We thank S. Stein and S. Baptista for technical support in animal training, and C. Kopecs for technical assistance. This work was supported by grants from NIH R01 DA047869 (IBW), F32MH118792 (SSB), F32NS101871 (LP), K99MH120047 (LP), U19 NS104648-01 (JWP, IBW), ARO W911NF1710554 (IBW), Brain Research Foundation (IBW), Simons Collaboration on the Global Brain (JWP, IBW), 1R01MH106689 (IBW), and the New York Stem Cell Foundation (IBW). IBW is a NYSCF—Robertson Investigator.

## Author Contributions

S.S.B. performed the experiments. I.R.S. developed the GLM-HMM. S.S.B. and I.R.S. analyzed the data with guidance from J.W.P. and I.B.W. L.P., Z.C.A., and B.E. provided technical and analysis support. J.M.I., A.L.H., and P.S. aided behavioral training. A.B. performed histology. J.C. provided support for electrophysiology. C.A.Z. and J.R.C provided support for the virtual corridor. S.S.B. and I.B.W. conceived the experimental work. S.S.B., I.R.S., J.W.P. and I.B.W. interpreted the results and wrote the manuscript.

## METHODS

### Animals

For optogenetic experiments we used both male and female transgenic mice on heterozygous backgrounds, aged 2-6 months of age, from the following three strains backcrossed to a C57BL/6J background (Jackson Laboratory, 000664) and maintained in-house: Drd1-Cre (n = 45, EY262Gsat, MMRRC-UCD), Drd2-Cre (n = 24, ER44Gsat, MMRRC-UCD), and A2a-Cre (n = 18, KG139Gsat, MMRRC-UCD). An additional 35 mice were excluded from all optogenetic analyses due to failed task acquisition (n = 11 mice) or failed viral/fiberoptic targeting of DMS (n = 8) or NAc (n = 16). An additional 4 Drd1-Cre mice, 3 A2a-Cre mice, and 2 Drd2-Cre mice were used for electrophysiological characterization of halorhodopsin (NpHR)-mediated inhibition, or fluorescent *in situ* hybridization (FISH) characterization of Cre expression profiles. FISH experiments also utilized 2 Drd1a-tdTomato mice (Jax, 016204). Mice were co-housed with same-sex littermates and maintained on a 12-hour light – 12-hour dark cycle. All surgical procedures and behavioral training occurred in the dark cycle. All procedures were conducted in accordance with National Institute of Health guidelines and were reviewed and approved by the Institutional Animal Care and Use Committee at Princeton University.

### Surgical procedures

All mice underwent sterile stereotaxic surgery to implant ferrule coupled optical fibers (Newport, 200 μM core, 0.37 NA) and a custom titanium headplate for head-fixation under isoflurane anesthesia (5% induction, 1.5% maintenance). Mice received a preoperative antibiotic injection of Baytril (5mg/kg, I.M.), as well as analgesia pre-operatively and 24-hours later in the form of meloxicam injections (2mg/kg, S.C.). A microsyringe pump controlling a 10μl glass syringe (Nanofill) was used to bilaterally deliver virus targeted to either the DMS (0.74 mm anterior, 1.5 mm lateral, −3.0 mm ventral) or the NAc (1.3 mm anterior, 1.2 mm lateral, −4.7 mm ventral). For optogenetic inhibition, the following viruses were used: AAV5-eF1a-DIO-eNpHR3.0-EYFP-WPRE-hGH (UPenn, 1.3 × 10^13^ parts/mL) or AAV5-eF1a-DIO-eNpHR3.0-EYFP-WPRE-hGH (PNI Viral Core, 2.2 × 10^14^ parts/mL, 1:5 dilution). For fluorescence *in situ* hybridization experiments, AAV5-eF1a-DIO-EYFP-hGHpA (PNI Viral Core, 6.0 × 10^13^ parts/mL) was used to label D1R^+^ and D2R^+^ neurons in D1R-Cre and A2A-Cre transgenic lines. In all experiments, virus was delivered at a rate of 0.2 μL/min for a total volume of 0.3-0.7 μL in the DMS, or 0.3-0.4 μL in the NAc. To accommodate patch fiber coupling, optical fibers were implanted at angles (DMS: 15°, 0.74 mm anterior, 1.1 mm lateral, −3.6 mm ventral; NAc: 10°, 1.3 mm anterior, 0.55 mm lateral, −5.0 ventral) and were then fixed to the skull using dental cement. Mice were allowed to recover and closely monitored for 5 days before beginning water-restriction and behavioral training.

### Optrode recording for NpHR validation

Following the surgical procedures described above, Cre-dependent NpHR was virally delivered bilaterally to the DMS of mice (n = 3 A2a-Cre; n = 2 D1R-Cre) via small (~300-uM) craniotomies made using a carbide drill (**Extended Data Fig. 1a**). The craniotomies were filled with a small amount of silicon adhesive (Kwik-Sil, World Precision instruments) and then covered with UV-curing optical adhesive (Norland Optical Adhesive 61), while a custom-designed headplate for head-fixation was cemented to the skull. Following a recovery period of >4 weeks, awake mice were head-fixed on a plastic running wheel attached to a breadboard via Thorlabs posts and holders, which was fixed immediately adjacent to a stereotaxic setup (Kopf) enclosed within a Faraday cage (**Extended Data Fig. 1b**). Silicon and optical adhesive was removed from the craniotomies and a 32-channel, single-shank silicon probe (A1×32-Poly3, NeuroNexus) coupled to a tapered optical fiber (65 uM, 0.22 NA) was stereotaxically inserted under visual guidance of a stereoscope and allowed to stabilize for ~30 minutes. Signals were acquired at 20 kHz using a digital headstage amplifier (RHD2132, Intan) connected to an RHD USB data acquisition board (C3100, Intan). A screw implanted over the cerebellum served as ground. Continuous signal was imported into MATLAB for referencing to a local probe channel and high-pass filtering at 200 Hz, and then imported into Offline Sorter v3 (Plexon) for spike thresholding and single-unit sorting. During recording, the optical fiber was connected via a patch cable to a 532-nm laser, which was triggered by a TTL pulse sent by a pulse generator controlled by a computer running Spike2 software. TTL pulse times were copied directly to the RHD USB data acquisition board. Laser sweeps consisted of forty deliveries of 5-s light (5-mW, measured from fiber tip), separated by 15-s intertrial intervals. From 1-3 recordings were made at different depths within a single probe penetration (minimum separation of 300-uM), with each hemisphere receiving 1-3 penetrations at different medial-lateral or anterior-posterior coordinates. For recordings in mice carried out over multiple days, craniotomies were filled with Kwik-Sil and covered with silicone elastomer between recordings (Kwik-Cast, World Precision Instruments).

### VR Behavior

#### Virtual reality setup

Mice were head-fixed over an 8-inch Styrofoam® ball suspended by compressed air (~60 p.s.i.) facing a custom-built Styrofoam® toroidal screen spanning a visual field of 270° horizontally and 80° vertically. The setup was enclosed within a custom-designed cabinet built from optical rails (Thorlabs) and lined with sound-attenuating foam sheeting (McMaster-Carr). A DLP projector (Optoma HD141X) with a refresh rate of 120 Hz projected the VR environment onto the toroidal screen (**Fig. 1e**).

An optical flow sensor (ADNS-3080 APM2.6), located beneath the ball and connected to an Arduino Due, ran custom code to transform real-world ball rotations into virtual-world movements (https://github.com/sakoay/AccumTowersTools/tree/master/OpticalSensorPackage) within the Matlab-based ViRMEn^54^ software engine (http://pni.princeton.edu/pni-software-tools/virmen). The ball and sensor of each VR rig were calibrated such that ball displacements (*dX* and *dY*, where X (and Y) are parallel to the anterior-posterior (and medial-lateral) axes of the mouse) produced translational displacements proportional to ball circumference in the virtual environment of equal distance in corresponding X and Y axes. The y-velocity of the mouse was given by 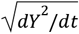, where *dt* was the elapsed time from the previous sampling of the sensor. The virtual view angle of mice was obtained by first calculating the current displacement angle as: ω = *atan*2(− *dX* · *sign*(*dY*), |*dY*|). Then the rate of change of view angle (*θ*) for each sampling of the sensor was given by:

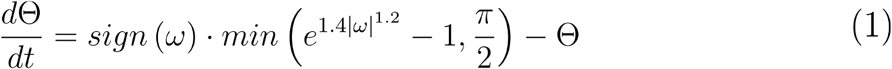

This exponential function was tuned to (1) minimize the influence of small ball displacements and thus stabilize virtual-world trajectories, and (2) increase the influence of large ball displacements in order to allow sharp turns into the maze arms^26^.

Reward and whisker air puffs were delivered by sending a TTL pulse to solenoid valves (NResearch), which were generated according to behavioral events on the ViRMEn computer. Each TTL pulse resulted in either the release of a drop of reward (~4-8ul of 10% sweetened condensed milk in water v/v) to a lick tube, or the release of air flow (40-ms, 15 psi) to air puff cannula (Ziggy’s Tubes and Wires, 16 gauge) directed to the left and right whisker pads from the rear position. The ViRMEn computer also controlled TTL pulses sent directly to a 532-nm DPSS laser (Shanghai, 200mW).

#### Behavioral shaping

Following post-surgical recovery, over the course of 4-7 days mice were extensively handled while gradually restricting water intake to an allotted volume of 1-2 mL per day. Throughout water-restriction mice were closely monitored to ensure no signs of dehydration were present and that body mass was at least 80% of the pre-restriction value. Mice were then introduced to the VR setup where behavior was shaped to perform the accumulation of evidence task as previously described in detail^26^ (**Supplementary Fig. 5a**) or the permanent cues (control #2) task (**Supplementary Fig. 5f**). We tested a total of 32, 34 and 20 mice in the accumulation of evidence, no distractors (control #1), and permanent cues (control #2) tasks, respectively. No mice received optogenetic testing in all three tasks, but 7 mice received optogenetic testing in the accumulation of evidence and no distractors task, and 19 mice received optogenetic testing in the no distractors and permanent cues tasks (**Supplementary Table 1**).

Shaping followed a similar progression in both tasks. In the first four shaping mazes of both procedures, a visual guide located in the rewarded arm was continuously visible, and the maze stem was gradually extended to a final length of 300-cm (**Supplementary Fig. 5a,f**). In mazes 5-7 of the evidence accumulation shaping procedure (**Supplementary Fig. 5a**), the visual guide was removed and the cue region was gradually decreased to 200-cm, thus introducing the full 100-cm delay region of the testing mazes. The same shift to a 200-cm cue region and 100-cm delay region occurred in mazes 5-6 of the permanent cues shaping procedure, but without removing the visibility of the visual guide (**Supplementary Fig. 5f**). In mazes 8-9 of evidence accumulation shaping, distractor cues were introduced to the non-rewarded maze side with increasing frequency (mean side ratio (s.d.) of rewarded::non-rewarded side cues of 8.3::0.7 to 8.0::1.6 m^−1^). Distractor cues were similarly introduced with increasing frequency in mazes 6-8 of the permanent cues shaping procedure, while the visual guide was removed in maze 7 and 8. In all evidence accumulation shaping mazes (maze 1-9) cues were only made visible when mice were 10-cm from the cue location and remained visible until trial completion. In the final evidence accumulation testing mazes (maze 10 and 11) cues were made transiently visible (200-ms) after first presentation (10-cm from cue location), while the mean side ratio of rewarded::non-rewarded side cues changed from 8.0::1.6 (**Supplementary Fig. 5a**, maze 10) to 7.7::2.3 m^−1^ (**Supplementary Fig. 5a**, maze 11). In contrast, throughout all shaping (maze 1-6) and testing mazes (maze 7-8) of the permanent cues task, cues were visible from the onset of a trial.

The median number of sessions to reach the first evidence accumulation testing maze (maze 10) was 22 sessions, while the mean number of sessions was 23.0 +/- 0.8 (**Supplementary Fig. 5b-c**). Mice typically spent between 2-5 sessions on each shaping maze before progressing to the next, with performance increasing or remaining stable throughout (**Supplementary Fig. 5d-e**; maze 9: 74.1 +/- 9.8 percent correct). The median number of sessions to reach the first permanent cues (control #2) testing maze (maze 7) was 17 sessions, while the mean number of sessions was 18.0 +/-1.5 (**Supplementary Fig. 5g-j**). Mice typically spent between 2-4 sessions on each shaping maze before progressing to the next, with performance increasing or remaining largely stable throughout (**Supplementary Fig. 5g-j**; maze 6: 87.0 +/- 4.3 percent correct).

#### Optogenetic testing mazes

The evidence accumulation task took place in a 330-cm long virtual T-maze with a 30-cm start region (−30 to 0-cm), followed by a 200-cm cue region and finally a 100-cm delay region (**Fig. 2a**, black, *left*). While navigating the cue region of the maze mice were transiently presented with high-contrast visual cues (wall-sized “towers”) on either maze side, which were also paired with a mild air puff (15 p.s.i, 40-ms) to the corresponding whisker pad. The side containing the greater number of cues indicated the future rewarded arm. A left or right choice was determined when mice crossed an x-position threshold > |15-cm|, which was only possible within one of the maze arms (the width of choice arms were +/- 25-cm relative to the center of the maze stem). Mice received reward (~4-8 μL of 10% v/v sweetened condensed milk in drinking water) followed by a 3-s ITI after turning to the correct arm at the end of the maze, while incorrect choices were indicated by a tone followed by a 12-s ITI. In each trial, the position of cues was drawn randomly from a spatial Poisson process with a rate of 8.0 m^−1^ for the rewarded side and 1.6 m^−1^ for the non-rewarded side (**Supplementary Fig. 5a**, maze 10) or 7.3::2.3 m^−1^ (**Supplementary Fig. 5a**, maze 11). Note that only maze 10 data was used for cross-task comparisons of optogenetic effects with permanent cues and no distractors control tasks in order to precisely match cue presentation statistics (**Fig. 2–3**; **Extended Data Fig. 3–6**). Visual cues (and air puffs) were presented when mice were 10-cm away from their drawn location and ended 200-ms (or 40-ms) later. Cue positions on the same side were also constrained by a 12-cm refractory period. Each session began with warm-up trials of a visually-guided maze (**Supplementary Fig. 5a**, maze 4), with mice progressing to the evidence accumulation testing maze after 10 trials (or until accuracy reached 85% correct). During performance of the testing maze if accuracy fell below 55% over a 40-trial running window, mice were transitioned to an easier maze in which cues were presented only on the rewarded side and did not disappear following presentation (**Supplementary Fig. 5a**, maze 7). These “easy blocks” were limited to 10 trials, after which mice returned to the main testing maze regardless of performance. Behavioral sessions lasted for ~1-hour and typically consisted of ~150-200 trials.

All features of the “no distractors” (control #1) task (**Fig. 2b**, magenta, *middle*; **Supplementary Fig. 5a,g**, maze 12) were identical to the evidence accumulation task (**Supplementary Fig. 5a**, maze 10) except: (1) distractor cues were removed from the non-rewarded side, and (2) a distal visual guide located in the rewarded arm was transiently visible during the cue region (0-200-cm).

All features of the “permanent cues” (control #2) task (**Fig. 2b**, cyan, *right*; **Supplementary Fig. 5g**, maze 8) were identical to the evidence accumulation task except: (1) reward and non-reward side visual cues were made permanently visible from trial onset. As in the evidence accumulation task, whisker air puffs were only delivered when mouse position was 10-cm from visual cue location. Note that mice underwent optogenetic testing on two permanent cues mazes (maze 7 and maze 8). Maze 8 shared identical reward to non-reward side cue statistics (8.0::1.6 m^−1^) as maze 10 of the evidence accumulation task. Therefore, for all cross-task comparisons of optogenetic inhibition only data from these mazes were analyzed (**Fig. 2–3**; **Extended Data Fig. 3–6**).

To discourage side biases, in all tasks we used a previously implemented debiasing algorithm^26^. This was achieved by changing the underlying probability of drawing a left or a right trial according to a balanced method described in detail elsewhere^26^. In brief, the probability of drawing a right trial, *p_R_*, is given by:

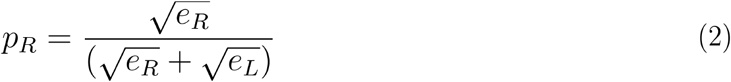

Where *e_R_* (and *e_L_*) are the weighted average of the fraction of errors the mouse has made in the past 40 right (and left) trials. The weighting for this average is based on a half-Gaussian with σ = 20 trials in the past, which ensures that most recent trials have larger weight on the debiasing algorithm. To discourage the generation of sequences of all-right (or all-left) trials, we capped 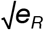 and 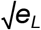 to be within the range [0.15, 0.85]. Because the empirical fraction of drawn right trials could significantly deviate from *p_R_*, particularly when the number of trials is small, we applied an additional pseudo-random drawing prescription to *p_R_*. Specifically, if the empirical fraction of right trials (calculated using a σ = 60 trials half-Gaussian weighting window) is above *p_R_*, right trials were drawn with probability 0.5 *p_R_*, whereas if this fraction is below *p_R_*, right trials were drawn with probability 0.5 (1 + *p_R_*).

#### Virtual corridor

Following shaping in the behavioral tasks above mice were transitioned to free navigation in a virtual corridor arena in the same VR apparatus described above. The virtual corridor was 6-cm in diameter and 330-cm in effective length (**Fig. 1e-f**). This included a start region (−10 to 0-cm), a reward location (310-cm) in which mice received 4 μL of 10% v/v sweetened condensed milk in drinking water, and a teleportation region (320-cm) in which mice were transported back to the start region following a variable ITI with mean of 2-s. Mice were otherwise allowed to freely navigate the virtual corridor over the course of ~70 minute sessions. The virtual environment was controlled by the ViRMEn software engine, with real-to-virtual world movement transformations as described above.

### Optogenetics during VR behavior

According to a previously published protocol^55^, optical fibers (200uM, 0.37 NA) were chemically etched using 48% hydrofluoric acid to achieve tapered tips of lengths 1.5-2 mm (DMS-targeted) or 1-1.5 mm (NAc-targeted). Following behavioral shaping in VR (and >6weeks of viral expression) mice underwent optogenetic testing. On alternate daily sessions, optical fibers in the left or right hemisphere were unilaterally coupled to a 532-nm DPSS laser (Shanghai, 200 mW) via a multi-mode fiberoptic patch cable (PFP, 62.5 μM). On a random subset of trials (10-30%), mice received unilateral laser illumination (5 mW, measured from patch cable) that was restricted to the first passage through 0-200-cm of the virtual corridor (**Fig. 1** and **Extended Data Fig. 2**), or the cue region (0-200cm) of each T-maze decision-making task (**Fig. 3**). The laser was controlled by TTL pulses generated using a National Instruments DAQ card on a computer running the ViRMEn-based virtual environment.

### Conditioned place preference test

Mice underwent a real-time conditioned place preference (CPP) test with bilateral optogenetic inhibition paired to one side of a two-chamber apparatus (**Supplementary Fig. 3**). The CPP apparatus consisted of a rectangular Plexiglass box with two chambers (29-cm x 25-cm) separated by a clear portal in the center. The same grey, plastic flooring was used for both chambers, but each chamber was distinguished by vertical or horizontal black and white bars on the chamber walls. During a baseline test, mice were placed in the central portal while connected to patch cables coupled to an optical commutator (Doric) and were allowed to freely move between both sides for 5 minutes. In a subsequent 20-min test, mice received continuous, bilateral optogenetic inhibition (532-nm, 5-mW) when located in one of the two chamber sides (balanced across groups). Video tracking, TTL triggering, and data analysis were carried out using Ethovision software (Noldus). Mice who displayed a bias for one chamber side greater than 45-s during the baseline test were excluded from analysis.

### Behavior analyses

#### Data selection

See **Supplementary Table 1** for a list of all mice with optogenetic testing data in the virtual corridor, the accumulation of evidence, no distractors, or permanent cues tasks. The following describes the trial selection criteria for inclusion in analyses throughout.

For cross-task comparisons (**Fig. 2–3**; **Extended Data Fig. 3–6**) we analyzed only trials from evidence accumulation maze 10 (**Supplementary Fig. 5a**), “no distractors” maze 12 (**Supplementary Fig. 5a,g**), and “permanent cues” maze 8 (**Supplementary Fig. 5g**), which each followed matching rewarded- and non-rewarded side cue probability statistics (save for the by-design absence of non-rewarded cues in the “no distractors” control task). For model-based analyses of the evidence accumulation task (**Fig. 4–7**, **Extended Data Fig. 7–10**) both maze 10 and maze 11 data were included, which differed only modestly in the side ratio of reward to non-reward side cues (**Supplementary Fig. 5a**, ~50% of trials were maze 10 or 11). In all tasks and all analyses throughout we removed initial warm-up blocks (**Supplementary Fig. 5a**, maze 4, approximately 5-15% of total trials). For model-based analyses of the evidence accumulation task (**Fig. 4–7**, **Extended Data Fig. 7–10**), we included interspersed “easy blocks” capped at 10 trials in length (**Supplementary Fig. 5a**, maze 7, see description above). These trial blocks comprised approximately ~5% of total trials, were included to avoid gaps in trial history, and were treated identical to the main evidence accumulation mazes by the models. These trials were removed from cross-task comparisons of optogenetic inhibition (**Fig. 2–3**; **Extended Data Fig. 3–6**).

For analysis of optogenetic inhibition during virtual corridor navigation (**Fig. 1** and **Extended Data Fig. 2**) we removed trials with excess travel >10% of maze stem (or >330-cm) and mice with <150 total trials from measures of y-velocity, x-position, and average view angle. Trials with excess travel had similar proportions across laser off and laser on trials and pathway-specific inhibition and control groups (indirect pathway: 8.1% of laser off and 8.2% of laser on trials; direct pathway: 8.2% of laser off and 8.1% of laser on trials; no opsin control: 7.0% of laser off and 6.9% laser on trials; exact trial N in figure legends). Excess travel trials reflected the minority of trials in which mice made multiple traversals of the virtual corridor, thus skewing measures of average y-velocity, x-position, and view angle during the majority of “clean” corridor traversals. Importantly, we excluded no trials in direct measurements of distance, per-trial view angle standard deviation, and trials with excess travel in order to detect potential effects of pathway-specific DMS inhibition (or DMS illumination alone) on these measures (**Fig. 1** and **Extended Data Fig. 2**).

Similarly, for all cross-task comparisons (**Fig. 2-3**; **Extended Data Fig. 3–6**) we removed trials with excess travel for all analyses comparing choice, y-velocity, x-position, and average view angle. To better capture task-engaged behavior we also only considered trial blocks in which choice accuracy was greater than >60% for these measures. Excess travel trials were not excluded for cross-task comparisons of effects on measures of distance, per-trial view angle standard deviation, and trials with excess travel. Exact trial and mouse N are reported in figure legends throughout.

For cross-state comparisons of motor output measures (**Extended Data Fig. 10**) we did not exclude trial blocks with choice accuracy <60%, given that GLM-HMM states were differentially associated with performance levels. However, for the reasons outlined above we removed excess travel trials from measures of y-velocity, x-position, and average view angle, and additionally only considered mice who occupied all three GLM-HMM states after trial selection. For measures of per-trial standard deviation in view angle, distance, and excess travel we applied no trial selection criteria, but only mice who occupied all three GLM-HMM states were included for analysis. Exact trial and mouse N are reported in the legend.

#### General performance indicators

Accuracy was defined as the percentage of trials in which mice chose the maze arm corresponding to the side having the greater number of cues (**Fig. 2c**). For measures of choice bias, sensory evidence and choice were defined as either ipsilateral or contralateral relative to the unilaterally-coupled laser hemisphere. Choice bias was calculated separately for laser off and on trials as the difference in choice accuracy between trials where sensory evidence indicated a contralateral reward versus when sensory evidence indicated an ipsilateral reward (% correct, contralateral-ipsilateral, positive values indicate greater contralateral choice bias) (**Fig. 3d,g,j,o**). Delta choice bias was calculated as the difference in contralateral choice bias between laser off and on trials (on-off, positive values indicate laser-induced contralateral choice bias) (**Fig. 3e,g,k,p**). In **Extended Data Fig. 8c-f**, reward at GLM-HMM state transition reflects the total amount of reward (mL) received from the start of the session up to the trial prior to a state transition (mice typically receive 1-1.5 mL per session). Reward rate at GLM-HMM state transition was calculated as the sum of reward received from the start of the session up to the trial prior to a state transition, divided by the sum of all trial durations from the start of the session up to the trial prior to a state transition. GLM-HMM transitions were defined as a within-session change in the most likely state based on the posterior probability of each state (see **GLM-HMM** below for more details).

#### Psychometric curve fitting

Psychometric performance was assessed based on the percentage of contralateral choices as a function of the difference in the number of contralateral and ipsilateral cues (#contra-#ipsi) (**Fig 6c-f**; **Extended Data Fig. 4**). In **Extended Data Fig. 4**, transparent lines reflect the mean performance of individual mice in bins (−16:4:16, #contra-#ipsi) of sensory evidence during laser off (black) and on (green) trials, while bold lines reflect the corresponding mean and s.e.m across mice. In **Fig. 6**, psychometric curves were fit to the following 4-parameter sigmoid using maximum likelihood fitting^56^:

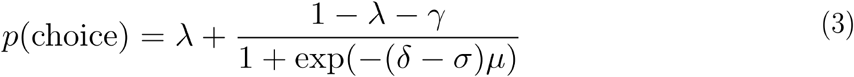

where λ and *γ* are the right and left lapse rates, respectively, *σ* is the offset, *μ* is the slope, and *δ* is the difference in the number of contralateral and ipsilateral cues on a given trial. Each point in **Fig. 6c-f** represents the binned difference in cues in increments of 4 from −16 to 16 (as in **Extended Data Fig. 4**), from which we calculated the percentage of contralateral choice trials for each bin.

#### Motor performance indicators

Y-velocity (cm/s) was calculated on every sampling iteration (120 Hz, or every ~8ms) of the ball motion sensor as dY/dt where *dY* was the change in Y-position displacement in VR and *dt* was the elapsed time from the previous sampling of the sensor. The y-velocity for all iterations in which a mouse occupied y-positions from 0-300-cm in 25-cm bins were then averaged across iterations in each bin to obtain per-trial y-velocity as a function of y-position. Binned y-velocity as a function of y-position was then averaged across trials for individual mice, and the average and standard error of the mean across mice reported throughout (**Fig. 1g**; **Fig. 2d**; **Extended Data Fig. 6a-c**; **Extended Data Fig. 10b,p,i,w**; averaged across y-position 0-200cm in **Extended Data Fig. 2b** and **Extended Data Fig. 3b**).

X-position trajectory (cm) as a function of y-position was calculated per trial by first taking the x-position at y-positions 0-300cm in 1-cm steps, which was defined as the x-position at the last sampling time t in which y(t) ≥ Y, and then averaging across y-position bins of 25-cm from 0 to 300cm. Binned x-position as a function of y-position was then averaged across left/right (or ipsilateral/contralateral) choice trials for individual mice, and the average and standard error of the mean across mice was reported throughout (**Fig. 1h**; **Fig. 2e**; and **Extended Data Fig. 10c,q**; averaged across y-positions 0-200cm in **Extended Data Fig. 2c**; **Extended Data Fig. 3c**; **Extended Data Fig. 6j-l**; and **Extended Data Fig. 10j,x**). Average view angle trajectory (degrees) was calculated in the same manner as x-position (**Fig. 1i**; **Fig. 2f**; and **Extended Data Fig. 10d,r**; average across y-positions 0-200cm in **Extended Data Fig. 2d**; **Extended Data Fig. 3d**; **Extended Data Fig. 6j-l**; and **Extended Data Fig. 10k,y**). View angle standard deviation was calculated by first sampling the per-trial view angle from 0-300cm of the maze in 5-cm steps. The standard deviation in view angle was then calculated for each trial, and then averaged across trials for individual mice. The average and standard error of the mean across mice are reported throughout (**Extended Data Fig. 2e**; **Extended Data Fig. 3f**; **Extended Data Fig. 6g-i**; and **Extended Data Fig. 10e,s,l,z**). This measure sought to capture unusually large deviations in single trial view angles, which would be indicative of excessive turning or rotations.

Distance traveled was measured per trial as the sum of the total x and y displacement calculated at each sampling iteration t, as 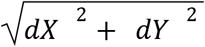. Distance traveled per trial was then averaged across trials for individual mice and the average and standard error of the mean across mice was reported throughout (**Fig. 1j**; **Fig. 2g**; **Extended Data Fig. 2f**; **Extended Data Fig. 6d-f**, *left*; **Extended Data Fig. 10f,t,m,aa**). Excess travel was defined as the fraction of trials with total distance traveled per trial (calculated as above) greater than 10% of maze length (or >330cm). The average and standard error of the mean across mice was reported throughout (**Extended Data Fig. 2g**; **Extended Data Fig. 3e**; **Extended Data Fig. 6d-f**, *right*; and **Extended Data Fig. 10g,u,n,bb**).

Decoding of choice based on the trial-by-trial x-position (**Extended Data Fig. 3g,i**) or view angle (**Extended Data Fig. 3h,j**) of mice was carried out by performing a binomial logistic regression using the MATLAB function *glmfit*. In **Extended Data Fig. 3g-h**, the logistic regression was fit separately for individual mice at successive y-positions in each T-maze stem (0-300cm in 25-cm bins), where the trial-by-trial average x-position (or view angle) at each y-position bin (calculated as above) was used to generate weights predicting the probability of a left or right choice given a particular x-position (or view angle) value. Individual mouse fits were weighted according to the proportion of left and right choice trials. 5-fold cross-validation (re-sampled for new folds 10 times) was used to evaluate prediction accuracy on held-out trials. A choice probability greater than or equal to 0.5 was decoded as a right choice, and prediction accuracy for individual mice was calculated as the fraction of predicted choices matching actual mouse choice, averaged across cross-validation sets. The same approach was used in **Extended Data Fig. 3i-j**, except that the training data was randomly sampled across all mice from a single task (50% of total trials, re-sampled 50 times; train on data from evidence accumulation, no distractors, or permanent cues task). The learned weights were then used to predict choice based on held out x-position (or view angle) data from all three tasks, with prediction accuracy calculated as the fraction of predicted choices matching actual choice, parsed by individual mice, and averaged across cross-validation sets. A package of code for behavioral analysis in VR-based T-maze settings is available at: https://github.com/BrainCOGS/behavioralAnalysis. In addition, all analyses described here can be replicated at https://github.com/ssbolkan/BolkanStoneEtAl.

### General statistics and Reproducibility

We performed one-way ANOVAs to assess effects of the factor task (three levels: evidence accumulation, no distractors, or permanent cue) on choice accuracy (**Fig. 2c**), distance traveled (**Fig. 1g**), average y-velocity (0-200cm) (**Extended Data Fig. 3b**), average x-position (0-200cm) on left or right choice trials (**Extended Data Fig 6c**), average view angle (0-200cm) on left or right choice trials (**Extended Data Fig. 3d**), percent trials with excess travel (**Extended Data Fig. 3e**), per-trial standard deviation in view angle (**Extended Data Fig. 3f**), delta (laser on-off) choice bias (**Extended Data Fig. 5b-d**), delta (laser on-off) distance traveled (**Extended Data Fig. 6d-f**, *left*), delta (laser on-off) percent trials with excess travel (**Extended Data Fig. 6d-f**, *right*), delta (laser on-off) per-trial standard deviation in view angle (**Extended Data Fig. 6g-i**), delta (laser on-off) average x-position (0-200cm) (**Extended Data Fig. 6j-l**, *left*), and delta (laser on-off) average view angle (0-200cm) (**Extended Data Fig. 6j-l**, *right*). Post-hoc comparisons between tasks were made when a main effect of the factor task had a p-value less than alpha <0.05/2 to account for multiple group comparisons (**Extended Data Fig. 5b-d**). One exception to this is **Extended Data Fig. 6j-l**, where all post-hoc comparisons were made for laser effects on delta (on-off) x-position and view angle (and displayed with corresponding exact p-values) to provide greater clarity around trend-level effects. We did not assume normality of the data in all post-hoc comparisons, which used the non-parametric, unpaired, two-tailed Wilcoxon rank sum test. A p-value below 0.025 was considered significant in order to correct for multiple comparisons (p = 0.05/2 comparisons per group). Exact p-values, degrees of freedom, and z-statistics are reported in the text and/or legends.

We performed one-way ANOVAs to assess effects of the factor group (three levels: indirect pathway inhibition, direct pathway inhibition, or no opsin illumination) on delta (laser on-off) y-velocity (**Extended Data Fig. 2b**), delta (laser on-off) x-position (**Extended Data Fig. 2c**), delta (laser on-off) view angle (**Extended Data Fig. 2d**), delta (laser on-off) per-trial standard deviation in view angle (**Extended Data Fig. 2e**), delta (laser on-off) distance (**Extended Data Fig. 2f**), delta (laser on-off), and delta (laser on-off) percent trials with excess travel (**Extended Data Fig. 2g**), and delta (laser on-off) preference (i.e. time) and speed during the real-time conditioned place preference test (**Supplementary Fig. 3**).

We performed a repeated-measure one-way ANOVA to assess effects of the within-subject factor state (three levels: GLM-HMM state 1, 2, or 3) on within-session accumulated reward or reward rate prior to GLM-HMM transition (**Extended Data Fig. 8c-f**), per-trial standard deviation in view angle (**Extended Data Fig. 10e,s**), distance traveled (**Extended Data Fig. 10f,t**), percent trials with excess travel (**Extended Data Fig. 10g,u**), delta (laser on-off) average x-position 0-200cm (**Extended Data Fig. 10j,x**), delta (laser on-off) average view angle 0-200cm (**Extended Data Fig. 10k,y**), delta (laser on-off) per-trial standard deviation in view angle (**Extended Data Fig. 10l,z**), delta (laser on-off) distance traveled (**Extended Data Fig. 10m,aa**), and delta (laser on-off) percent trials with excess travel (**Extended Data Fig. 10n,bb**). Post-hoc comparisons between groups were made when a main effect of the factor task had a p-value <0.05. One exception is in **Extended Data Fig. 6j,k,x,y**, where all post-hoc comparisons were made for laser effects on delta (on-off) x-position and view angle (and displayed with corresponding exact p-values) to provide greater clarity around trend-level effects. We did not assume normality of the data in all post-hoc comparisons, which used the non-parametric, unpaired, two-tailed Wilcoxon rank sum test. A p-value below 0.025 was considered significant in order to correct for multiple comparisons (p = 0.05/2 comparisons per group). Exact p-values, degrees of freedom, and z-statistics are reported in the text and/or legends.

In **Fig. 3e,h,k,p**, we used the non-parametric, unpaired, two-tailed Wilcoxon rank sum test to assess effects of indirect or direct pathway inhibition versus no opsin illumination of brain tissue on delta (laser on-off) choice bias. A p-value below 0.025 was considered significant in order to correct for multiple comparisons (p = 0.05/2 comparisons per group). Exact p-values, degrees of freedom, and z-statistics are reported in the text and/or legends.

Related to **Fig. 1a-c**, **Extended Data Fig. 1**, and **Supplementary Fig. 2**, we used a paired, two-tailed Wilcoxon signrank test on cross-trial average firing rates (baseline pre versus laser on or laser on versus laser post) to determine significance of laser modulation of single neuron activity. A bonferroni-corrected p-value was used to determine significance (p < 0.00083 for 60 indirect pathway neurons or p < 0.001 for 50 direct pathway neurons).

All experiments, but not analysis, were conducted blind to experimental conditions. All experiments were conducted over multiple cohorts (see **Supplementary Table 1** for details of individual mouse testing across experiments). Individual cohorts consisted of a random selection of test groups (e.g. DMS indirect and direct pathway inhibition, and DMS no-opsin mice). We did not account for cohort effects in our statistical analyses, but no obvious cohort-dependent effects were qualitatively observable. No statistical method was used to predetermine group sample sizes; rather animal and trial N were targeted to match or exceed similar studies.

### Bernoulli GLM

#### Coding of covariates and choice output

We coded the external covariates (referred to as inputs in **Fig. 4b**) and output (the mouse’s choice) on each trial as follows:

Δ cues: an integer value from −16 to 16, divided by the standard deviation of the Δ cues across all sessions in all mice, representing the standardized difference between the number of cues on the right and left sides of the maze.
Laser: a value of 1,-1, or 0 depending on whether optogenetic inhibition was on the right hemisphere, left hemisphere, or off, respectively.
Previous choice: a value of 1or −1 if the choice on a previous trial was to the right or left, respectively. We set the value to 0 at the start of each session when there was an absence of previous choices (e.g. for the third trial of a session, previous choices 3-6 would be coded as 0).
Previous rewarded choice: a value of 1, −1, or 0 depending on whether the previous choice was correct and to the right, correct and to the left, or incorrect, respectively.
Choice output: a value of 1 or 0 depending on whether the mouse turned right or left.

#### Fitting

We used a Bernoulli generalized linear model (GLM), also known as logistic regression, to model the binary (right/left) choices of mice as a function of task covariates. This also corresponds to a 1-state GLM-HMM (**Fig. 4**; **Extended Data Fig. 7a**). The model was parameterized by a weight vector (carrying weights for sensory evidence, choice and reward history, and DMS inhibition). On each trial *t*, the weights map the external covariates to the probability of each choice *y_t_*. The model can be written:

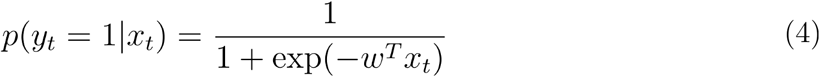

We then fit the model by penalized maximum likelihood, which involved minimizing the negative log-likelihood function plus a squared penalty term on the model weights. The log-likelihood function is given by the conditional probability of the choice data **Y** = *y*_1_, … *y_T_* given all the external covariates **X** = *x*_1_, … *x_T_*, considered as a function of the model parameters:

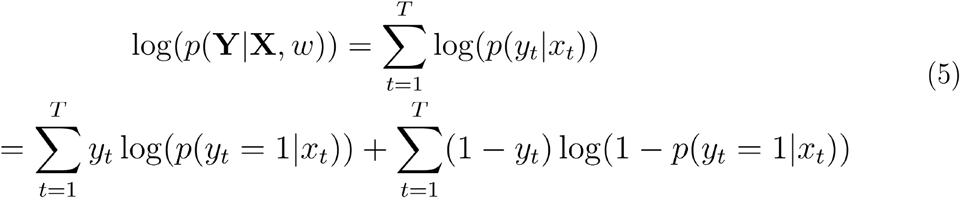

We then minimized the loss function, given by 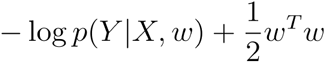, using python’s scipy.optimize.minimize. This can be interpreted as a log-posterior over the weights, with 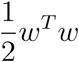 representing the negative log of a Gaussian prior distribution with mean zero and variance, which regularizes by penalizing large weight values. We computed the posterior standard deviation of the fitted GLM weights (shown as error bars in **Fig. 4c-d**) by taking the diagonal elements of the inverse negative Hessian (matrix of second derivatives) of the log-likelihood at its maximum^57,58^.

### GLM-HMM

#### Model architecture

To incorporate discrete internal states, we used a hidden Markov model (HMM) with a Bernoulli GLM governing the decision-making behavior in each state. The model is defined by a transition matrix and a vector of GLM weights for each state. The transition matrix contains a fixed set of probabilities that govern the probability of changing from a state *z* ∈ {1,… *K*} on trial *t* to any other state on the next trial. We refer to these as transition probabilities, which can be abbreviated as follows:

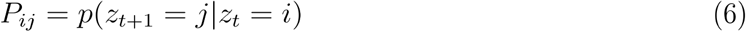

Each GLM has a unique set of weights *w_k_* that map the external covariates *x_t_* (coded as described in the section **Bernoulli GLM**) to the probability of the choice *y_t_* for each of the *k* states. These probabilities can be expressed as a modified version of equation 4

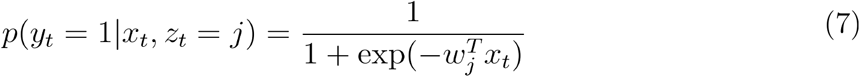

where now the choice probability on each trial is dependent on both the external covariates (inputs) and the state on that trial and is determined by state-dependent GLM weights^30–32^. We refer to these as observation probabilities, which can be abbreviated as follows:

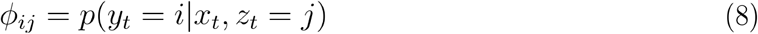

#### Fitting

We fit the GLM-HMM to the data using the expectation-maximization (EM) algorithm^31^. The EM algorithm computes the maximum likelihood estimate of the model parameters using an iterative procedure that involves an E-step (expectation), in which the posterior distribution of the latent variables is calculated, followed by an M-step (maximization), in which the values of the model parameters are updated given the posterior distribution of the latents. These steps are repeated until the log-likelihood of the model converges on a local optimum^58^.

The log-likelihood (also referred to as the log marginal likelihood) is obtained from the joint probability distribution over the latent states **Z** = *z*_1_, … *z_T_* and the observations **Y** = *y*_1_, …*y_T_* on each trial given the model parameters *θ*. Marginalizing over the latents, the log-likelihood is computed as the log of the sum over states of the marginal probabilities and is written:

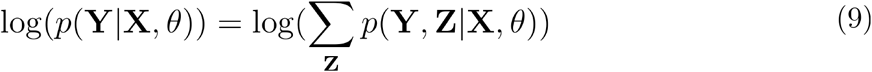

The set of parameters *θ* governing the model consists of a transition matrix and the state-dependent GLM weights, which we described above. We initialized the transition matrix by sampling from a Dirichlet distribution with a larger concentration parameter over the diagonal entries (*α_ii_* = 5, *α_ij_* = 1), reflecting the fact that the probability of staying in the same state from one trial to the next should be larger than the probability of transitioning to a different state. For the GLM weights, we reasoned that the true values for each state would likely be in approximately the same range as the true values for the one-state (GLM) case. Therefore, we initialized the per-state GLM weights *w_k_* with *k* ∈ {1, …, *K*} by first fitting a basic GLM (see **Bernoulli GLM**) to find *w*_0_. Then, since we didn’t want the initial weights to be the same in each state, we initialized *w_k_* = *w*_0_ + *ϵ_k_* where 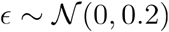.

The goal of the E-step of the EM algorithm is to compute *p*(**Z|X, Y**, *θ*), the posterior probability of the latent states given the observations and the model parameters. This can be obtained using a two-stage message passing algorithm known as the forward-backward algorithm or the Baum-Welch algorithm^31^. The forward pass, sometimes called “filtering,” finds the normalized conditional probability 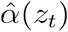 for each state *z* at trial *t* by iteratively computing

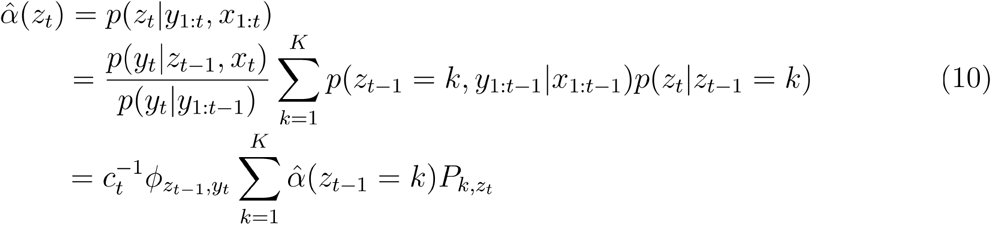

where *c_t_* = *p*(*y_t_*|*y*_1:*t*–1_) is a scale factor, 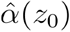, the prior probability over states before any data are observed, is given as a uniform distribution over states, and *K* is the total number of states. Note that this is a normalized version of the algorithm that avoids underflow errors (see Bishop Chapter 13)^58^.

The backward step, also referred to as “smoothing,” takes the information from the forward pass and works in the reverse direction, carrying the information about future states backwards in time to further refine the latent state probabilities. Here we find the normalized conditional probability 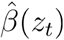 for each state *z* at trial *t* by iteratively computing

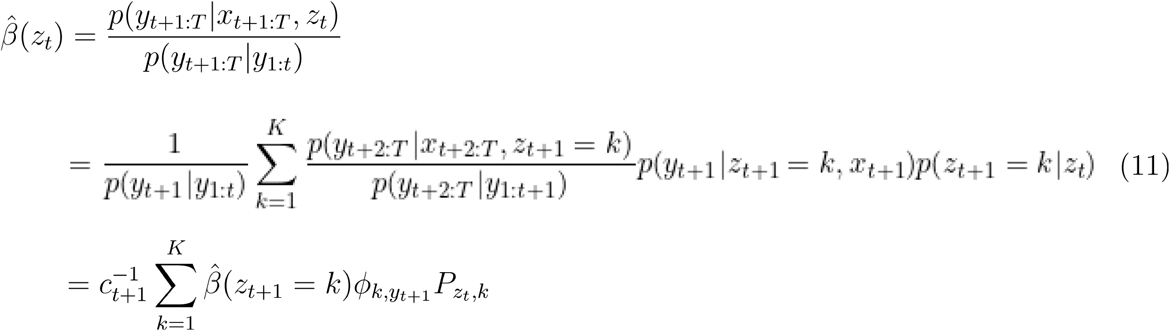

where *β*(*z_T_*) = 1.

From these two conditional probabilities we can calculate the marginal posterior probabilities of the latent states:

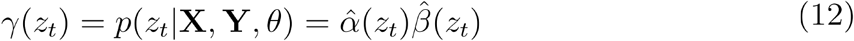

which was the goal of the E-step. We can also compute the joint posterior distribution of two successive latents:

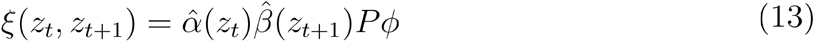

Which will be important for computing updates in the M-step. Because the format of the data included sessions from several different mice over many days, we computed the forward-backward pass separately for each session. This ensured that the learned transition probabilities would not take into account the effect of the last trial of one session on the first trial of the next session.

The M-step of the EM algorithm takes the newly computed posterior probabilities of the latents and uses them to update the values of the model parameters (equations 6–8) by maximizing the expression for *P* and *w*. Since the transition probabilities are fixed, we can compute their updates using the closed form solution:

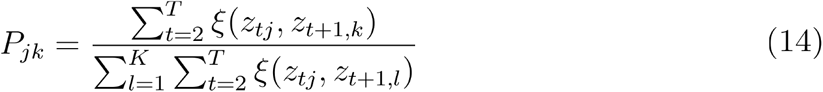

This closed-form update can be derived by applying the appropriate Lagrange multipliers to the complete-data log-likelihood function^58^.

Maximization for *w* involves minimizing the negative of the log-likelihood function, weighted by the marginal posterior probabilities of the latent states, plus a squared penalty term on the model weights. This penalty can be interpreted as the negative log of a Gaussian prior with mean zero and variance 1, which regularizes by penalizing large weight values. The resulting loss function is

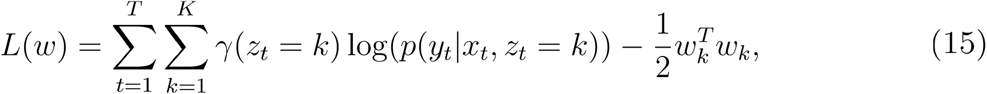

which we optimized using numerical optimization and the L-BFGS-B algorithm as previously described (see Bernoulli GLM).

Both E and M steps of the EM algorithm are guaranteed to increase the log-likelihood. We alternated E and M steps until the difference between the log-likelihoods over ten iterations was smaller than a given tolerance (tol = 1e-3). Because the EM algorithm only guarantees that the log-likelihood will converge upon a local optimum^58^, we fit the model 20 times using different initializations of the weights and transition matrix and verified that the top four or more fits all converged on the same solution (meaning that the weights for each fit were the same within a tolerance of +/- 0.05) in order to confirm that the algorithm had indeed found the global optimum. After determining the best fit, we computed the posterior standard deviation of the fitted GLM weights (shown as error bars in **Fig. 6a-b** and **Extended Data Fig. 7d-f**) by taking the inverse Hessian of the optimized log-likelihood.

#### Model selection

In **Extended Data Fig. 7a**, we performed cross-validation on the data from both the indirect and direct pathway inhibition groups. To obtain a test set, we selected ~20% of sessions from the data to hold out from model fitting. Test sessions were chosen by randomly selecting an approximately equal number of sessions from each of the 13 mice in either group. Constraining the held-out data in this way ensured that the cross-validation results were not affected by possible individual differences across mice. We then calculated the log-likelihood of the test data after fitting the model under parameterizations of 1-5 states to the remaining ~80% of sessions. We express the log-likelihood in units of bits per session (bps), defined:

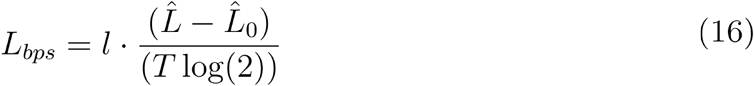

Where *l* is the average session length, *T* is the number of trials in the test set, and 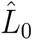 is the log-likelihood of the test set data under the bias-only Bernoulli GLM. To obtain the bias term *b* we computed:

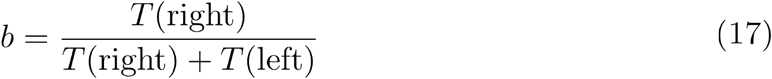

Where *T*(side) is the number of trials in the test set in which the mice turned in that direction. For all cross-validation results presented in the paper, we report the averaged *L_bps_* from five different test sets. We followed the same procedure as above in **Extended Data Fig. 7b**, selecting the optimal number of previous choices using a 3-state GLM-HMM under parameterizations of 1-8 previous choices while holding the number of all other external inputs (Δ cues, laser, bias, and previous rewarded choice) constant.

#### Testing

In **Fig. 5c**, we compared the performance of the GLM-HMM to the GLM by calculating the log-likelihood of the test sets of individual mice. To do so, we held out data and fitted the model across all animals using the same procedure described above. However, we then split the test set by mouse (thus creating 13 different test sets) and calculated the log-likelihood for each individual animal, thus expressing the log-likelihood in units of mouse bits per session (mbps):

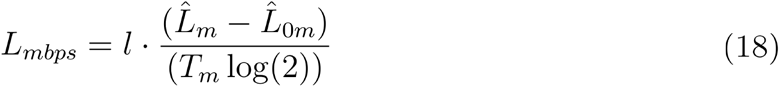

Here, 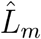 is the optimized log-likelihood of the model in question (either the GLM or 3-state GLM-HMM) for a single mouse. Similarly, 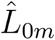 is the optimized log-likelihood under the bias-only Bernoulli GLM and *T_m_* is the total number of trials for that mouse. We then repeated the procedure for five test sets and took the average of the results for each mouse.

In **Fig. 5d**, we evaluated the prediction accuracy of the GLM for each animal by taking the same training and test sets that we used to find the log-likelihoods and using equation 2 to calculate the probability of turning right on each trial. We then compared this probability to the mouse’s actual choice on that trial, labelling the trial as correct if the model predicted a 50% or greater probability of turning in the direction of the mouse’s true choice. We then calculated the prediction accuracy for each mouse as the number of correct trials divided by the total number of trials for that mouse. To evaluate the prediction accuracy of the GLM-HMM for each animal, we computed *p*(*y_t_*|*x*_1:*t*–1_, *y*_1:*t*–1_), or the predictive distribution for trial *t* of the test set using the observations from trials 1 to *t* – 1. This arises from averaging over the state probabilities given previous choice data to get a prediction for a particular trial. That is, we ran the forward pass (see Fitting) to obtain the state probabilities *p*(*z_t_*|*x*_1:*t*–1_, *y*_1:*t*–1_), computed the initial choice probabilities *p*(*y_t_*|*x_t_*, *y_t_*) using equations 7 & 8, and then calculated the predictive distribution as follows:

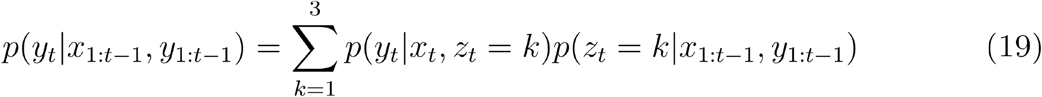

We then ran this forward over all the trials in the test set for each mouse. Finally, we computed the prediction accuracy using the same method described for the GLM prediction accuracy.

#### State assignments

To determine the most likely state on each trial (**Fig. 6c-i**; **Fig. 7g,j**; **Extended Data Fig. 7g-h**; **Extended Data Fig. 8, 10**; and **Supplementary Fig. 4**), we assigned each trial to the maximum posterior probability over states given the inputs and choice data:

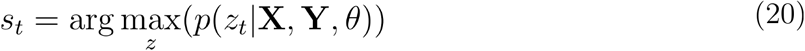

#### Simulating data

For the analyses in **Fig. 6e-f**, **Extended Data Fig. 7**, and **Extended Data Fig. 9**, we evaluated the ability of the 3-state GLM-HMM to predict choices and state transitions that matched the animals’ actual behavior in each state. For the covariates for the simulation, we kept the evidence (Δ cues) and optogenetic inhibition from the real data, but populated the trial history covariates using simulated previous choices. To simulate choices on each trial, we first computed the observation probabilities (equations 7 & 8) using 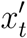 (the external covariates) and *w_k_* (the learned weights from the model fitted to real data). The state *k* on each trial was randomly chosen from a distribution given by the learned transition probabilities *P*_*z*_*t*+1_, *z_t_*_ from the model fitted to real data. We then randomly generated choices 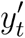 from the distribution of observation probabilities. Repeating this process for each trial to obtain 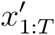 and 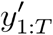 we fitted the model to the simulated data using the same procedure described previously (see *Fitting*) to obtain the posterior probability over states. For **Fig. 6e-f** and **Extended Data Fig. 7**, we computed the psychometric curves for each state using these posterior probabilities and the simulated choices (see *Psychometric curve fitting*).

#### Model comparisons

For the two alternative model comparisons with restricted transition probabilities (**Fig. 7k-m**), we fit the 3-state GLM-HMM using the same general procedure as described above. However, in the case where we disallowed transitions during a session, (**Fig. 7k**), the transition matrix was fixed to the identity matrix and we only fit the state-dependent GLM weights. In the case where we disallowed transitions in and out of state 2 (**Fig. 7l**), we derived a constrained M step that forced the transition probabilities for state 2 to 0. In detail, the constrained M step involved zeroing out the transition probabilities associated with state 2 and then renormalizing so the rows of the transition matrix summed to 1. Note that the three sessions that appear to still allow transitions in and/or out of state 2 for mice inhibited in the direct pathway of the DMS (**Fig. 7l**, right) were due to rare cases where the model had high uncertainty about the state, and the most probable state flipped between state 2 and another state at some point during the session. In **Fig. 7m**, solid curves denote the average log-likelihood for five different test sets. Held-out data for test sets was selected as a random 20% of sessions, using the same number of sessions for each mouse.

### Fluorescent *in situ* hybridization and stereological quantification

*In situ* hybridization (**Supplementary Fig. 1**) was performed using the RNAscope Multiplex Fluorescent Assay (ACD, No. 323110) with the following probes: Mm-Drd1a (406491), Mm-Drd2-C2 (406501-C2, 1:50 dilution in C1 solution), and Cre-01-C3 (474001-C3, 1:50 dilution in C1 solution). Likely due to lower expression of Cre mRNA in D1R-Cre and A2a-Cre mice we did not detect unambiguous Cre fluorescent signal in these lines. We therefore relied on Cre-dependent viral expression of AAV5-DIO-EYFP to report Cre^+^ neurons alongside Drd1a and Drd2 probes in sections from 2 D1R-Cre and 2 A2R-Cre mice, but used all three probes in sections from 2 D2R-Cre mice. In D1R-Cre and A2R-Cre mice the Drd1a and Drd2 probes were fluorescently linked to TSA Plus Cy-3 and TSA Plus Cy-5, respectively (Perkin Elmer). In D2R-cre mice, Drd1a, Drd2, and Cre probes were linked to TSA Plus Cy-3, TSA Plus Fluorescein, or TSA Plus Cy-5, respectively. All fluorophores were reconstituted in DMSO according to Perkin Elmer instructions and diluted 1:1200 in TSA buffer included in the RNAscope kit. Post *in situ* hybridization slides were cover-slipped using Fluoromount-G containing DAPI (SouthernBiotech). We then obtained 20x confocal z-stacks from the DMS, NAcCore, and NAcShell in all lines and manually quantified specificity, penetrance, and D1R^+^/D2R^+^ overlap using LASX software (Leica). Specificity was determined as the percentage of the following: GFP^+^ neurons co-expressing Drd1 in D1R-Cre mice (DMS, n = 5 sections, 193 cells; NAcCore, n = 5 sections, 298 cells; NAcShell, n = 5 sections, 363 cells), GFP^+^ neurons co-expressing Drd2 in A2A-Cre mice (DMS, n = 4 sections, 144 cells; NAcCore, n = 4 sections, 326 cells; NAcShell, n = 4 sections, 312 cells), or Cre^+^ neurons co-expressing Drd2 in D2R-Cre mice (DMS, n = 5 sections, 1,302 cells; NAcCore, n = 5 sections, 1,104 cells; NAcShell, n = 5 sections, 1,187 cells). Penetrance was determined as the percentage of Drd2^+^ neurons co-expressing Cre in D2R-Cre mice (DMS, n = 5 sections, 1,269 cells; NAcCore, n = 5 sections, 1,055 cells; NAcShell, n = 5 sections, 1,144 cells). We did not assess penetrance in D1R-Cre or A2a-Cre lines because our Cre-dependent viral reporter did not fully penetrate all Cre^+^ neurons. Quantification of D1R^+^/D2R^+^ overlap in striatal regions was carried out on 2 D2R-Cre mice and/or 2 D1R-tdTomato mice and measured as both the percentage of D1R^+^ neurons that were D2R^+^ (DMS, n = 10 sections, 2,423 cells; NAcCore, n = 10 sections, 2,196 cells; NAcShell, n = 10 sections, 2,220 cells) and the percentage of D2R^+^ neurons that were D1R^+^ (DMS, n = 5 sections, 868 cells; NAcCore, n = 5 sections, 834 cells; NAcShell, n = 5 sections, 874 cells).

### Histology

Mice were anesthetized with a 0.05 mL injection of Euthasol (i.p.) and transcardially perfused with 1x phosphate-buffered saline (PBS), followed by fixation with 4% paraformaldehyde (PFA). Whole brains with intact fiberoptic implants were post-fixed in 4% PFA for 1-3 days, followed by brain dissection and another 24 hours of post-fixation in PFA. For optogenetic experiments, brains were then transferred to PBS for coronal sectioning (50 μM) on a vibratome. Viral expression and fiberoptic placements were assessed under slide-scanning (NanoZoomer, Hamamatsu) or single slide (Leica) epifluorescent microscopes (**Supplementary Fig. 6**). For FISH experiments, post-fixation dissected brains were transferred through a sucrose gradient: 10% sucrose in PBS for 6-8 hours, 20% sucrose in PBS overnight, and 30% sucrose in PBS overnight. Coronal sections (18 μM) containing the DMS and NAc were made using a cryostat, mounted uncoverslipped on Superfrost plus slides (Fisher), and stored at −80° prior to the FISH protocol. After the FISH protocol, tile-scanning and cellular resolution images of cover-slipped slides were acquired using a confocal microscope (Leica TCS SP8).

## Data Availability

The data that support the findings of this study is publicly available on figshare at the following public Digital Object Identifier: https://doi.org/10.6084/m9.figshare.17299142.v1.

## Code Availability

Code for general use applications of GLM-HMM analyses developed in this study, including all applications to the present data set, are available on GitHub at the following public repository: https://github.com/irisstone/glmhmm. Code to analyze data and regenerate all other plots in this study is publicly available at https://github.com/ssbolkan/BolkanStoneEtAl.

## EXTENDED DATA FIGURES

**Extended Data Fig. 1.**
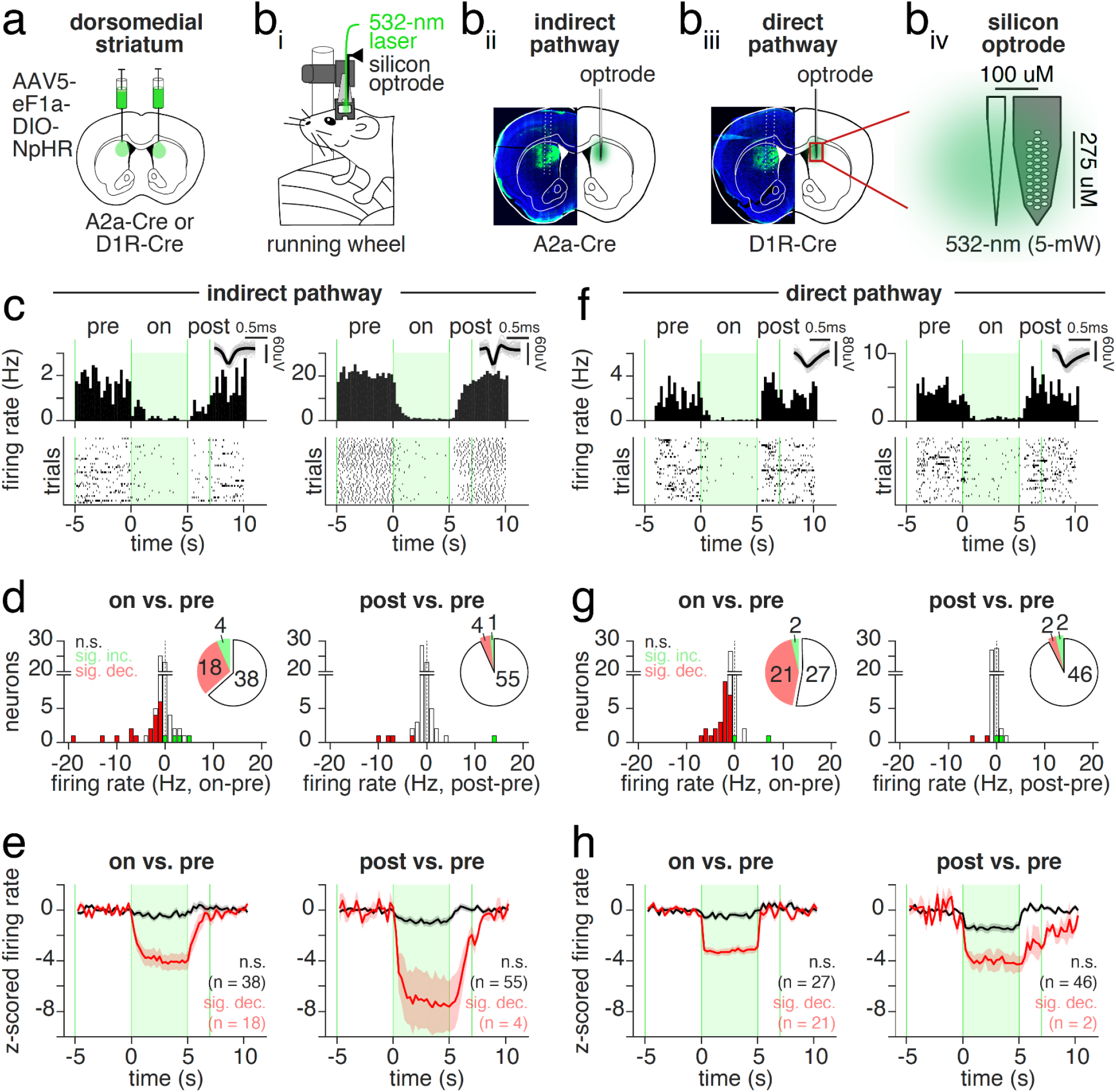
Optogenetic inhibition of DMS pathways is effective, generating little post-inhibitory rebound, nor excitation during the inhibition period. (**a**) Schematic of viral delivery of AAV5-eF1a-DIO-NpHR to the dorsomedial striatum (DMS) of A2a-Cre or D1R-Cre mice. (**b,i**) Schematic of electrophysiological recording and laser delivery (532-nm, 5-mW) to the DMS in awake, head-fixed mice ambulating on a running wheel. (**b,ii**) Example recording electrode tracks and cre-dependent NpHR expression in an A2a-Cre mouse targeting the indirect pathway of the DMS. (**b,iii**) As in **b,ii** but in a D1R-Cre mouse targeting the DMS direct pathway. (**b,iv**) Schematic of silicon optrode recording tip, including tapered optical fiber coupled to a 32-channel silicon probe. (**c**) Two example peristimulus time histograms (PSTH) (*top*) and raster plots of trial-by-trial spike times (*bottom*) from single neurons recorded from the DMS of an A2a-Cre mouse. Inset at top displays average spike waveform (black) and 100 randomly sampled spike waveforms (grey) for each neuron. A trial consisted of 5-s without laser (*pre*, −5 to 0-s), 5-s of 532-nm light (5-mW) delivery (*on*, 0 to 5-s), followed by a 10-s ITI (40 trials per recording site). The first 2-s following laser offset (*post*, 5-7-s) was used to assess post-inhibitory effects. (**d**) *Left*: Histogram of change in average firing rate (*on-pre*, Hz) for all neurons (n = 60) recorded from the DMS of A2a-Cre mice (n = 3). Colors indicate non-significant (black, n = 38 neurons), significantly decreased (red, n = 18 neurons) or increased (green, n = 4 neurons) changes in firing rate determined via paired, two-tailed signrank comparison of average across-trial baseline (pre) or laser (*on*) firing rates. A Bonferroni-corrected significance threshold was used to account for multiple neuron comparisons (p < 0.00083, or p = 0.05/60 neuron comparisons). *Right*: same as *left* but for change in firing rate (*post-pre*, Hz): non-significant (n = 55 neurons), significantly decreased (n = 4) or increased (n = 1). Insets display pie-chart summaries of the proportion of non-significant (black unfilled), significantly decreased (red) or increased (green) neurons. (**e**) *Left*: Mean +/- SEM z-scored firing rate across all non-significantly modulated *on* vs *pre* (black, n = 38) or significantly decreased *on* vs *pre* (red, n = 18) neurons recorded from A2a-Cre mice. *Right*: same as *left* but for all non-significantly modulated *post* vs *pre* (black, n = 55) or significantly decreased *post* vs *pre* (red, n = 4) neurons. (**f**) Same as **c** but for example neurons recorded from the DMS of D1R-Cre mice. (**g**) Same as **d** but for all neurons (n = 50) recorded from the DMS of D1R-Cre mice (n = 2). *Left* (on-pre): non-significant (n = 27), significantly decreased (n = 21), or increased (n = 2). *Right* (post-pre): non-significant (n = 46), significantly decreased (n = 2) or increased (n = 2). A Bonferroni-corrected significance threshold was used to account for multiple neuron comparisons (p < 0.001, or p = 0.05/50 neuron comparisons). (**h**) same as **e** but for neurons recorded from the DMS of D1R-Cre mice.

**Extended Data Fig. 2.**
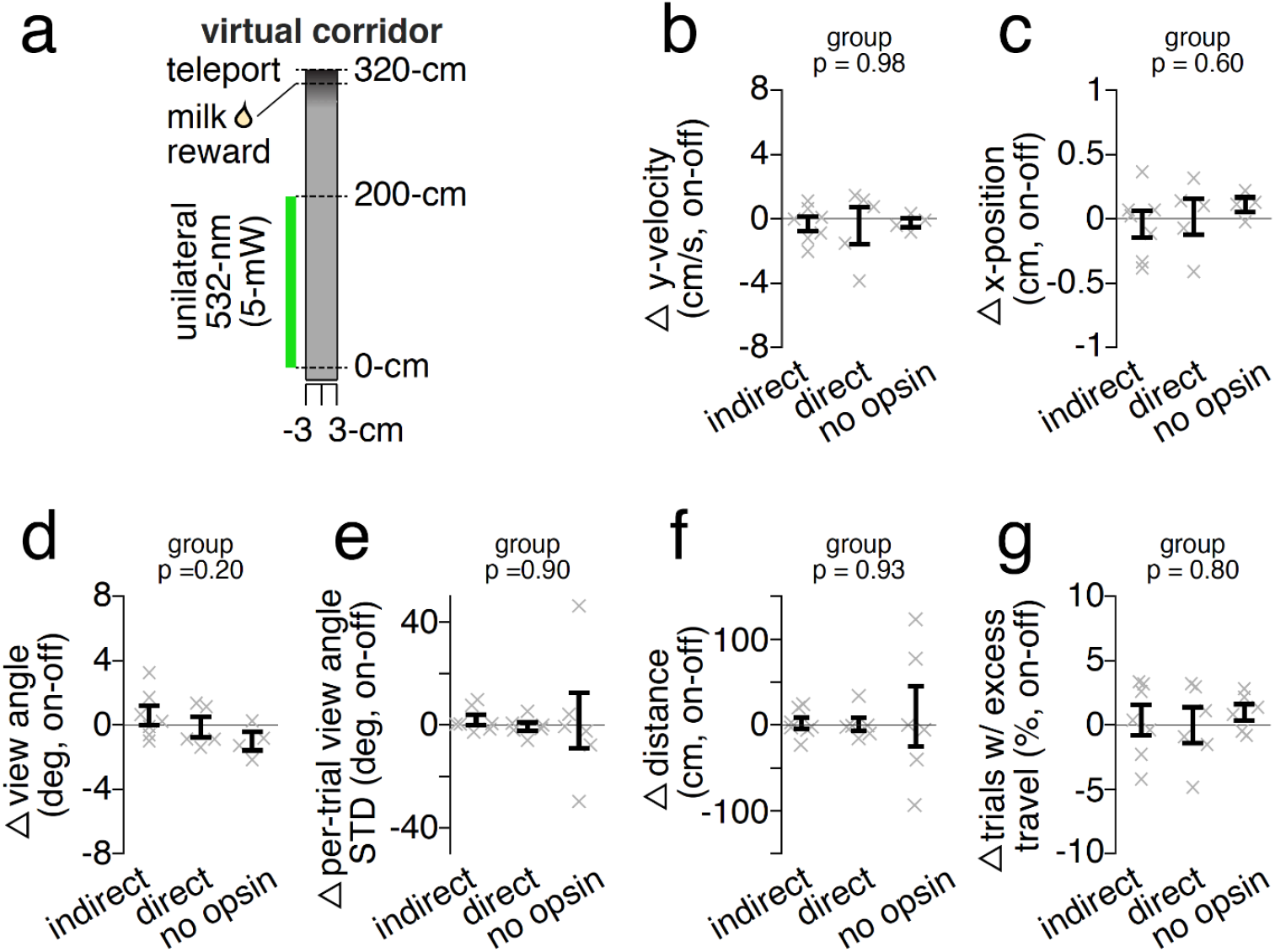
Non-significant motor effects of DMS pathway inhibition compared to non-opsin expressing controls during virtual corridor navigation. (**a**) Schematic of virtual corridor and unilateral delivery of 532-nm light (5-mW) restricted to 0-200cm. (**b**) Difference in average y-velocity (cm/s) during laser on and off trials (on-off) for mice receiving indirect (n = 7 mice, n = 1,712 laser off and n = 1,288 laser on trials) or direct (n = 6 mice, n = 1,088 laser off and n = 757 laser on trials) pathway inhibition of the DMS, or DMS illumination alone (no opsin; n = 5 mice, n = 1,178 laser off and n = 827 laser on trials). p-value denotes significance of one-way ANOVA of group on delta y-velocity (p = 0.98, F_2,13_ = 0.02). (**c**) Same as **b** but for difference in x-position (cm, on-off) contralateral to the laser hemisphere (p = 0.60, F_2,13_ = 0.53). (**d**) Same as **c** but for difference in view angle (deg, on-off) contralateral to the laser hemisphere (p = 0.20, F_2,13_ = 1.90). (**e**) Same as **c** but for difference in mean standard deviation in view angle (deg, on-off). The mean of the standard deviation in view angles sampled in 5-cm steps from 0-300 cm were calculated per trial, and then averaged across all laser off (or on) trials for a mouse (p = 0.94, F_2,16_ = 0.06). Indirect: n = 7 mice, n = 2,109 laser off and n = 1,574 laser on trials; direct: n = 6 mice, n = 1,330 laser off and n = 930 laser on trials; no opsin: n = 6 mice, n = 1,688 laser off and n = 1,199 laser on trials). (**f**) As in **e** but for difference in total distance travelled (cm, on-off) to complete a trial (p = 0.93, F_2,16_ = 0.08). (**g**) As in **e** but for the difference in percentage of trials with excess travel (defined as >10%of corridor length to reward, or >330cm) (p = 0.76,F_2,18_ = 0.28). Throughout solid black lines indicate mean +/- S.E.M. across mice and transparent ‘x’ denote individual mouse mean throughout.

**Extended Data Fig. 3.**
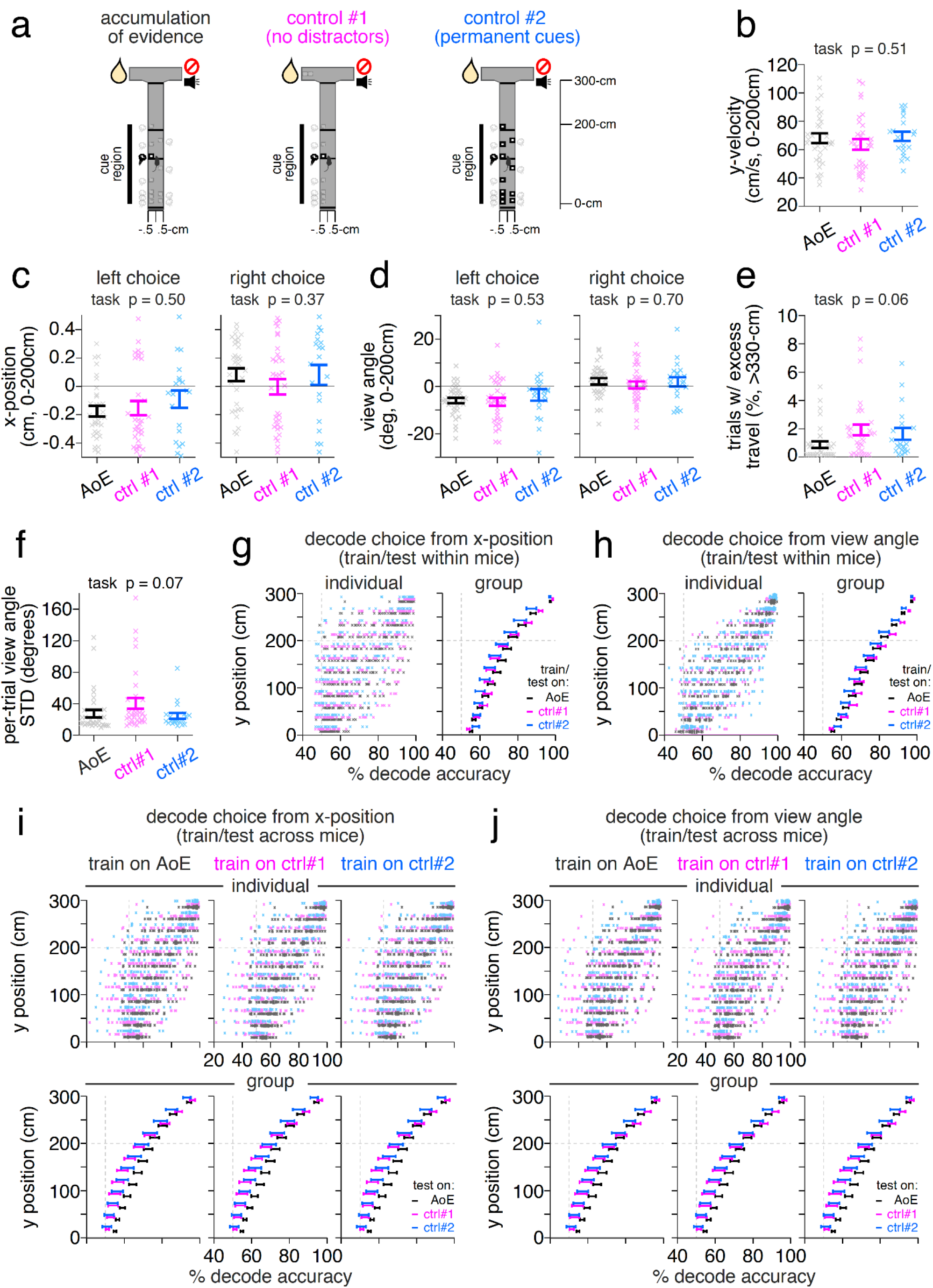
Similar motor performance across three virtual reality T-mazes. Schematic of three virtual reality (VR)-based T-mazes that differ in task requirements. (**b**) Average y-velocity (cm/s) of mice during the cue region (0-200cm) of the accumulation of evidence task (black, n = 32 mice, n = 52,381 trials), no distractors (ctrl #1) task (magenta: n = 32 mice, 56,783 trials), or permanent cues (ctrl #2) task (cyan: n = 20 mice, n = 27,870 trials). Solid bars denote mean +/- S.E.M. across mice while transparent ‘x’ denotes individual mouse mean. p-value denotes one-way ANOVA of task on y-velocity (p = 0.51, F_2,80_ = 0.67). (**c**) Same as **b** but for average x-position (cm) during the cue region (0-200cm) on left and right choice trials. p-value denotes one-way ANOVA of task on x-position (left choice: p = 0.50, F_2,80_ = 0.70; right choice: p = 0.37, F_2,80_ = 1.0). (**d**) Same as **b** but for average view angle (degrees) during the cue region (0-200cm) on left and right choice trials (left choice: p = 0.53, F_2,80_ = 0.64; right choice: p = 0.70, F_2,80_ = 0.37). (**e**) As in **b** but for average percent of trials with excess travel (defined as travel >10% of maze stem, or >330cm). Accumulation of evidence: n = 32 mice, n = 53,833 trials; control #2 (no distractors): n = 32 mice, n = 60,074 trials; control #2 (permanent cues): n = 20 mice, n = 29,192 trials. p-value denotes one-way ANOVA of task on excess travel (p = 0.06, F_2,81_ = 2.9). (**f**) As in **b** but for mean standard deviation in view angle (degrees) per trial (n as in **e**). p-value denotes one-way ANOVA of task on view angle deviation (p = 0.07, F_2,81_ = 2.8). (**g**) Average accuracy of decoding left/right choice based on the trial-by-trial x-position (cm) of mice as a function of y-position in the maze (0-300cm in 25-cm bins). Training and test trial sets were selected within individual mice (80% train, 5-fold cross-validation, re-sampled 10 times). *Left*: Each ‘x’ depicts decoding accuracy at each y-position bin for individual mice performing the evidence accumulation (black), no distractors (ctrl #1, magenta), or permanent cues (ctrl #2, cyan) tasks. *Right*: Group mean and +/-S.E.M. across mice for each task (n as in **b**). (**h**) Same as **f** but for average accuracy of decoding left/right choice based on the trial-by-trial view angle (degrees) of mice (n as in **b**). (**i**) Average accuracy of decoding left/right choice based on the trial-by-trial x-position (cm) of mice as a function of y-position in the maze (0-300cm in 25-cm bins). Training trial sets were randomly selected across all mice (50% total trials, re-sampled 50 times) performing either the accumulation of evidence (*left*, AoE, black), no distractors (*middle*, ctrl#1, magenta), or permanent cues (*right*, ctrl#2, cyan) tasks. Testing trial sets were the 50% of held-out trials in the task used for training, or all trials in the alternate tasks. *Top*: Each ‘x’ depicts average decoding accuracy across all training/tests sets at each y-position bin for individual mice performing the evidence accumulation (black), no distractors (ctrl #1, magenta), or permanent cues (ctrl #2, cyan) tasks. *Right*: Group mean and +/-S.E.M. across mice for each task (n as in **a**). (**j**) Same as **I** but for average accuracy of decoding left/right choice based on the trial-by-trial view angle (degrees) of mice (n as in **b**).

**Extended Data Fig. 4.**
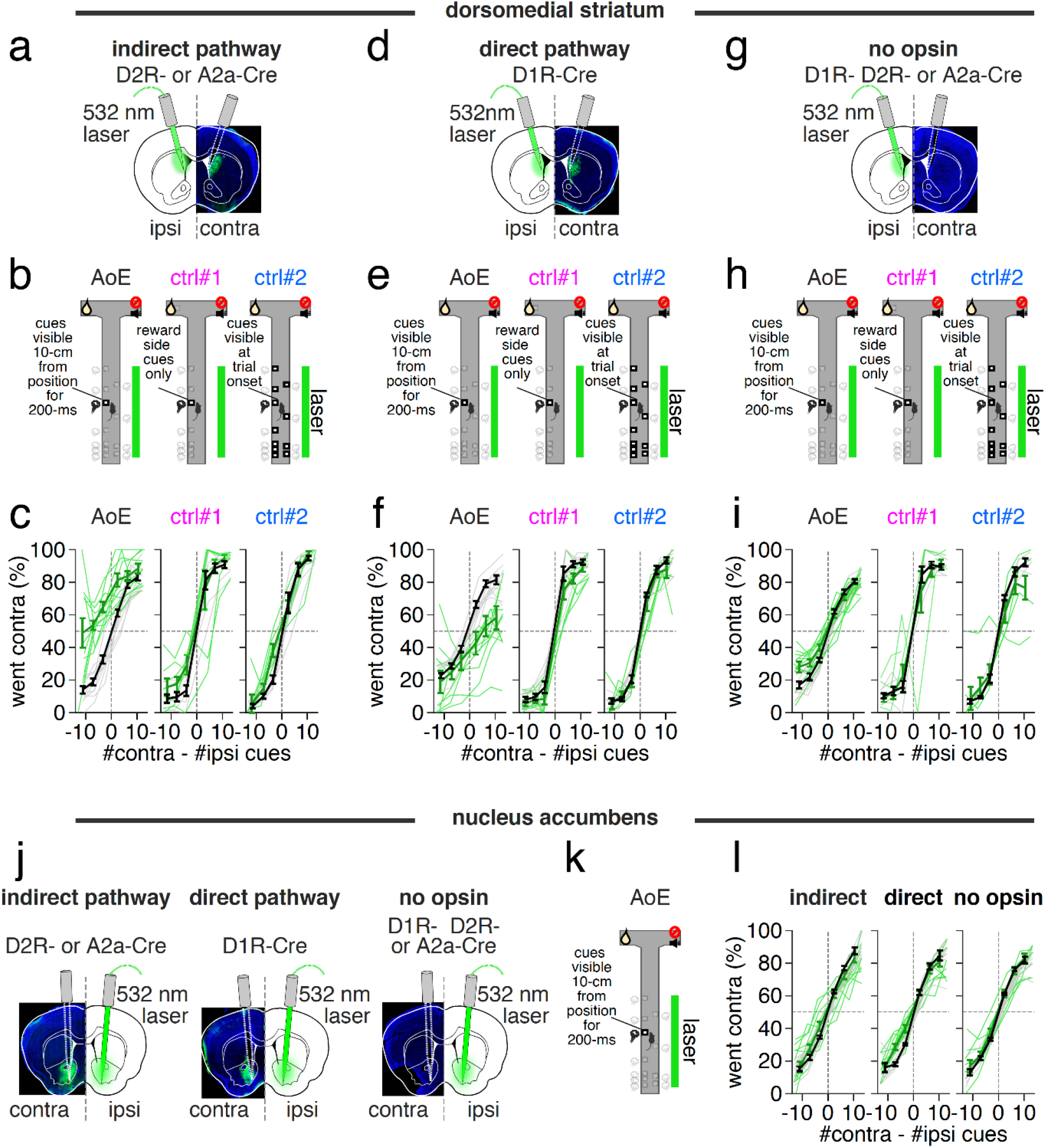
Effects of pathway-specific DMS and NAc inhibition on psychometric performance across virtual reality tasks. (**a**) Schematic of unilateral indirect pathway DMS inhibition with choice defined ipsilateral or contralateral to the hemisphere receiving 532-nm laser illumination. (**b**) Schematic of three virtual reality based decision-making tasks (*left*: accumulation of evidence; *middle*: control #1, no distractors; *right*: control #2, permanent cues) and laser illumination restricted to the cue region (0-200cm). (**c**) Percent of contralateral choice trials as a function of the difference in sensory cues (contralateral-ipsilateral) binned in increments of 5 from −15 to 15. Transparent lines indicate individual mouse mean during laser off (grey) and on (green) trials for mice receiving indirect-pathway DMS inhibition during the evidence accumulation (black, *left*), no distractors (magenta, ctrl#1, *middle*), or permanent cues (cyan, ctrl#2, *right*). Thick lines indicate mean +/- S.E.M. across mice at each evidence bin during laser off (black) and on (green) trials. (**d**) Same as **a** but for mice receiving unilateral direct pathway DMS inhibition. (**e**) same as **b**. (**f**) Same as **c** but for mice receiving direct pathway DMS inhibition. (**g**) Same as **a** but for mice receiving unilateral DMS illumination in the absence of NpHR (no opsin). (**h**) Same as **b**. (**i**) same as **c** but for mice receiving unilateral DMS illumination in the absence of NpHR (no opsin). (**j**) Schematic of unilateral inhibition of NAc indirect (*left*) or direct (*middle*) pathway, or NAc illumination in the absence of NpHR (no opsin). (**k**) Schematic of accumulation of evidence task and delivery of 532-nm light during the cue region (0-200cm). (**l**) As in **c** but for psychometric comparison between groups receiving NAc indirect or direct pathway inhibition, or NAc illumination in the absence of NpHR (no opsin).

**Extended Data Fig. 5.**
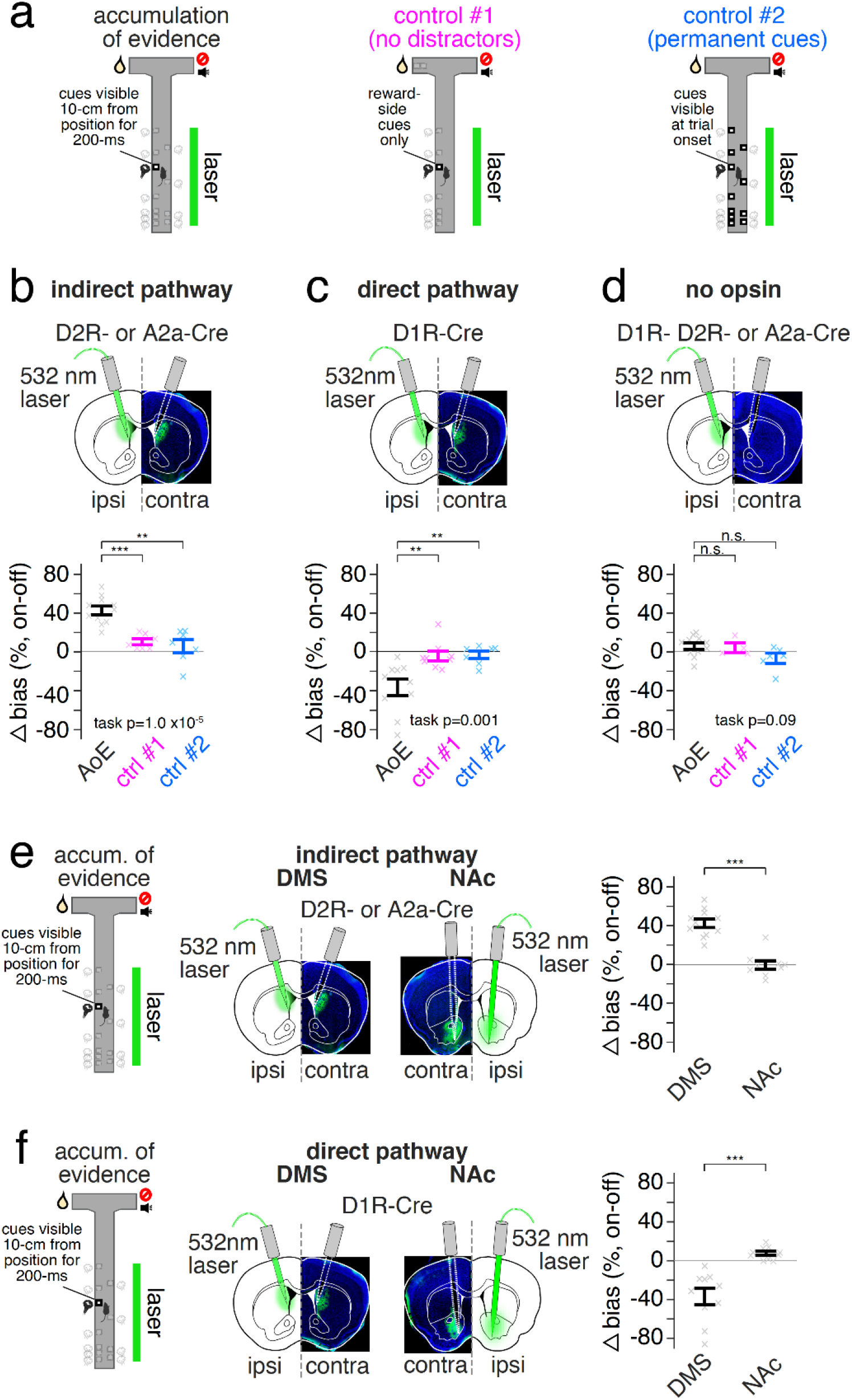
Effects of pathway-specific DMS inhibition on choice are larger in the most demanding task, and stronger than effects of pathway-specific NAc inhibition. (**a**) Schematic of three virtual reality based decision-making tasks (*left*: accumulation of evidence; *middle*: control #1, no distractors; *right*: control #2, permanent cues). (**b**) Schematic of unilateral indirect pathway DMS inhibition with choice defined ipsilateral or contralateral to the hemisphere receiving 532-nm laser illumination (*top*). Difference in choice bias (%, contralateral - ipsilateral) between laser on and off trials (on-off) in mice performing the accumulation of evidence (AoE, black), no distractors (ctrl #1, magenta), or permanent cues (ctrl #2, cyan) tasks. p-value denotes one-way ANOVA of task on delta (on-off) choice bias (p = 1.0 × 10^−5^, F_2,22_ = 20.2). Post-hoc comparisons reflect unpaired, two-tailed Wilcoxon ranksum tests on delta (on-off) choice bias (AoE, n = 11, vs ctrl #1, n = 7: p = 8.0 × 10^−4^, z = 3.4; AoE vs ctrl #2, n = 7: p = 0.001, z = 3.3). (**c**) Same as **b** but for direct pathway DMS inhibition. p-value denotes one-way ANOVA of task on delta (on-off) choice bias (p = 0.001, F_2,23_ = 9.4). Post-hoc comparisons reflect two-tailed, unpaired Wilcoxon ranksum tests (AoE, n = 10, vs ctrl #1, n = 9: p = 0.002, z = −3.0; AoE vs ctrl #2, n = 7: p = 0.005, z = −2.8). (**d**) Same as **b** but for DMS illumination in the absence of NpHR (no opsin). p-value denotes one-way ANOVA of task on delta (on-off) choice bias (p = 0.09, F_2,16_ = 2.8). Post-hoc comparisons reflect two-tailed, unpaired Wilcoxon ranksum tests (AoE, n= 11, vs ctrl #1, n = 4: p = 0.65, z = 0.46; AoE vs ctrl #2, n = 6: p = 0.06, z = 1.8). (**e**) Schema of evidence accumulation task (*left*), unilateral inhibition of indirect pathway in the DMS (*middle left*) or NAc (*middle right*), and delta (on-off) choice bias in mice receiving indirect pathway DMS (n = 11) or NAc (n = 9) inhibition (*right*). Statistical comparison reflects two-tailed, unpaired Wilcoxon ranksum test (DMS vs NAc: p = 2.6 × 10^−4^, z = 3.6). (**f**) Same as **e** but for direct pathway DMS (n = 10) or NAc (n = 10) inhibition. Statistical comparison reflects two-tailed, unpaired Wilcoxon ranksum test (DMS vs NAc: p = 1.8 × 10^−4^, z = −3.7). Throughout solid bars denote mean +/- S.E.M. across mice and transparent ‘x’ denote individual mouse means.

**Extended Data Fig. 6.**
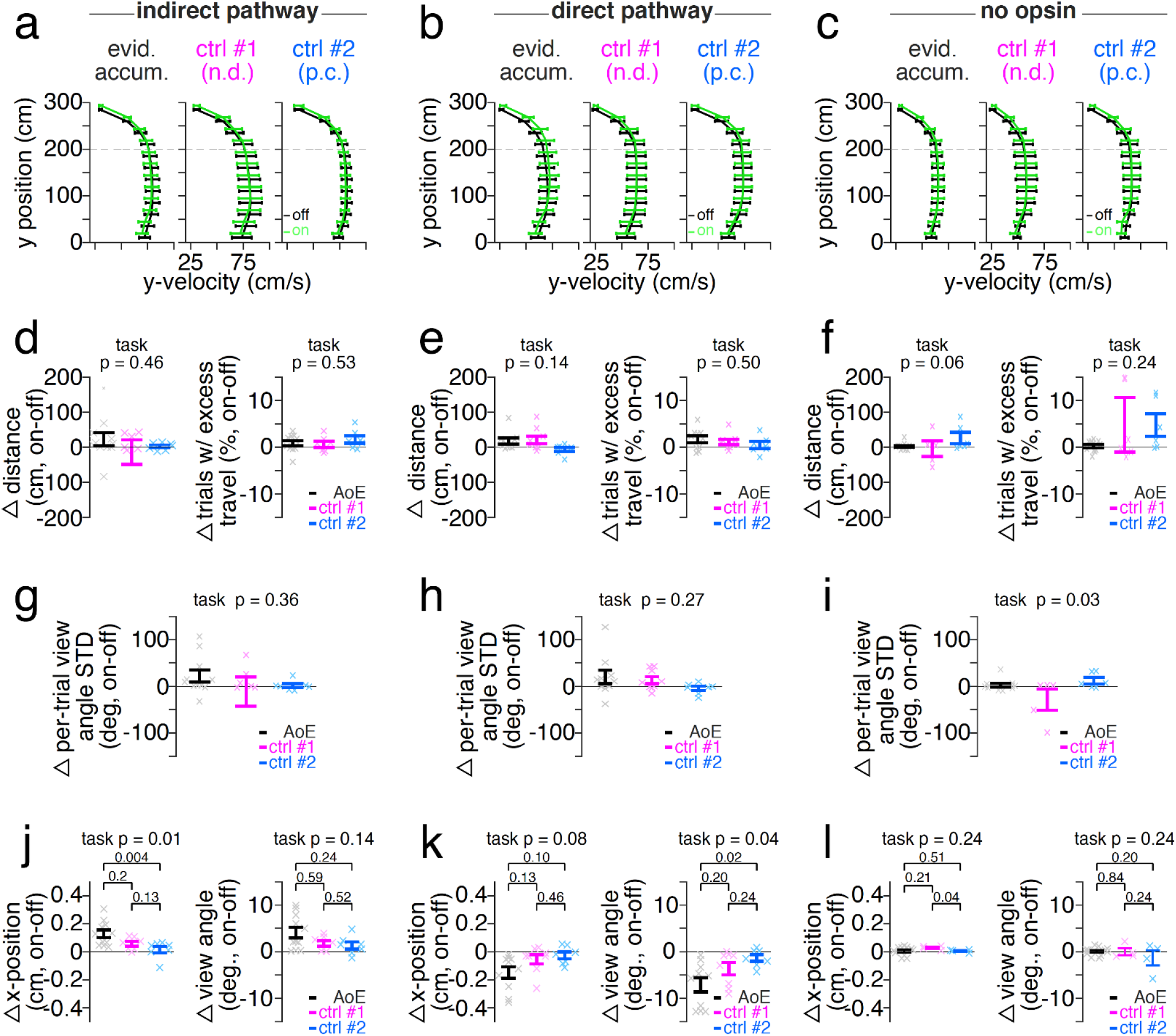
Inhibition of DMS pathways has limited impact on motor performance across VR-based decision-making tasks. (**a**) Mean +/- S.E.M. y-velocity (cm/s) as a function of y-position (0-300cm in 25cm bins) during laser off (black) or laser on (green) trials across mice receiving DMS indirect pathway inhibition during the evidence accumulation (*left*: n = 11 mice, n = 16,935 laser off and n = 3,390 laser on trials), no distractors (*middle*, ctrl #1: n = 7 mice, n = 13,706 laser off and n = 3,288 laser on trials) or permanent cues (*right*, ctrl #2: n = 6 mice, n = 4,033 laser off and n = 929 laser on trials). (**b**) Same as **a** but for mice receiving direct pathway inhibition during the evidence accumulation (*left*: n = 10 mice, n = 14,030 laser off and n = 3,103 laser on trials), no distractors (*middle*, ctrl #2: n = 8 mice, n = 14,647 laser off and n = 3,682 laser on trials) or permanent cues (*right*, ctrl #3: n = 7 mice, n = 6,061 laser off and n = 1,494 laser on trials) tasks. (**c**) Same as **a** but for mice receiving DMS illumination in the absence of NpHR (no opsin) during the evidence accumulation (*left*: n = 11 mice, n = 21,422 laser off and n = 5,113 laser on trials), no distractors (*middle*, ctrl #1: n = 4 mice, n = 3,654 laser off and n = 901 laser on trials), or permanent cues (*right*, ctrl #2: n = 4 mice, n = 3,975 laser off and n = 923 laser on trials) tasks. (**d**) Mean +/- S.E.M. in delta (on-off) distance (cm) traveled (left) and delta (on-off) trials (%) with excess travel greater than 10% of maze stem (or >330cm) (right) in mice receiving indirect pathway inhibition during the evidence accumulation (black, n = 11 mice, n = 22,090 laser off and n = 4,378 laser on trials), no distractors (magenta, n = 7 mice, n = 14,826 laser off and n = 3,591 laser on trials), or permanent cues (n = 6 mice, n = 4,447 laser off and n = 1050 laser on trials) tasks. p-value denotes one-way ANOVA of task on delta (on-off) distance (p = 0.45, F_2,22_ = 0.81) or excess travel (p = 0.52, F_2,22_ = 0.66). (**e**) Same as **d** but for delta (on-off) distance (cm) traveled (left) or delta percent trials with excess travel (right) in mice receiving direct pathway inhibition during the evidence accumulation (black, n = 10 mice, n = 20,914 laser off and n = 4,721 laser on trials), no distractors (magenta, n = 9 mice, n = 15,778 laser off and n = 3,992 laser on trials), or permanent cues (n = 7 mice, n = 6,430 laser off and n = 1,591 laser on trials) tasks. p-value denotes one-way ANOVA of task on delta (on-off) distance (p = 0.13, F_2,23_ = 2.2) or excess travel (p = 0.50, F_2,23_ = 0.71). (**f**) Same as **d** but for delta (on-off) in distance (cm) traveled (left) or percent trials with excess travel (right) in mice receiving DMS illumination in the absence of NpHR (no opsin) during the evidence accumulation (black, n = 11 mice, n = 28,557 laser off and n = 6,772 laser on trials), no distractors (magenta, n = 5 mice, n = 4,118 laser off and n = 1,002 laser on trials), or permanent cues (n = 6 mice, n = 4,360 laser off and n = 1,038 laser on trials) tasks. p-value denotes one-way ANOVA of task on delta (on-off) distance (p = 0.06, F_2,19_ = 3.3) or excess travel (p = 0.23, F_2,19_ = 1.6). (**g**) Same as **d** but for delta (on-off) in per-trial standard deviation in view angle in mice receiving DMS indirect pathway inhibition across tasks (p = 0.34, F_2,22_ = 1.1, n as in **d**). (**h**) Same as **g** but for mice receiving DMS direct pathway inhibition across tasks (p = 0.27, F_2,23_ = 1.4, n as in **e**). (**i**) Same as **g** but for mice receiving DMS illumination (no opsin) in the absence of NpHR (p = 0.03, F_2,19_ = 4.3, n as in **f**). (**j**) Delta (on-off) x-position (cm) (*left*) or view angle (degrees) (*right*) during the cue region (0-200 cm) in mice receiving DMS indirect pathway inhibition during the accumulation of evidence (black), no distractors (control #1, magenta), or permanent cues (control #2, cyan) tasks (n as in **a**). One-way ANOVA of task on delta (on-off) x-position (p = 0.01, F_2,22_ = 5.6). Post-hoc, two-tailed, unpaired Wilcoxon ranksum test on delta (on-off) x-position (AoE v control #1: p = 0.2, z = 1.3; AoE v control #2: p = 0.004, z = 2.9; control #1 v control #2: p = 0.13, z = 1.5). One-way ANOVA of task on delta (on-off) view angle (p = 0.14, F_2,22_ = 2.2). Post-hoc, two-tailed, unpaired Wilcoxon ranksum test on delta (on-off) view angle (AoE v control #1: p = 0.58, z = 0.5; AoE v control #2: p = 0.24, z = 1.78; control #1 v control #2: p = 0.52, z = 0.6). (**k**) Same as **j** but for mice receiving DMS direct pathway inhibition (n as in **b**). One-way ANOVA of task on delta (on-off) x-position (p = 0.08, F_2,23_ = 2.8). Post-hoc, two-tailed unpaired Wilcoxon ranksum test on delta (on-off) x-position (AoE v control #1: p = 0.13, z = −1.5; AoE v control #2: p = 0.1, z = −1.6; control #1 v control #2: p = 0.46, z = −0.7). One-way ANOVA of task on delta (on-off) view angle (p = 0.02, F_2,23_ = 3.6). Post-hoc, two-tailed, unpaired Wilcoxon ranksum test on delta (on-off) view angle (AoE v control #1: p = 0.21, z = −1.3; AoE v control #2: p = 0.03, z = −2.1; control #1 v control #2: p = 0.24, z = −1.6). (**l**) Same as **j** but for mice receiving DMS illumination in the absence of NpHR (no opsin, n as in **c**). One-way ANOVA of task on delta (on-off) x-position (p = 0.24, F_2,18_ = 1.54). Post-hoc, two-tailed, unpaired Wilcoxon ranksum test on delta (on-off) x-position (AoE v control #1: p = 0.21, z = −1.24; AoE v control #2: p = 0.51, z = 0.06; control #1 v control #2: p = 0.04, z = 2.0). One-way ANOVA of task on delta (on-off) view angle (p = 0.23, F_2,18_ = 1.56). Post-hoc, two-tailed, unpaired Wilcoxon ranksum test on delta (on-off) view angle (AoE v control #1: p = 0.84, z = 0.19; AoE v control #2: p = 0.20, z = 1.2; control #1 v control #2: p = 0.24, z = 1.7). Throughout solid bars denote mean +/- S.E.M. and transparent ‘x’ indicates individual mouse mean.

**Extended Data Fig. 7.**
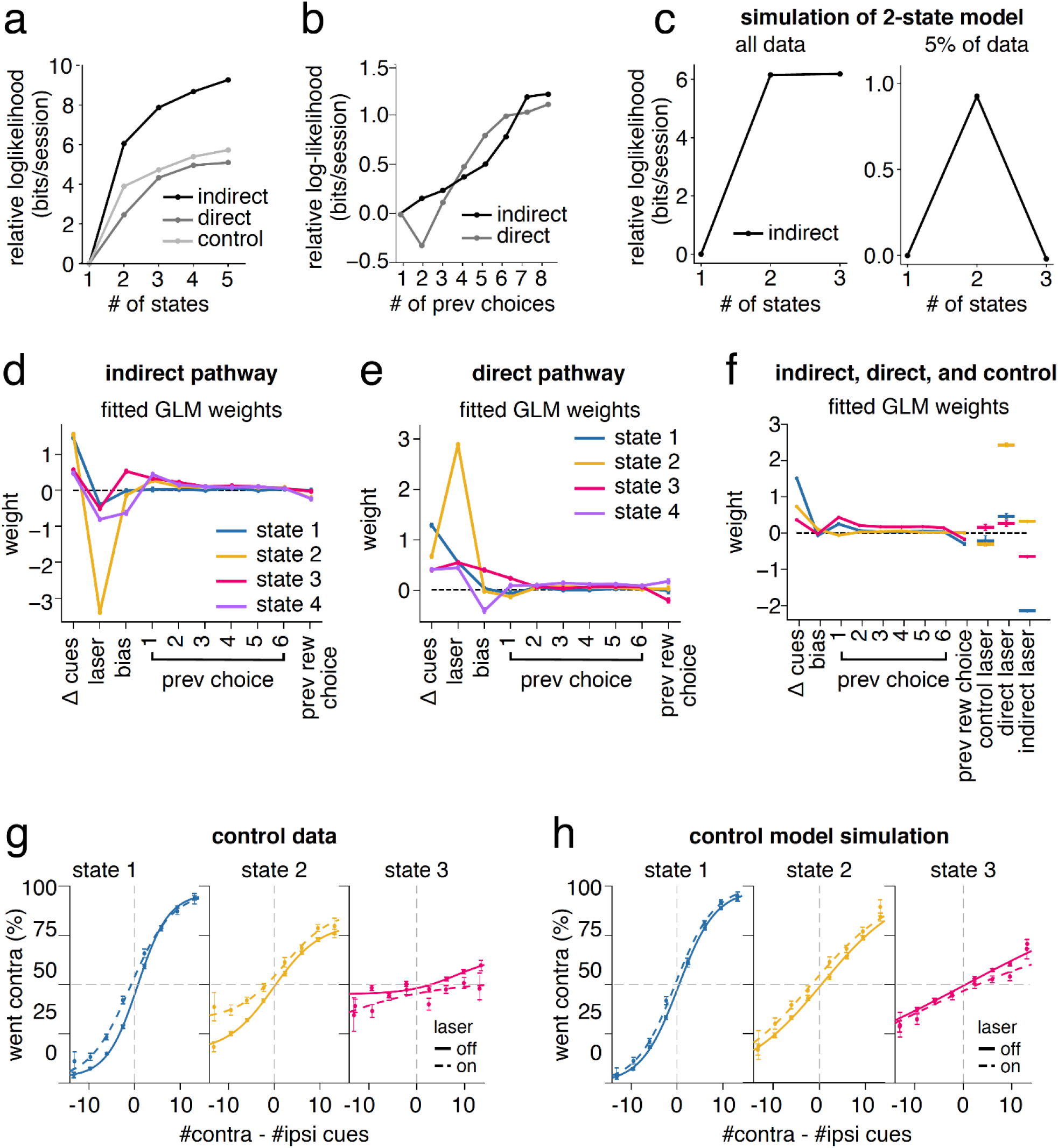
Model selection and control data analyses for the GLM-HMM. (**a**) (**a**) Comparison of the log-likelihood of the data using GLM-HMMs with different numbers of states for mice inhibited in the DMS direct pathway (dark gray), or indirect pathway (light gray), and mice without DMS opsin (black). All values are relative to the log-likelihood of the standard GLM (1-state GLM-HMM). Values are calculated in bits per session (see **Methods**). Solid curves denote mean +/- S.E.M. of five different test sets. Held-out data for test sets was selected as a random 20% of sessions, using the same number of sessions for each mouse. (**b**) Same as **a** but with different numbers of previous choice covariates using a 3-state GLM-HMM. (**c**) Comparison of the log-likelihood of simulated data using GLM-HMMs with different numbers of states. Data was simulated from a 2-state GLM-HMM that had been fit to data for mice inhibited in the indirect pathway of the DMS and then cross-validation performed either on the entire simulated dataset (~54000 trials, left) or a subset of 5% of the data (2600 trials, right). All values are relative to the log-likelihood of the 1-state GLM. Values are calculated in bits per session (see Methods). Solid curves denote the average of five different test sets. Held-out data for test sets was selected as a random 20% of sessions. Performing cross validation on a small subset of the data serves to demonstrate that the log-likelihood does in fact decrease as the model starts to overfit. This is difficult to see with large datasets where overfitting is less of a concern and therefore the log-likelihood begins to flatten rather than decrease. (**d**) Fitted GLM weights for the 4-state model using aggregated data from all mice inhibited in the indirect pathway of the DMS. Error bars denote (+/-1) posterior standard deviation for each weight. The magnitude of the weight represents the relative importance of that covariate in predicting choice, whereas the sign of the weight indicates the side bias. (**e**) Same as **d** but for mice inhibited in the DMS direct pathway. (**f**) GLM weights fitted to a concatenated data set consisting of the indirect, direct, and control (no opsin) groups. Solid lines on the left connect covariates that are shared across groups. Horizontal marks on the right denote laser weights, which were learned separately for each group. Error bars denote the posterior standard deviation of each weight. (**g**) Percent of contralateral choice based on the difference in contralateral versus ipsilateral cues in each trial for mice in the control (no opsin) group. To compute psychometric functions, trials were assigned to each state by taking the maximum of the model’s posterior state probabilities on each trial. Error bars denote +/-1 S.E.M. for light off (solid) and light on (dotted) trials. Solid curves denote logistic fits to the concatenated data across mice for light off (solid) and light on (dotted) trials. (**h**) Same as **f** but for data simulated from the model fit to mice in the control group (see **Methods**).

**Extended Data Fig. 8.**
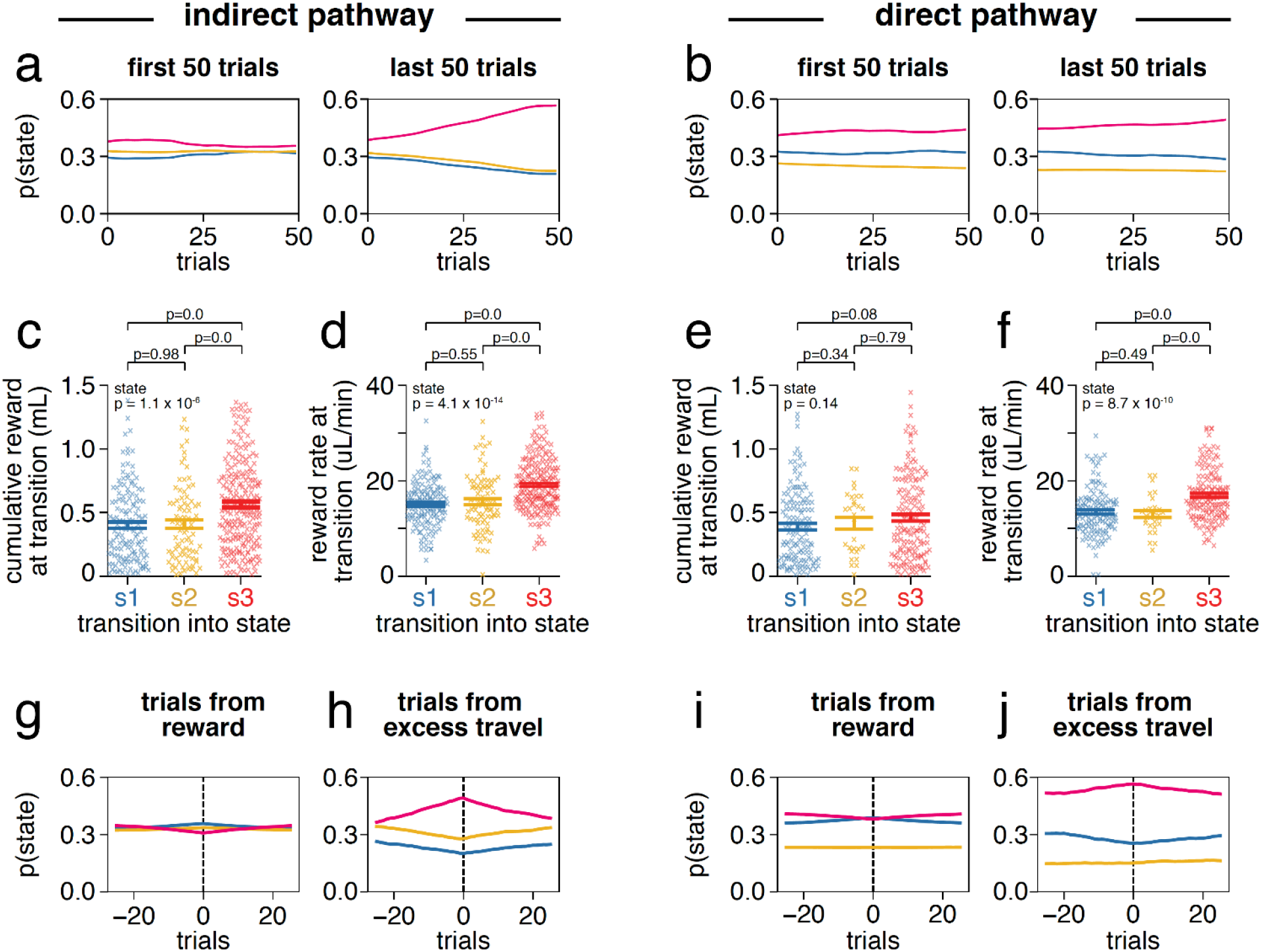
GLM-HMM state 3 is associated with indicators of task disengagement. (**a**) The mean posterior probability of each state over the first and last 50 trials of a session, averaged across all sessions for mice inhibited in the indirect pathway of the DMS (n=271 sessions). (**b**) Same as **a** but for mice receiving DMS direct pathway inhibition (n=266 sessions). (**c**) Mean +/- S.E.M. of the cumulative reward received in a session prior to transitions into state 1 (n = 142), state 2 (n = 85), or state 3 (n = 237) in indirect pathway mice. One-way ANOVA of transition state on cumulative reward (p = 1.0 × 10^−6^; F_2,460_ = 14.2). Unpaired, two-tailed Wilcoxon ranksum comparison between transition types (state 1 vs 2: p = 0.96, z = −0.03; state 2 vs 3: p = 0, z = −3.6; state 1 vs 3: p = 0, z = −4.5). (**d**) Mean +/- S.E.M. of the reward rate (uL/min) in a session prior to transitions into each state for indirect pathway mice. Reward rate was calculated as the sum of reward received from the start of the session up to the transition trial divided by the sum of the duration of all trials from the start of the session up to the transition trial. One-way ANOVA of transition state on reward rate (p = 4.1 × 10^−14^; F_2,460_ = 32.9). Unpaired, two-tailed Wilcoxon ranksum comparison between transition types (state 1 vs 2: p = 0.55, z = −0.6; state 2 vs 3: p = 0, z = −4.9; state 1 vs 3: p = 0, z = −7.4). (**e**) Same as **c** but for direct pathway mice (state 1: n = 140; state 2: n = 29; state 3: n = 159). One-way ANOVA of transition state on cumulative reward (p = 0.14; F_2,325_ = 1.99). Unpaired, two-tailed Wilcoxon ranksum comparison between transition types (state 1 vs 2: p = 0.35, z = −0.9; state 2 vs 3: p = 0.78, z = −0.27; state 1 vs 3: p = 0.08, z = −1.74). (**f**) Same as **d** but for direct pathway mice. One-way ANOVA of transition state on reward rate (p = 8.7 × 10^−10^; F_2,325_ = 22.6). Unpaired, two-tailed Wilcoxon ranksum comparison between transition types (state 1 vs 2: p = 0.49, z = 0.69; state 2 vs 3: p = 0.0, z = −4.2; state 1 vs 3: p = 0.0, z = −5.9). (**g**) The mean posterior probability of each state aligned +/- 25 trials to trials in which reward was received for indirect pathway mice. (**h**) Same as **g** but state probability aligned to trials with excess travel (defined as 10% greater than the maze stem, or 330cm). (**i**) Same as **g** but for direct pathway mice. (**j**) Same as **h** but for direct pathway mice.

**Extended Data Fig. 9.**
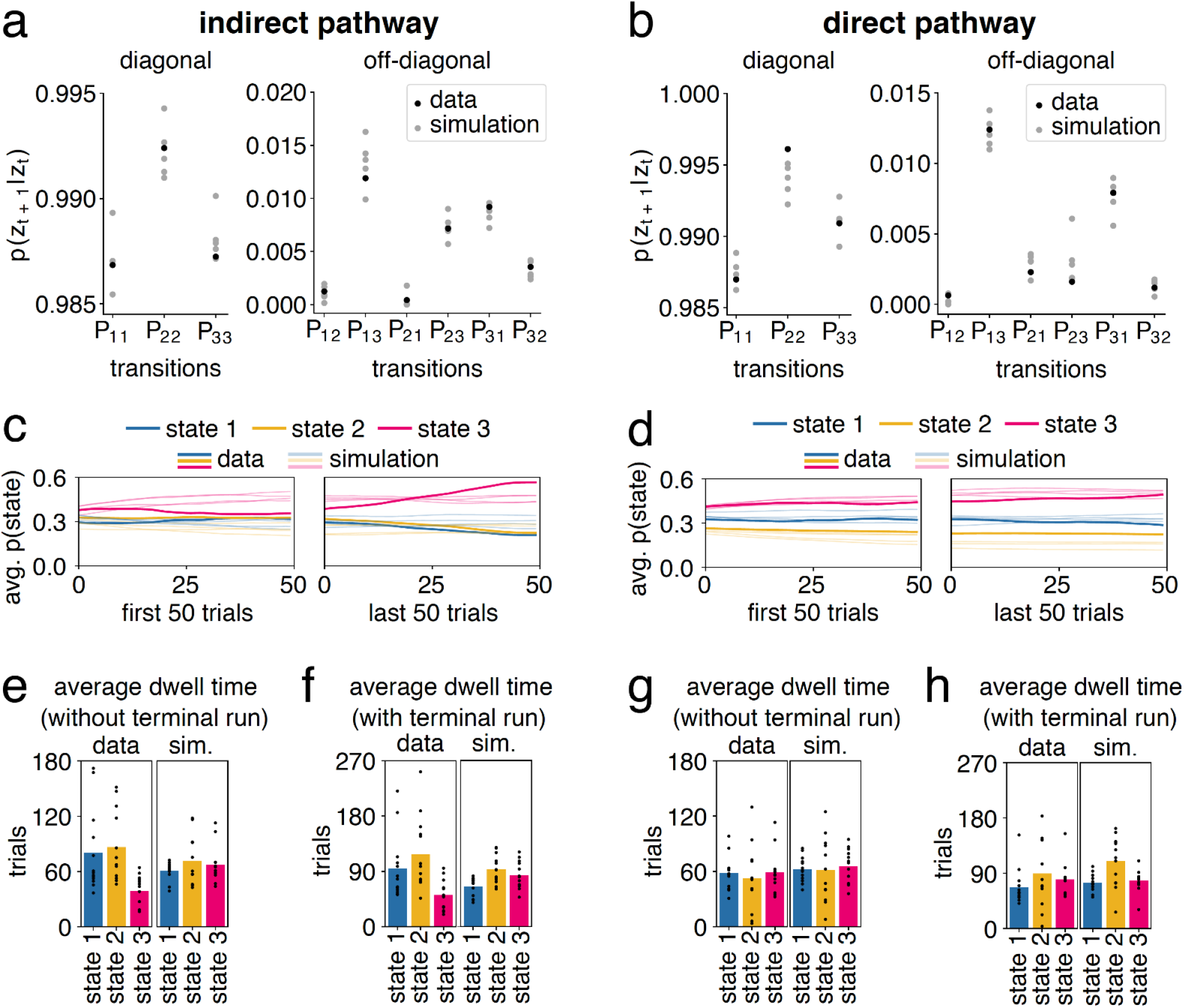
Model simulations recapitulate transition and state characteristics of real data. (**a**) Transition probabilities of the model fit to data from mice inhibited in the DMS indirect pathway (black) and from five simulated datasets generated from the model fit to mice inhibited in the indirect pathway of the DMS (gray), shown separately for diagonal (left) and off-diagonal (right) probabilities. (**b**) Same as **a** but for mice inhibited in the direct pathway of the DMS. (**c**) The posterior probability of each state over the first and last 50 trials of a session, averaged across all sessions for mice inhibited in the indirect pathway of the DMS (n=271). Dark lines denote average for real data (same as Fig. 7E) and faded lines indicate averages for each of the five simulations. (**d**) Same as **c** but for mice inhibited in the direct pathway of the DMS (dark lines are the same as shown in Fig. 7F). (**e**) Dwell times showing the average consecutive number of trials that mice inhibited in the DMS indirect pathway spent in each state for real data (left; range 39-86 trials, average session length 202 trials, same as shown in **Fig. 7g**) and one simulated dataset (right; range 60-71 trials, average session length 202 trials). Black dots show averages for individual mice (n=13). We removed the last run in each session (including any run that lasted the entire session length) from the analysis, as the termination of the session prematurely truncated the length of those runs. (**f**) Same as **e** but without removing the last run in each session for real data (left; range 51-118 trials, average session length 202 trials) and one simulated dataset (right; range 65-93 trials, average session length 202 trials). (**g**) Same as **e** but for mice inhibited in the direct pathway of the DMS for real data (left; range 52-59 trials, average session length 185 trials, same as shown in **Fig. 7g**) and one simulated dataset (right; range 61-66 trials, average session length 185 trials). Black dots show averages for individual mice (n=13). (**h**) Same as **g** but without removing the last run in each session for real data (left; 67-89 trials, average session length 185 trials) and one simulated dataset (right; range 74-110 trials, average session length 185 trials).

**Extended Data Fig. 10.**
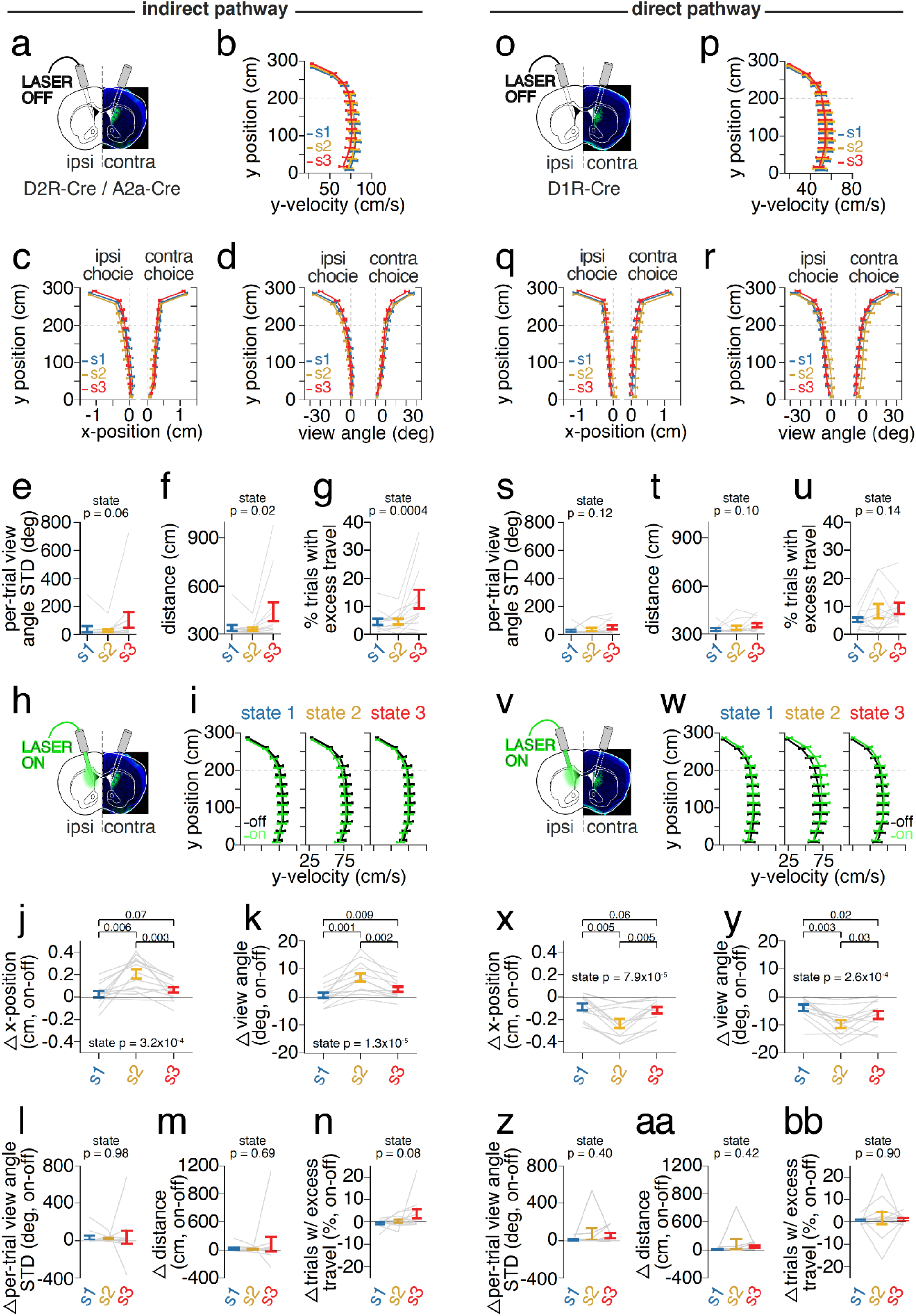
Comparison of motor performance across GLM-HMM states with and without pathway-specific DMS inhibition. (**a**) Schematic denoting analysis of motor performance across GLM-HMM states on laser off trials only (panels **b-g**) in mice unilaterally coupled to a fiberoptic for indirect pathway inhibition. (**b**) Average y-velocity (cm/s) during laser off trials as a function of y-position in the maze (0-300 cm in 25-cm bins) in indirect pathway mice across GLM-HMM states (state 1, blue: n = 13,394 trials; state 2, yellow: n = 13,570 trials; state 3, red: n = 16,982 trials). (**c**) As in **b** but for average x-position (cm) on ipsilateral or contralateral choice trials (n as in **b**). (**d**) As in **c** but for average view angle (degrees) on ipsilateral and contralateral choice trials (n as in **b**). (**e**) Mean per-trial standard deviation in view angle during laser off trials across GLM-HMM states (state 1, blue: n = 13,854 trials; state 2, yellow: n = 14,201 trials; state 3, red: n = 18,258 trials). p-value denotes one-way repeated measures ANOVA of state on view angle deviation (p = 0.06, F_2,24_ = 3.2). (**f**) As in **e** but for average distance traveled (cm) per trial. p-value denotes one-way repeated measures ANOVA of state on distance (p = 0.02, F_2,24_ = 5.0, n as in **e**). (**g**) As in **e** but for average percent of trials with excess travel. p-value denotes one-way repeated measures ANOVA of state on excess travel (p = 0.0004, F_2,24_ = 10.9, n as in **e**). (**h**) Schematic denoting analysis of effects of indirect pathway DMS inhibition on motor performance across GLM-HMM states in **i-n**. (**i**) As in **b** but for average y-velocity on laser off (black) or laser on (green) trials across GLM-HMM states (n of laser off trials as in **b-g**, n of laser on trials: state 1, blue: n = 2,302 trials; state 2, yellow: n = 1,858 trials; state 3, red: n = 3,005 trials). (**j**) As in **c** but for delta (on-off) x-position (cm) during the cue region (0-200cm) across GLM-HMM states in mice with indirect pathway inhibition. p-value denotes one-way repeated measures ANOVA of state on delta x-position (p = 3.2×10^−4^, F_2,24_ = 11.4, n as in **i**). Post-hoc comparisons reflect two-tailed, paired Willcoxon signed rank tests between states (state 1 vs state 3: p = 0.07, z = 1.7; state 1 vs state 2: p = 0.006, z = 2.7; state 2 vs state 3: p = 0.03, z = 2.4). (**k**) As in **j** but for delta (on-off) view angle (degrees). p-value denotes one-way repeated measures ANOVA of state on delta view angle (p = 1.2×10^−5^, F_2,26_ = 18.7, n as in **i**). Post-hoc comparisons reflect two-tailed, paired Willcoxon signed rank tests between states (state 1 vs state 3: p = 0.009, z = 2.6; state 1 vs state 2: p = 0.001, z = −3.18; state 2 vs state 3: p = 0.002, z = −3.1). (**l**) Same as **e** but for delta (on-off) mean per-trial view angle standard deviation across GLM-HMM states in mice with indirect pathway inhibition (n of laser off trials as in **e-g**, n of laser on trials: state 1, blue: n = 2,887 trials; state 2, yellow: n = 2,713 trials; state 3, red: n = 2,970 trials). p-value denotes one-way repeated measures ANOVA of state on delta view angle deviation (p = 0.97, F_2,24_ = 0.03, n as in **l**). (**m**) Same as **f** but for delta (on-off) in mean per-trial distance (cm) traveled across GLM-HMM states with indirect pathway inhibition (p = 0.68, F_2,24_ = 0.38, n as in **l**). (**n**) Same as **g** but for delta (on-off) in percent of trials with excess travel across GLM-HMM states with direct pathway inhibition (p = 0.08, F_2,24_ = 2.8, n as in **l**). (**o**) As in **a** but schematic denoting analysis of motor performance across GLM-HMM states on laser off trials only in mice unilaterally coupled to a fiberoptic for direct pathway inhibition in **p-u**. (**p**) As in **b** but for y-velocity (cm/s) on laser off trials across GLM-HMM states in direct pathway mice (state 1, blue: n = 12,294 laser off and n = 2,302 laser on trials; state 2, yellow: n = 9,201 laser off and n = 1,858 laser on trials; state 3, red: n = 16,239 laser off and n = 3,005 laser on trials). (**q**) As in **c** but x-position (cm) for direct pathway mice (n as in **p**). (**r**) As in **d** but for view angle (degrees) for direct pathway mice (n as in **p**). (**s**) As in **e** but for mean per-trial view angle standard deviation across GLM-HMM states in direct pathway mice (state 1, blue: n = 13,403 laser off and n = 2,508 laser on trials; state 2, yellow: n = 9,555 laser off and n = 1,969 laser on trials; state 3, red: n = 18,292 laser off and n = 3,450 laser on trials). p-value denotes one-way repeated measures ANOVA of state on per-trial view angle standard deviation (p = 0.12, F_2,24_ = 2.3). (**t**) As in **f** but for distance (cm) in direct pathway mice (p = 0.1, F_2,24_ = 2.5). (**u**) As in **g** but for percent trials with excess travel in direct pathway mice (p = 0.14, F_2,24_ = 2.1). (**v**) As in **h** but schematic denoting analysis of effects of direct pathway DMS inhibition on motor performance across GLM-HMM states in **w-bb**. (**w**) As in **i** but for the mean y-velocity (cm/s) on laser on (green) and off (black) trials across GLM-HMM states in direct pathway mice. (**x**) As in **j** but for the delta (on-off) x-position (cm) across GLM-HMM states in direct pathway mice. p-value denotes one-way repeated measures ANOVA of state on delta x-position (p = 7.9×10^−5^, F_2,24_ = 14.9). Posthoc comparisons reflect two-tailed, paired Willcoxon signed rank tests between states (state 1 vs state 3: p = 0.06, z = 1.8; state 1 vs state 2: p = 0.005, z = 2.8; state 2 vs state 3: p = 0.005, z = 2.8). (**y**) As in **k** but for delta (on-off) view angle (degrees) across GLM-HMM states in direct pathway mice. p-value denotes one-way repeated measures ANOVA of state on delta view angle (p = 2.6×10^−4^, F_2,24_ = 12.3). Posthoc comparisons reflect two-tailed, paired Willcoxon signed rank tests between states (state 1 vs state 3: p = 0.03, z = 2.3; state 1 vs state 2: p = 0.003, z = 2.98; state 2 vs state 3: p = 0.03, z = 2.1). (**z**) As in **l** but for delta (on-off) mean per-trial view angle standard deviation (degrees) in direct pathway mice (p = 0.40, F_2,24_ = 0.94, n as in **s-u**). (**aa**) as in **m** but for delta (on-off) in mean distance (cm) traveled in direct pathway mice (p = 0.43, F_2,24_ = 0.89). (**bb**) as in **n** but for delta (on-off) in percent trials with excess travel in direct pathway mice (p = 0.90, F_2,24_ = 0.1). Throughout solid colored bars denote mean +/- S.E.M. while transparent grey lines reflect individual mouse mean.

## SUPPLEMENTARY FIGURES

**Supplementary Fig. 1.**
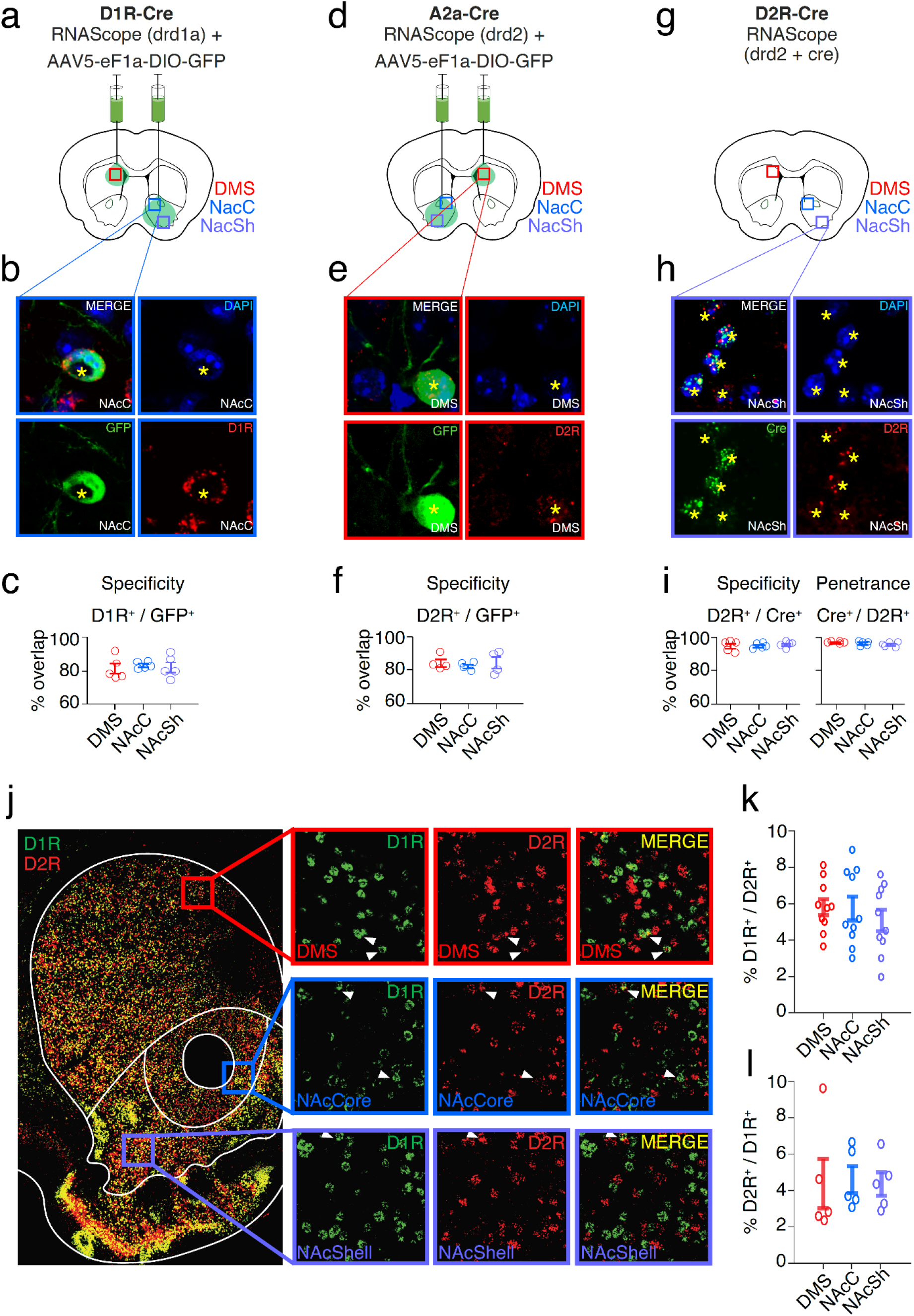
Transgenic mouse lines faithfully report indirect and direct pathways across striatal subregions. (**a**) Schematic of viral delivery of AAV5-eF1a-DIO-GFP to the dorsomedial striatum (DMS) or nucleus accumbens (NAc) on opposite hemispheres of D1R-Cre mice. Red, blue, and purple squares denote representative areas for stereological quantification of viral co-expression with a drd1 mRNA probe (RNAScope) in the DMS, NAc core (NAcC), or NAc shell (NAcSh), respectively. (**b**) Example fluorescent confocal image (63x objective, 5x digital zoom) of the NAc core from a D1R-Cre mouse displaying virally-expressed GFP (green), drd1 mRNA (red), and DAPI (blue). (**c**) Percentage of GFP^+^ neurons co-expressing drd1 mRNA from 2 D1R-Cre mice across the DMS (red; n = 5 sections; 193 GFP^+^ neurons), NAcC (blue; n = 5 sections; 298 GFP^+^ neurons), or NacSh (purple; n = 4 sections; 312 GFP^+^ neurons). (**d**) Same as **a**, but for quantification of viral co-expression with a drd2 mRNA probe in A2a-Cre mice. (**e**) Same as **b**, but for an example image of the DMS from an A2a-Cre mouse and displaying drd2 mRNA (red). (**f**) Same as **c**, but for percentage of virally-expressed GFP^+^ neurons co-expressing drd2 mRNA in 2 A2a-Cre mice across the DMS (red; n = 4 sections; 312 GFP^+^ neurons), NAcC (blue; n = 4 sections; 326 GFP^+^ neurons), or NacSh (purple; n = 4 sections; 312 GFP^+^ neurons). (**G**) Same as **A** and **D**, but for quantification of co-expression of drd2 and cre mRNA in 2 D2R-Cre mice. (**h**) Same as **b** and **e**, but for an example image of the NAcSh from a D2R-Cre mouse and displaying cre mRNA (green) and drd2 mRNA (red). (**i**) *Left*: same as **c** and **f**, but for percentage of neurons with cre mRNA co-expressing drd2 mRNA in 2 D2R-Cre mice across the DMS (red; n = 5 sections; 1302 cre^+^ neurons), NAcC (blue; n = 5 sections; 1,104 cre^+^ neurons), or NacSh (purple; n = 4 sections; 1,187 cre^+^ neurons). *Right*: same as *left* but for neurons with drd2 mRNA co-expressing cre mRNA across DMS (red; n = 5 sections; 1,269 drd2^+^ neurons), NAcC (blue; n = 5 sections; 1,055 drd2^+^ neurons), or NacSh (purple; n = 5 sections; 1,114 cre^+^ neurons). Solid bars denote mean and s.e.m. throughout. (**j**) Example fluorescent confocal microscopy image of a coronal section from a DR2-Cre mouse that underwent fluorescent *in situ* hybridization with probes targeting drd1a and drd2 receptor mRNA. *Left*: 20x magnification tilescan spanning dorsal and ventral striatum. *Right top*: 63x confocal images of dorsomedial striatum (DMS, red square) and expression of drd1a mRNA (green), drd2 mRNA (red), and merged image of both (yellow). White triangles indicate co-expression of receptor probes in single neurons. *Right middle*: same as *right top* but for 63x confocal images of nucleus accumbens core (NAcC, blue square). *Right bottom*: same as *right top* but for 63x confocal images of nucleus accumbens shell (NAcSh, purple square). (**k**) Percentage of drd2^+^ neurons co-expressing drd1a mRNA from 2 D2R-Cre and 2 D1R-tdTomato mice in the DMS (red; n = 10 sections; 2,423 drd2^+^ neurons), NAcC (blue; n = 10 sections; 2,196 drd2^+^ neurons), or NacSh (purple; n = 10 sections; 2,220 drd2^+^ neurons). Circles indicate mean overlap from individual sections. (**l**) Same as **k**, but for percentage of drd1a^+^ neurons co-expressing drd2 mRNA from 2 D1R-tdTomato mice in the DMS (red; n = 5 sections; 868 drd1a^+^ neurons), NAcC (blue; n = 5 sections; 834 drd1a^+^ neurons), or NacSh (purple; n = 5 sections; 874 drd1a^+^ neurons). Throughout solid bars reflect mean +/- S.E.M. and transparent ‘o’ denote individual slice mean. Each staining was repeated on 2 independent samples (mice) per group with similar results.

**Supplementary Fig. 2.**
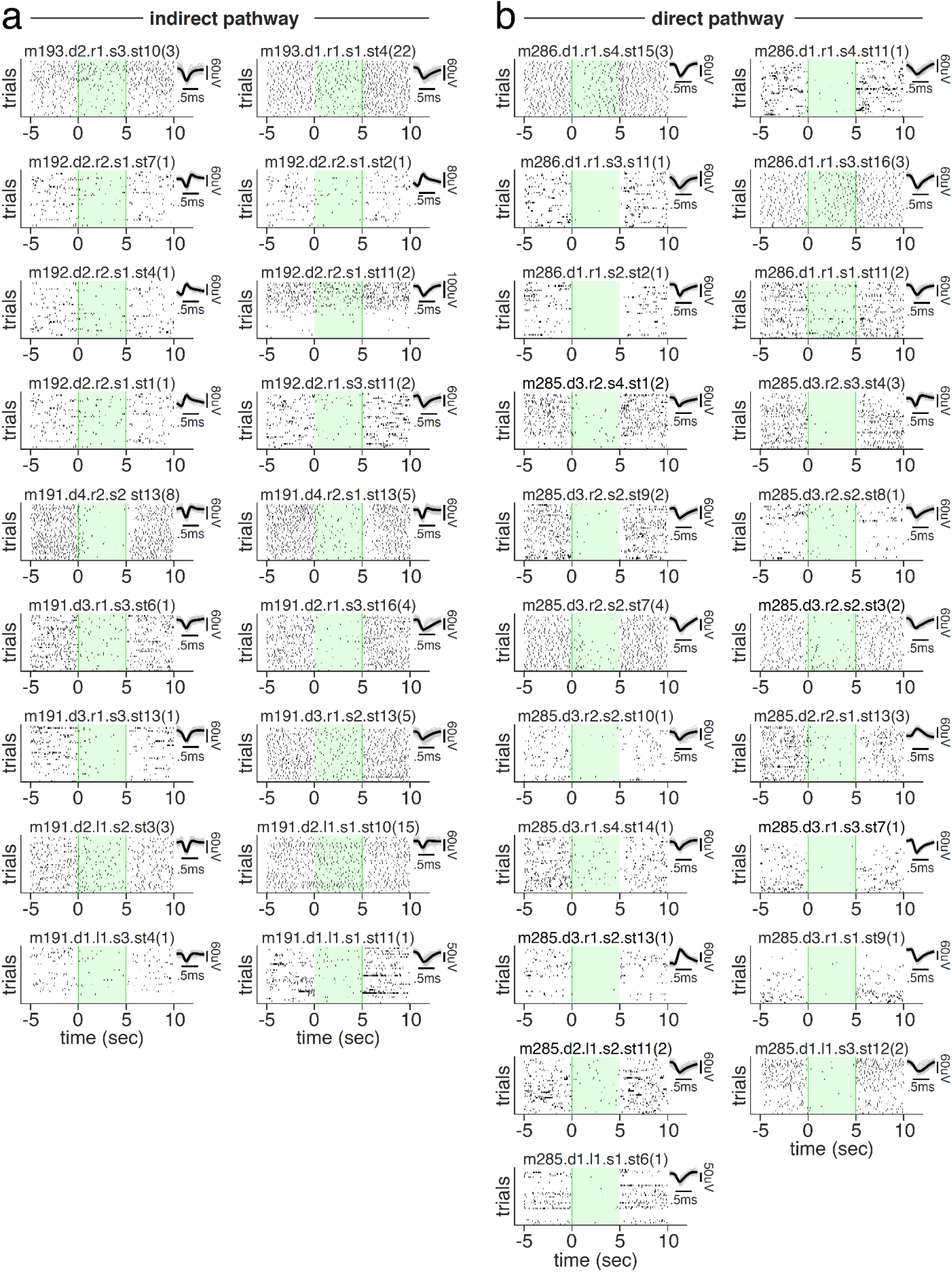
Indirect and direct pathway inhibition of the DMS is stable across time. (**a**) Trial-by-trial raster plots of single neuron spiking during laser off baseline (−5 to 0s), 532-nm (5-mW) laser delivery (0 to 5s), and post laser offset (5 to 10s) for all significantly inhibited neurons (n = 18/60) recorded from A2a-Cre mice expressing Cre-dependent NpHR in the DMS. 40 total trials of laser sweeps per recording site (~15 minutes), ordered in time top to bottom. Individual neuron labels indicate: m (mouse), d (day of recording), r/l (right/left hemisphere and penetration number), s (site or depth of recording probe numbered ventral to dorsal), and st (probe stereotrode channel). Number in parenthesis indicates the number of spikes sub-sampled for display. Inset displays average (bold) and 100 randomly sampled spike waveforms (grey). (**b**) As in **a** but for all significantly inhibited neurons recorded from D1R-Cre mice expressing Cre-dependent NpHR in the DMS (n =21/50).

**Supplementary Fig. 3.**
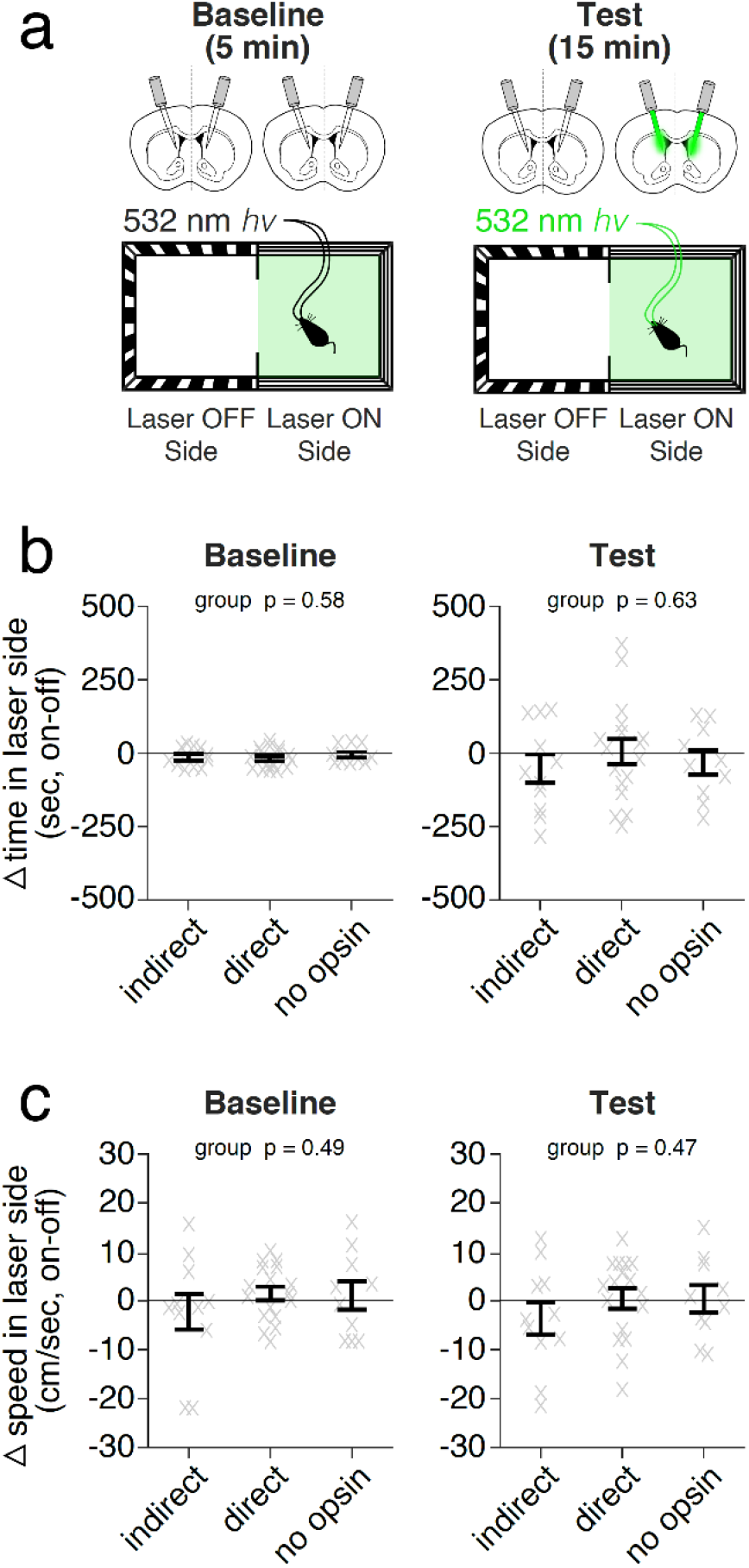
No detectable effect of indirect or direct pathway inhibition on spatial preference and speed during a real-time conditioned place preference test. (**a**) Schematic of real-time conditioned place preference chamber and bilateral 532-nm laser illumination (5-mW) of the DMS. Left and right sub-chambers of equal size but with repeating vertical or horizontal black-and-white bar patterning distinguished each side, respectively. Mice underwent a 5-min preference test (*left*, Baseline) without any laser illumination, followed by a 20-min preference test (*right*, Test) in which mice received bilateral laser illumination only when occupying one of the two chamber sides (counterbalanced across mice). No illumination (Laser OFF) and illumination (Laser ON) sides during Baseline were defined based on the subsequent Test illumination side. (**b**) Delta time spent in chamber side (laser OFF - laser ON) during 5-min Baseline (*left*) and 20-min Test (*right*) for mice receiving DMS indirect (n = 9 mice) or direct (n = 20 mice) pathway inhibition, or DMS illumination alone (no opsin, n = 9 mice). Error bars denote mean and s.e.m. Grey transparent ‘x’ indicates individual mice. p-value denotes one-way ANOVA of group on delta time in the chamber side during Baseline (*left*: p = 0.73, F_2,27_ = 0.31) or Test (*right*: p = 0.10, F_2,27_ = 2.55). (**c**) Average speed when mice occupied laser off (black) or laser on (green) chamber sides during Baseline (*left*) or Test (*right*) for same groups and order as in **b**. Solid bars indicate mean and s.e.m. Transparent grey lines indicate individual mouse mean. p-value denotes interaction of two-factor (between-subject: group, within-subject: laser) repeated measure ANOVA on speed during Baseline (*left*: group x laser interaction: p = 0.16, F_1,35_ = 2.07; laser: p = 0.28, F_1,35_ = 1.20) or Test (*right*: group x laser interaction: p = 0.07, F_1,35_ = 3.6; laser: p = 0.10, F_1,35_ = 2.8).

**Supplementary Fig. 4.**
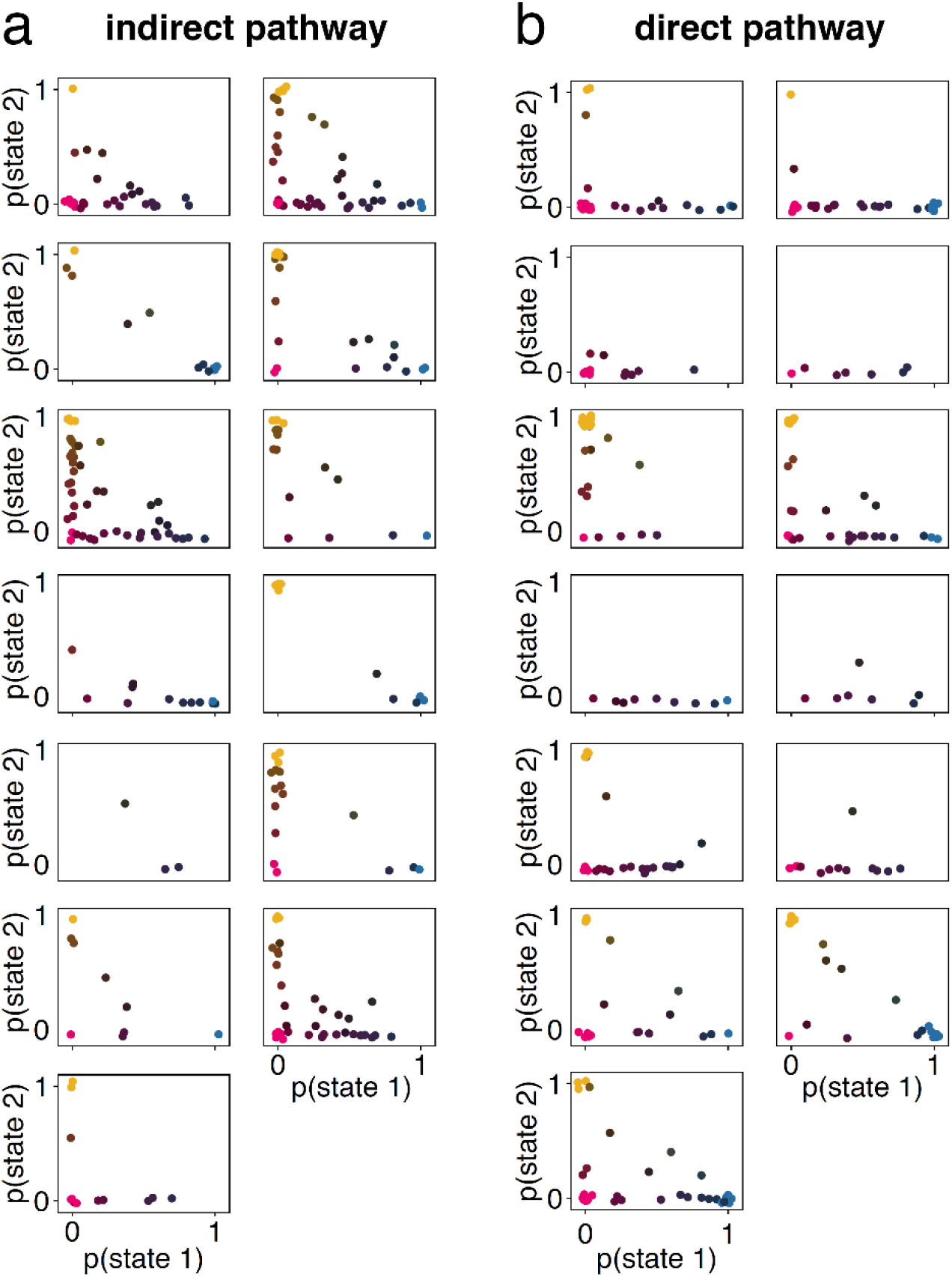
Individual mice visit multiple types and numbers of states over the course of sessions. (**A**) The fraction of trials that mice inhibited in the indirect pathway of the DMS spent in each state in each session. Each box represents a different mouse (n=13) and each dot in each box represents an individual session for that mouse. Color-coding reinforces the state composition of each session (e.g. blue indicates the mouse spent 100% of the session in state 1). A small amount of Gaussian noise was added to the position of each dot for visualization purposes. (**B**) Same as **A** but for mice inhibited in the direct pathway of the DMS (n=13).

**Supplementary Fig. 5.**
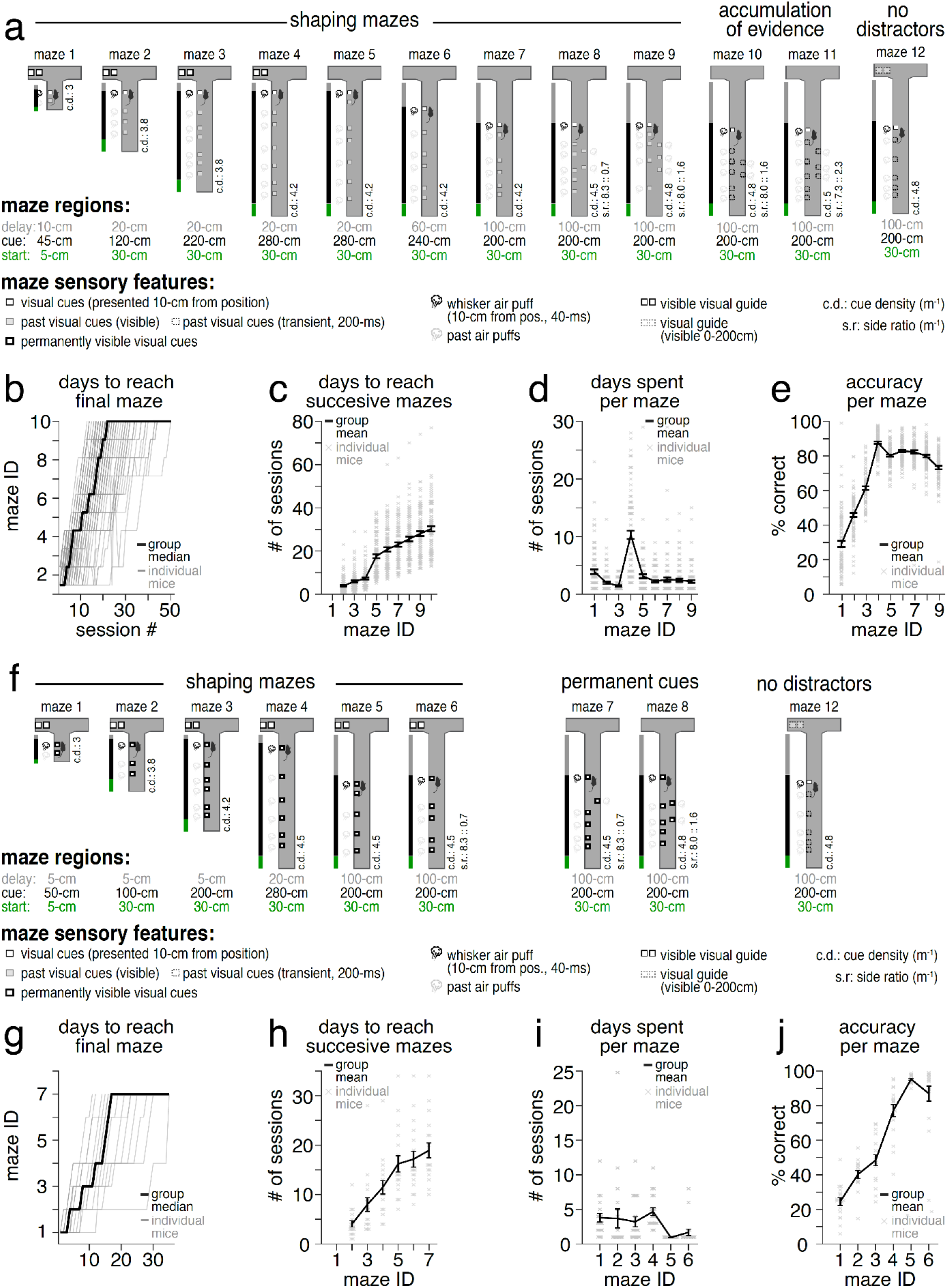
Behavioral shaping for virtual-reality T-maze tasks. (**a**) Schema of shaping mazes (*top*: maze 1-9) and subsequent optogenetic testing mazes (*far right*) for the accumulation of evidence task. Mazes varied according to the following sensory features: the length of start, cue, and delay regions (green, black, and grey bars and colored text, respectively), whether visual cues were presented 10-cm from cue position (black outline, white filled square) and remained visible (grey square) or disappeared 200-ms after presentation (black dotted, unfilled square) or were permanently available from trial outset (bold black border, white filled), the presence of left or right whisker air puffs (15-psi, 40-ms) which were delivered upon first instance of being 10-cm from visual cue position (solid vs grey puff symbol), whether a visual guide was located in the rewarded arm (black double square) or if the visual guide was only visible during the cue region (grey double square), the density of cues during the cue region (c.d.), and whether distractor cues occurred on the non-rewarded maze side (side ratio, s.r.: mean density per meter). Following shaping mazes 1-9, optogenetic testing was carried out on mazes 10 and 11 (accumulation of evidence), and maze 12 (no distractors). (**b**) Solid black line depicts the across-mouse median number of sessions spent on each shaping maze (mazes 1-9) until reaching the first testing maze (maze 10) (group median: 22 sessions). Grey transparent lines depict the median number of sessions for individual mice (n = 79). (**c**) Cumulative number of sessions to reach each successive maze until the first testing maze (maze 10) (group mean: 23.0 +/- 0.8 sessions). Solid black lines depict mean and s.e.m. across mice, and transparent ‘x’ denote individual mice. (**d**) Total number of sessions spent on each shaping maze. Solid black lines depict mean and s.e.m., and transparent ‘x’ denote individual mice at each respective shaping maze (maze 1-9). (**e**) Percent correct performance across shaping mazes. Solid black lines depict mean and s.e.m., and transparent ‘x’ denote individual mice at each respective shaping maze (maze 1-9). (**f**) Same as **a** but for shaping (*left*, maze 1-6) for the permanent cues task. Following shaping mazes 1-6, optogenetic testing was carried out on mazes 7 and 8 (permanent cues) and maze 12 (no distractors). (**g**) Same as **b** but for permanent cues shaping (group median: 17 days; n = 20 mice). (**h**) Same as **c** but for permanent cues shaping (group mean: 18.9 +/- 1.5 sessions). (**i**) Same as **d** but for permanent cues shaping. (**j**) Same as **e** but for permanent cues shaping.

**Supplementary Fig. 6.**
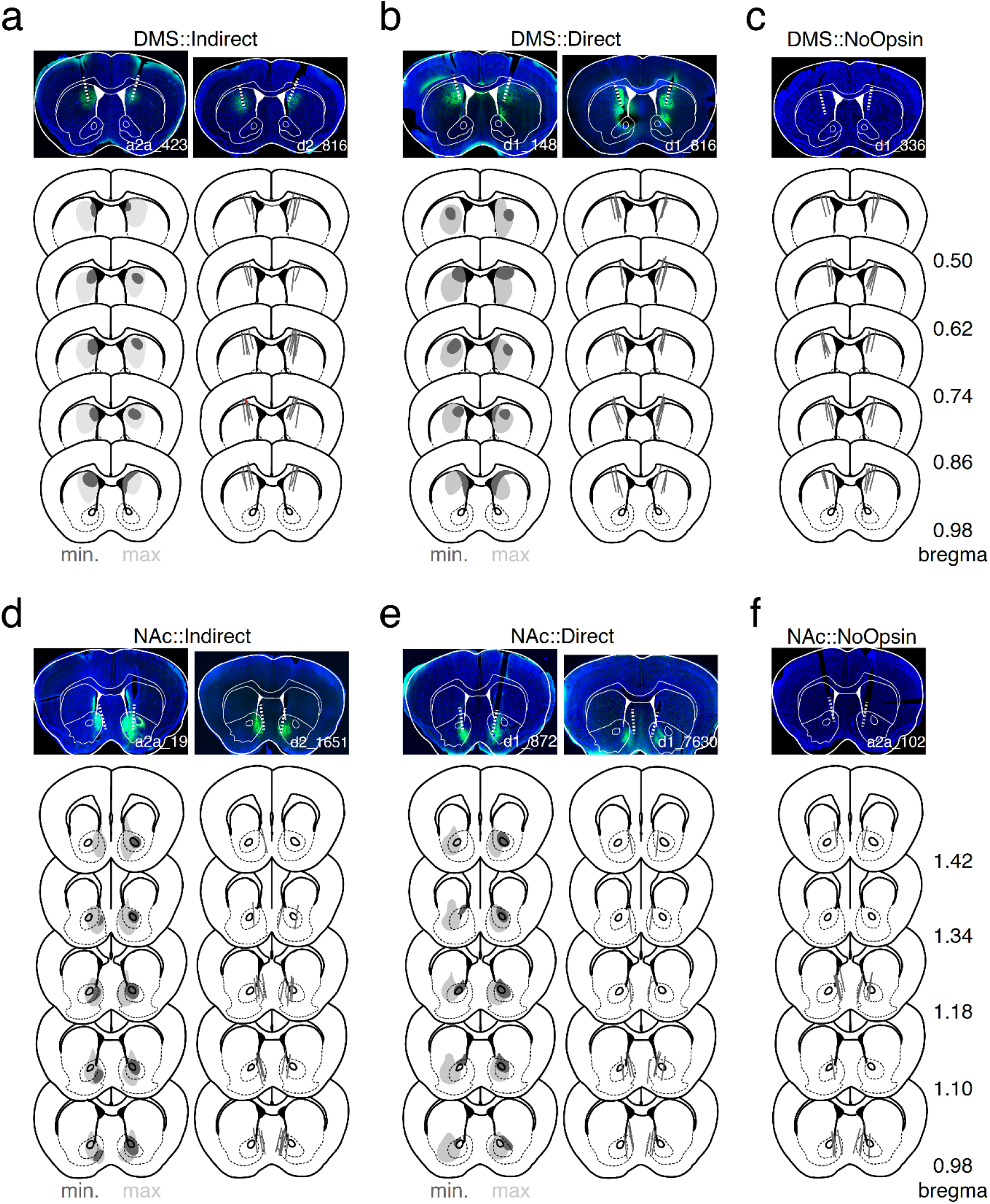
Histological confirmation of viral expression and fiberoptic placement. (**A**) *Top*: two individual mouse examples of cre-dependent NpHR-GFP expression in and fiberoptic targeting of the dorsomedial striatum (DMS) in D2R-Cre or A2a-Cre mice. *Bottom left*: schematic representation of the minimum (dark grey) and maximum (light grey) spread of NpHR expression in all mice targeting the indirect pathway of the DMS (DMS::Indirect, n = 21 mice). *Bottom right*: summary of tapered fiberoptic tip location and angled track for all mice targeting the indirect pathway of the DMS. (**B**) Same as **A** but for all experiments targeting the direct pathway of the DMS (DMS::Direct, n = 23 mice). (**C**) Same as **A** but for fiberoptic targeting only for all control mice receiving DMS illumination in the absence of NpHR (DMS::NoOpsin, n = 17 mice). (**D**) Same as **A** but for experiments targeting the indirect pathway of the nucleus accumbens (NAc::Indirect, n = 9 mice). (**E**) Same as **A** but for experiments targeting the direct pathway of the nucleus accumbens (NAc::Direct, n = 10 mice). (**F**) Same as **C** but for all control mice receiving NAc illumination in the absence of NpHR (NAc::NoOpsin, n = 7).

**Supplementary Table 1.**
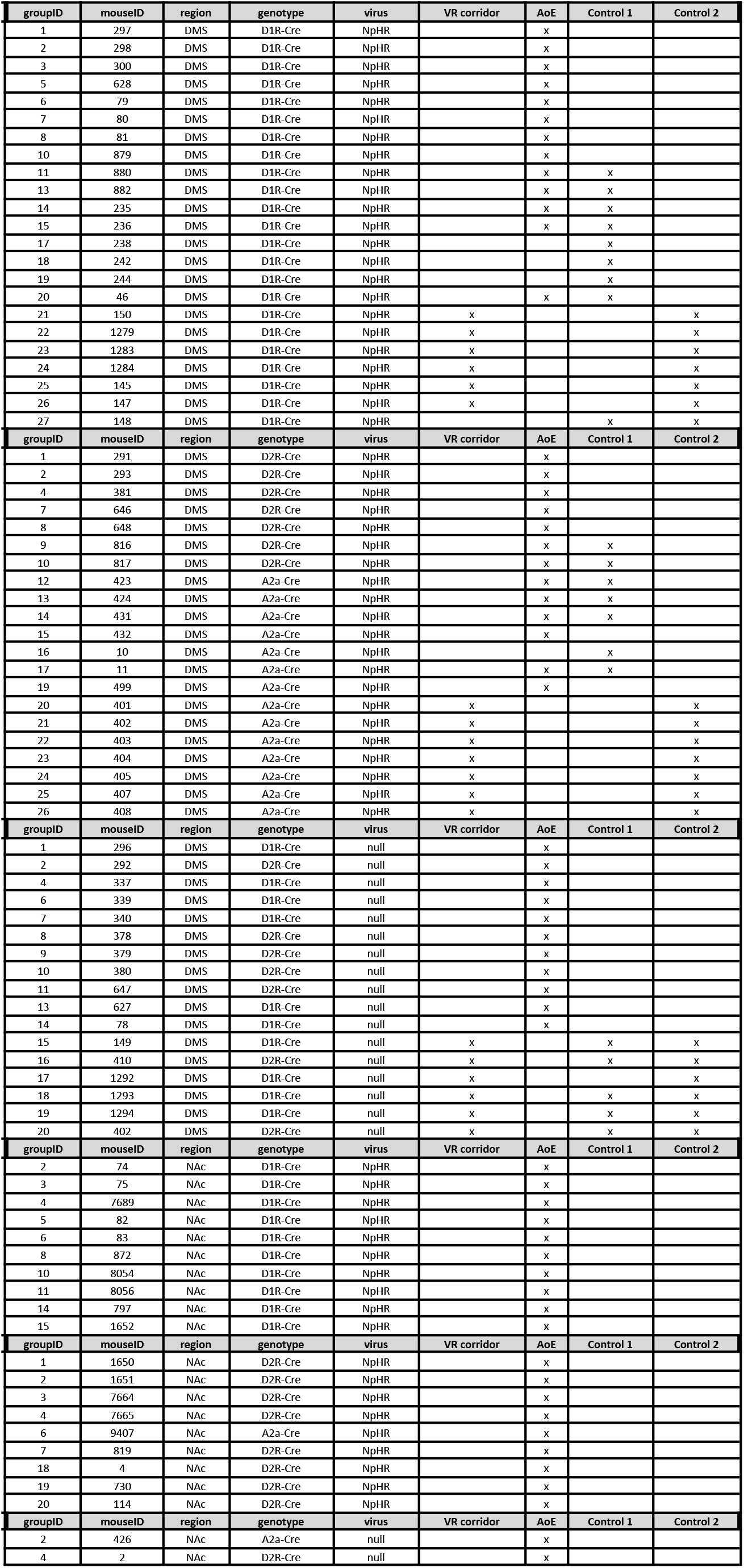
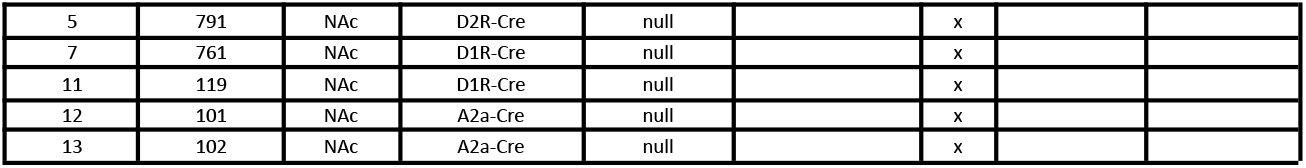
Summary of individual mice included in data sets. GroupID reflects the unique identifier of individual mice within a cohort. MouseID reflects unique mouse identifier. Region reflects target of bilateral fiberoptic implants (DMS or NAc). Genotype reflects transgenic background of mice (A2a-, D2R-, or D1R-Cre). Virus reflects delivery of AAV5-DIO-NpHR-EYFP (NpHR) to fiberoptic targeted structure or absence of opsin (null). In columns VR corridor, AoE (accumulation of evidence task), control 1 (no distractors task), and control 2 (permanent cues task), an ‘x’ indicates a mouse underwent optogenetic testing in this maze and was included in subsequent analyses. Note that trial selection criterion (see **Methods**) provided additional thresholds for inclusion in analyses.

